# Labels as a Feature: Network Homophily for Systematically Discovering human GPCR Drug-Target Interactions

**DOI:** 10.1101/2024.03.29.586957

**Authors:** Frederik G. Hansson, Niklas Gesmar Madsen, Lea G. Hansen, Tadas Jakočiūnas, Bettina Lengger, Jay D. Keasling, Michael K. Jensen, Carlos G. Acevedo-Rocha, Emil D. Jensen

## Abstract

Machine learning (ML) has revolutionized drug discovery by enabling the exploration of vast, uncharted chemical spaces essential for discovering novel patentable drugs. Despite the critical role of human G protein-coupled receptors (hGPCRs) in FDA-approved drugs, exhaustive in-distribution drug-target interaction (DTI) testing across all pairs of hGPCRs and known drugs is rare due to significant economic and technical challenges. This often leaves off-target effects unexplored, which poses a considerable risk to drug safety. In contrast to the traditional focus on out-of-distribution (OOD) exploration (drug discovery), we introduce a neighborhood-to-prediction model termed Chemical Space Neural Networks (CSNN) that leverages network homophily and training-free graph neural networks (GNNs) with Labels as Features (LaF). We show that CSNN’s ability to make accurate predictions strongly correlates with network homophily. Thus, LaFs strongly increase a ML model’s capacity to enhance in-distribution prediction accuracy, which we show by integrating labeled data during inference. We validate these advancements in a high-throughput yeast biosensing system (3773 DTIs, 539 compounds, 7 hGPCRs) to discover novel DTIs for FDA-approved drugs and to expand the general understanding of how to build reliable predictors to guide experimental verification.

## 1 Introduction

Machine learning (ML) methods have significantly advanced high-throughput drug discovery pipelines by enabling virtual compound screening for large chemical spaces, the vastness of which are inaccessible to fully explore experimentally. Furthermore, public drug-target interaction (DTI) databases are growing at an exponential rate, hence enabling the further enhancement of ML methods at predicting compound bioactivity *(1, 2)*. Human G protein-coupled receptors (hGPCRs) are one of the most important classes of drug targets and include over 286 non-olfactory receptors *(3)*, which are targeted by approximately 35% of all FDA-approved pharmaceuticals *(4)*.

Despite the critical role of hGPCRs in drug discovery and their documented promiscuity *(3, 5)*, it remains uncommon to exhaustively test all hGPCRs against all drugs in a library *(6)*. Illustratively, there are a lot of empty *in-distribution* labels missing (Figure 1a). For instance, for 186 thousand unique hGPCR-targeting compounds across 128 hGPCRs, only 1.5 % of all possible activities have been documented in publicly sourced data. Thus, 98.5 % of DTIs remain in the dark. Naturally, this is largely due to the significant economic and technical challenges involved at this scale. Nevertheless, most approved hGPCR-targeting drugs are accompanied by famously extensive medicine package leaflets documenting the incidence rate of side-effects, which are likely due to off-target drug-interaction(s) *(7, 8)*.

**Figure 1:**
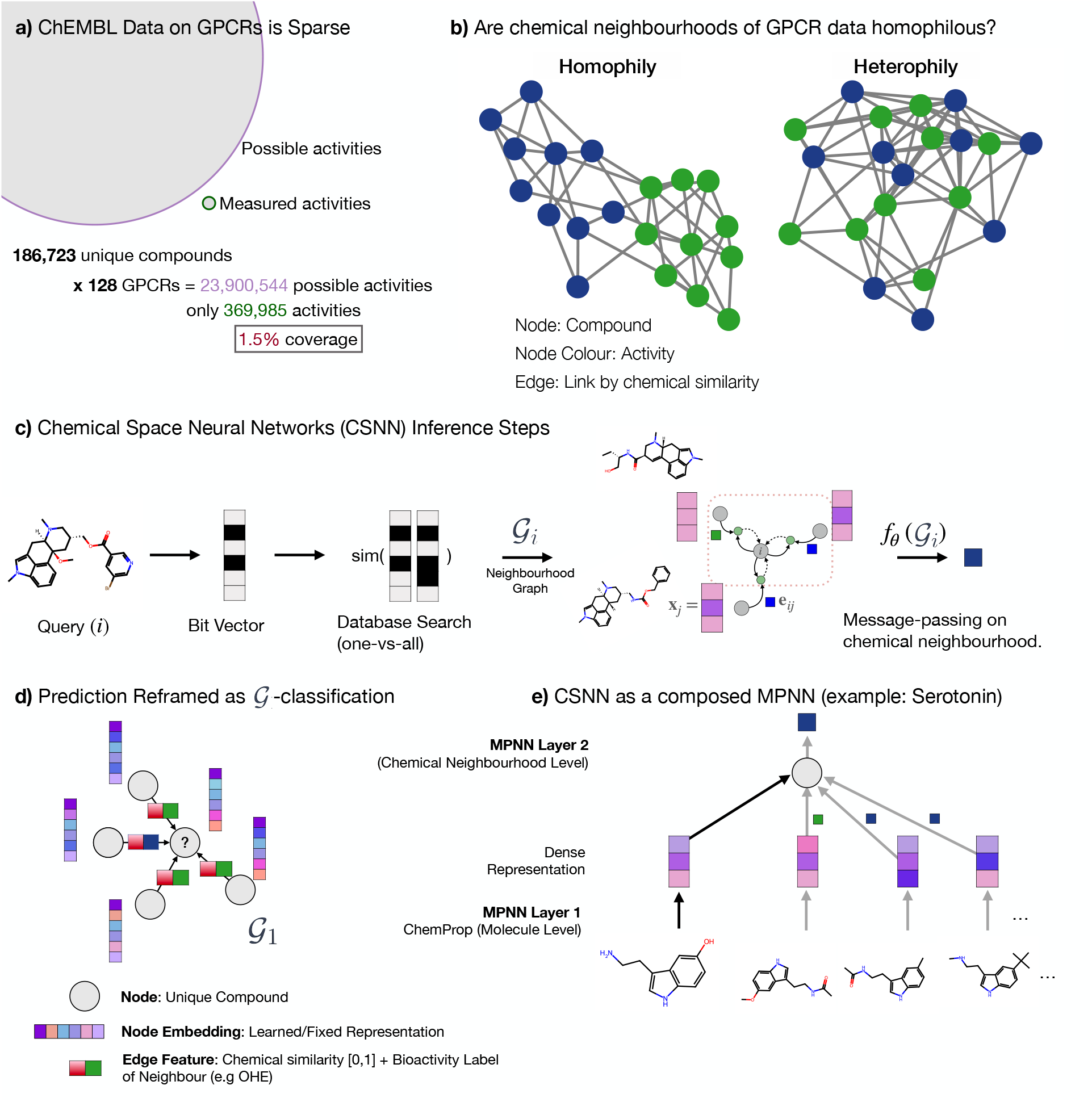
Introducing Network Homophily and Transductive Node Classification. **(a)** Collected data on bioactivity classes for 186 K unique hGPCR-targeting compounds across 128 hGPCRs only has 369 K of 23.9 M possible activities. This indicates the sparse annotation in public databases. **(b)** A visual example to introduce network homophily: “similarity breeds connection” *(10)*. **(c)** Inference for CSNN to illustrate how labels as a feature are used. First the query compound is encoded by a bit-vector, then a one-vs-all database search returns compounds, which are chemically similar and labels on those compounds. A neighbourhood graph 𝒢_*i*_ is constructed that is fed to the ML network *(f*_*θ*_). **(d)** Instead of transductive node classification, we simplify the task to transductive graph classification by introducing a one-hop directed graph (incident on the query) with neighbourhood LaFs as edge attributes. **(e)** The CSNN framework can be viewed a as composed message-passing neural network (MPNN): First at the molecule level (MPNN on atoms) and then again at the neighbourhood level including LaFs.

Solving this problem by exhaustively documenting off-target effects is thus the opposite of drug discovery pipelines: These slowly move toward undocumented regions of chemical space (out-of-distribution (OOD) and patentability). However, in this case, robust and reliable *in-distribution* prediction is required to narrow down the DTIs selected for experimental verification.

A critical inductive bias in many ML models is the assumption of smoothness or locality, where there is a presumed continuity or proximity between the input space and the output space. This implies proximity between similar compounds in chemical space. In chemoinformatics, this is known as the chemical homophily principle *(9)* and is more generally known as network homophily. Intuitively, “similarity breeds connection” and is found in a range of social, economic behavior as well as network based databases *(10)*. By analogy, similar molecules have similar bioactivity and, if the network is homophilous, prediction is straight-forward when a chemical neighbourhood is known (Figure 1b).

Yet, ML for drug discovery has primarily focused on compound-to-prediction architectures rather than considering the chemical data available in the chemical neighbourhood during inference (neighbourhood-to-prediction) (See Table S10 for a comparison). If indeed DTI data is homophilous for hGPCRs, then it would be beneficial to include data during inference to increase reliability and robustness of predictions.

An emerging paradigm known as training free graph neural networks (TFGNNs) *(11)*, show how labels as a features (LaF) is an admissible operation in transductive node classification tasks: labels of neighbouring nodes are used to update the learned representations. The paradigm proves how this approach increases the expressive power over Graph Neural Networks (GNNs). Empirically, the results show that even without training, prediction accuracy is incredibly high on common benchmark GNN datasets. Here, we demonstrate similar training free predictions that are on-par or outperform conventional trained compound-to-prediction architectures. Instead of framing the task as transductive node classification, we re-frame it as a graph classification task with neighbourhood labels as edge features (Figure 1d) in a directed (transductive) graph, which integrates available data during inference. We demonstrate strong network homophily in hGPCR bioactivity data and show a strong correlation between network homophily and machine learned prediction accuracy. By exploiting the graph homophily of chemical space networks (CSN) we develop Chemical Space Neural Networks (CSNN) and combine the method with a high-throughput yeast biosensing system for experimentally validating novel-DTIs. Finally, we correlate predictions with experimental results and discover 14 novel DTIs potentially linked with off-target effects.

## 2 Results

### Building the architecture of Chemical Space Neural Networks

As a basis for our work, we compiled a dataset containing cleaned bioactivity data on hGPCRs available in ChEMBL*(12)* and the IUPHAR/BPS guide to pharmacology*(13)*. In total, the dataset comprises 186,723 unique molecules represented as SMILES and labeled with known bioactivity classes against the 128 respective hGPCRs (Supplementary File S1).

Figure 1a shows that ChEMBL has >369,000 annotated activities across 128 hGPCRs. This is, however, a vanishing fraction of all possible interactions (1.5 % coverage), leaving knowledge of the majority (98.5 %) of DTIs in the dark. ChEMBL is growing at a rate of 780 K activities per year (Supplementary Figure S17), which illustrates the need for constructing reliable predictive algorithms to fill in the missing interactions.

Given the homophily principle assumption, compounds which are close in chemical space often have similar bioactivity class on a given hGPCR, it is thus conceivable that a chemical neighbourhood is strongly indicative of a compound’s bioactivity class. This is illustrated in Figure 1b, d, where we extended upon works by Yu-Chen Lo et. al *(14)* and Zengrui Wu et. al *(6, 15)* to develop Chemical Space Neural Networks (CSNN). The key advantage is illustrated in Figure S1: Commonly, methods using ML for hGPCR drug-discovery are compound-to-prediction architectures. This takes a compound from the chemical space and directly infers its target through some learned or fixed representation (e.g., chemical fingerprints) with ML models. CSNN, developed in this work, takes a local chemical neighbourhood by querying a database during inference, learning a local representation using LaF, and then using the constructed neighbourhood graph for predicting DTIs (regression and classification).

Figure 1c illustrates the inference process of CSNN, which operates akin to the transductive node classification task. First, a query compound *i* is represented by a bit-vector, then a one-vs-all database search (by Tanimoto similarty) returns chemically similar neighbours (related compounds). From a pre-trained directed message-passing neural network (D-MPNN, ChemProp) a dense representation is obtained for the query and neighbouring compounds and edge features are compiled to a neighbourhood graph 𝒢_*i*_ as illustrated in Figure 1d. Here, neighbouring compound labels are used as edge features. Then a trained MPNN takes the graph 𝒢_*i*_ mapping it to it’s y-value (classification/regression): *f*_*θ*_ : 𝒢_*i*_ → **y**_*i*_. Effectively, the method is composing MPNN layers at the level of compounds (MPNN on atoms) and again at the neighbourhood level (MPNN between DTIs) as shown in Figure 1e. To overcome the current limitations of CSNs, we develop an extremely fast way of constructing the chemical neighbours by reducing the computation of the 186 K x 186 K adjacancy matrix (CSN) to 20 min (described in methods and additional results in S8). Inference for a one-vs-all (1-vs-186 K) lookup takes 0.187 ± 0.002 seconds.

### Chemical Neighbourhoods enable training frees and confident predictions of hGPCR bioactivity classes

With an established all-vs-all CSN (Fig. 2a) from which one can derive important neighbourhood statistics, we can from the visualised example presented in Figure 2b see that the the query molecule has primarily ‘Agonists’ in its neighbourhood, which is in fact the true label of the compound for the hGPCRs shown. More generally, across 201,120 unique DTIs and across all 128 hGPCRs and 6 bioactivity classes, the neighbourhood achieved an F1 score of up to 93 % and shows remarkable performance across bioactivity classes (See Figure 2c and S3). Here we simply calculate the frequency of each label in the neighbourhood of a query and take the most frequent label as the prediction (Argmax operation), then compare with the true label on a given hGPCR. Thus the neighbourhood labels becomes a feature for inference. This demonstrates the strong homophily of the hGPCR-DTI space and shows a training free GNN approach with LaFs. To enable ML on these neighbourhoods, we structure the data into the standardised open-source PyTorch Geometric framework *(16)* (Figure 2d).

**Figure 2:**
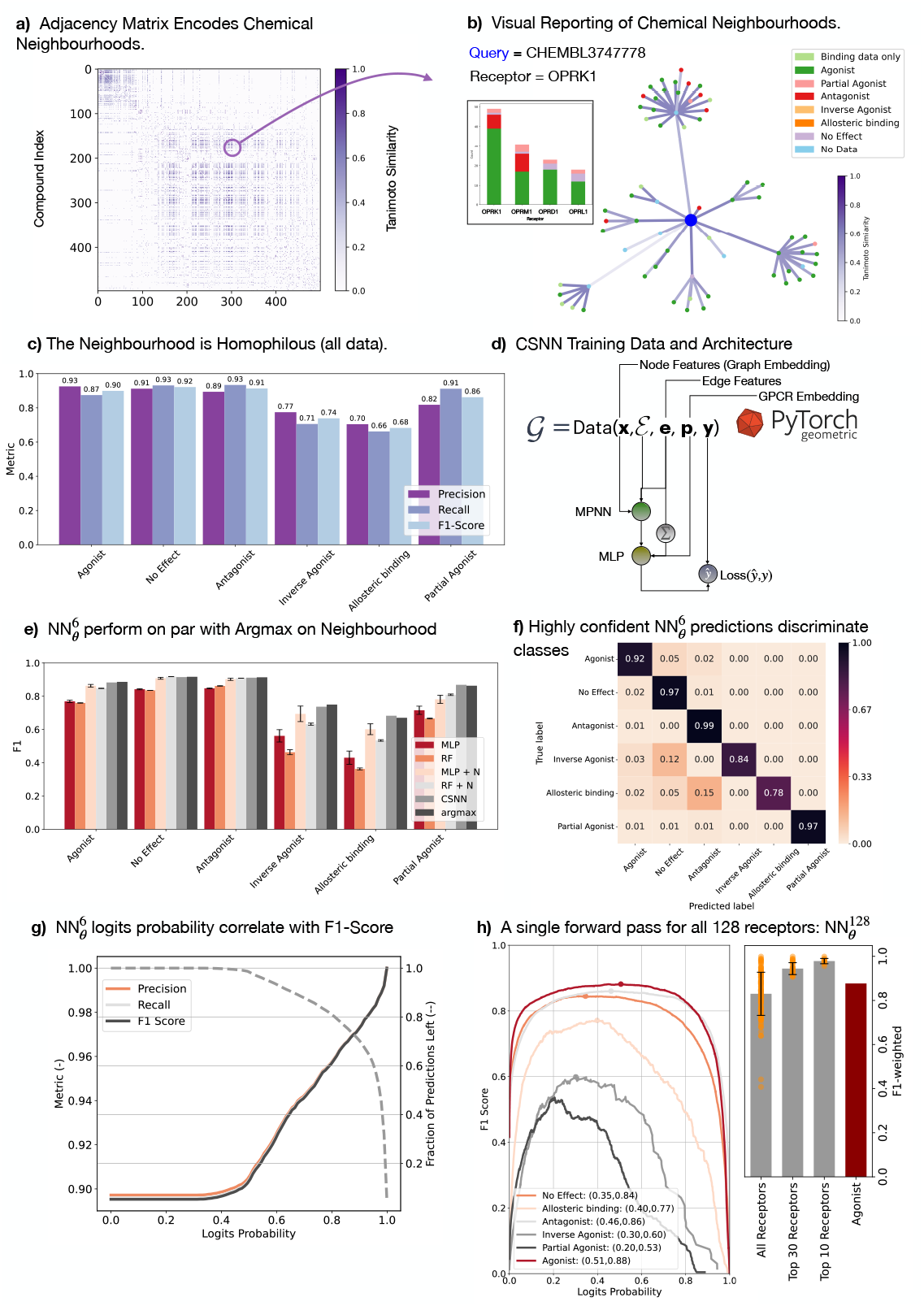
Benchmarking Bioactivity Label Prediction with LaFs. **(a)** The adjacancy matrix (186 K × 186 K possible connections) parameterised the all-vs-all chemical space neighbourhood. **(b)** Querying the CSN can return data on related compounds already illustrating graph homophily (agonist are over represented and the true label for OPRK1 is agonist). **(c)** Training free prediction metrics using the most frequent LaF in the neighbourhood, demonstrates the strong network homophily. **(d)** Structuring the dataset into an accessible format and CSNN architecture. **(e)** Prediction metrics on test set for ML methods: without LaFs (MLP, RF), with LaFs (MLP + N, CSNN), and a full MPNN on chemical neighbourhoods (CSNN) compared to the training free prediction (Argmax). **(f)** When the class label is confident probability >0.8, the CSNN method (referred to as 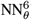) produces high-quality class discrimination. **(g)** As 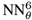 class label is filtered by logits probability, the performance metrics tend toward perfect predictions. **(h)** 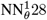 CSNN model (one forward pass for class labels across all 128 hGPCRs using LaFs) shows strong performance metrics for most hGPCRs.

In Figure 2e, f and g we illustrate the performance metrics of ML methods that operate on chemical neighbourhoods contrasted with normal compound-to-prediction methods (without a neighbourhood of LaFs). In Figure 2e we compare common models (Multilayer Perceptron: MLP, and Random Forest: RF) without the neighbourhood using only the compound representation and with the neighbourhood (+ N) frequency vector (LaF concatenated). Just this LaF inclusion significantly improves performance. Interestingly, the developed CSNN method (GNN on chemical neighbourhood) is only on par with the Argmax prediction. We refer to this method hence-forth as 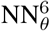 as it produces logits for the six bioactivity classes.

However, if one only includes highly-confident predictions (logits are strongly peaked, high class probability), the performance outperforms Argmax classification metrics (Fig. 2f,g). This falls in line with the intuition provided in *(11)*, that LaFs improve the expressive power of GNNs. At a high class probability (> 0.8) the classification is almost without error on the test set (see Fig. S2a for complete class summaries and Fig. S2b,c for ROC-AUC curves). The advantage of neural network over Argmax predictions is that the former are probabilistic and that the model confidence directly correlates with the mean ROC-AUC (Fig. S5). For further and more complete investigations of the 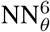 model, see supplementary text and Figures S2 -S5.

To generalise the transductive node classification method to include all data on all hGPCRs in the neighbourhood and reduce the number of forward passes to predict labels for all compound-hGPCR pairs, we develop a 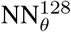 CSNN model, which in a single forward pass produces bioactivity class logits for all 128 hGPCRs at once. The CSNN takes a graph 𝒢 as input and predicts bioactivity as **y** ∈ ℝ^1*×*128*×*6^, which represents class-wise logits. The input graph contains all available data on all 128 hGPCRs as edge features (LaF), thus allowing for mixing between data channels. This was intuitively advantageous, as related hGPCRs show similar signalling responses to the same molecule, thus enhancing the predictive performance by integrating all available data in the sparsely label dataset. Results of this method are reported in Figure 2h, which shows that for the top-10 scoring hGPCRs, classification was near perfect (class weighted F1-score of 0.978 ± 0.012). Across all hGPCRs, the classification weighted F1-score was 0.83 ± 0.10, whereas if only the agonist class assignment is considered, the F1-score was 0.877. Further quality metrics and full class summaries are provided in Figure S6a,b,c, where we demonstrate a strong correlation between testing F1-score and training support for each bioactivity class and ROC-AUC > 0.95 for all classes, thus outperforming the 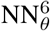 model.

### Benchmarking Neighbourhood Methods on Regression Tasks and newly generated datasets

As LaF showed strong promise in predicting class labels, we perform a benchmark study on a recently published method and dataset (pdCSM *(17)*) in the more challenging regression setting for binding affinity predictions. We compute a CSN on all compounds and allow training nodes to refer to each other, while restricting testing compounds only to refer to the training data during inference (admissible transductive framework). Figure 3a, shows a schematic training example with three nodes (compounds) and their true − log_10_*(K*_*i*_) values. Taking a mean over the neighbourhood (6.20) is an acceptable prediction of the true regression label (6.14). Figure 3b shows that the mean prediction *< Y*_*neigh*_ *>* is accurate across all hGPCRs in the dataset (MSE = 0.685, MAE = 0.624, Pearson = 0.757). Regression plots split by hGPCR are found in Figure S9 and as a function of aggregation technique (mean, max, min, weighted) in Figure S10.

**Figure 3:**
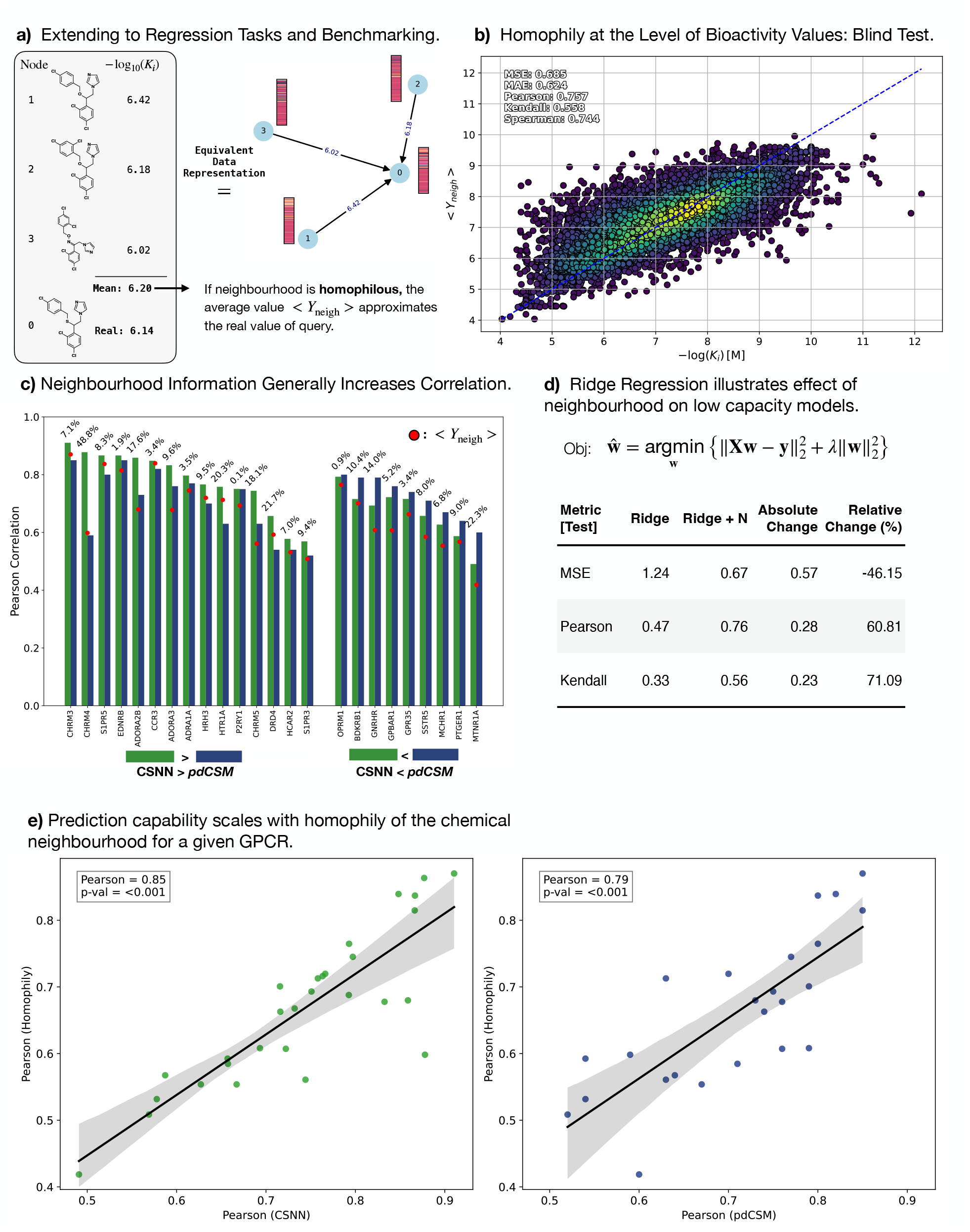
Benchmarking LaFs in the Regression Setting. **(a)** An illustrated training example: node 0 is the query, nodes 1-3 are neighbouring compounds and their bioactivity label (LaF). The mean over the neighbourhood is 6.20, which is close to the true label of 6.14 (units: − log_10_*(K*_*i*_)). **(b)** Training free mean value *(< Y*_*neigh*_ *>)* predictions on the test set using LaFs across all hGPCRs in the pdCSM dataset shows strong performance metrics. **(c)** Comparing the published pdCSM method (RF on RDKit compound representations) with LaFs (CSNN) on the same compound representations. In most cases LaFs improve the Pearson correlation coefficient. The *< Y*_*neigh*_ *>* prediction closely track the top performing method. **(d)** Illustrating the effect of a low-capacity Ridge regression model without (column: Ridge) and with LaFs (column: Ridge + N). The input dimension differs only by one column (the mean value over neighbourhood). **(e)** Interestingly, the test metrics on a given hGPCR correlates strongly with the Pearson correlation under the homphily assumption (mean value is a good prediction). The LaF methods outperform none LaF methods (compound-to-prediction architectures).

To benchmark our method, we reconstruct molecule representations used for pdCSM with available data from *(17)* (see dataset summary in Table S9, in which training/test data points 47013/7114 were used based on the published data). We chose pdCSM due to the relative dataset size and published source data. We instantiate a CSNN using a random forest (RF) model as done in the published benchmark method (pdCSM), but include the neighbourhood mean value prediction, which allows a comparison between methods without and with neighbourhood information (LaF). Unlike the pdCSM method, which has no available training or inference code, we do not perform feature selection as their code was not available to replicate the method. Nevertheless, CSNN outperforms pdCSM on 15/24 targets (Fig. 3c, and Figs. S13, S14), and requires only an increase in feature dimension by one (the mean-value of the neighbourhood). Furthermore, CSNN have a guard-rail against OOD predictions, as without a neighbourhood no predictions are made. This is not the case for the benchmark method, which will predict *K*_*i*_ values for *any* SMILES compound. Interestingly, the maximum Pearson correlation achieved on a given hGPCR is highly correlated with the mean-value prediction derived from the homophily assumption (Fig. 3c, e), implying that the more homophilous the neighbourhood (mean-value is a good approximation) is, the better the ML model performs. Accordingly, the pdCSM method only outperforms a training-free prediction *(< Y*_*neigh*_ *>)* on 17/24 hGPCR targets, while CSNN outperforms on all (24/24) hGPCRs. Overall, these results confirm the smoothness inductive bias from the introduction between the input and output space *(f*_*θ*_ : ℝ^*n*^ → ℝ), allowing one to predict the utility of a ML-model depending on the homophily of the dataset.

The effect of having LaFs can be observed in a low capacity model, like Ridge regression as judged by the change in MSE, Pearson correlation and Kendall correlation without and with (+N) the neighbourhood information using the same fixed representation (Fig. 3d). The MSE is reduced by 46 % and the Pearson correlation coefficient increases by 60 %. Again, LaF has a strong effect on reducing the complexity of the learning task, as postulated in *(11)*. The results are less striking but still apparent for RF models with or without LaF, it drives the MSE error down by 7.89 % and increases Pearson correlation by 2 % (Fig. S8) for a fixed model size.

Argmax and CSNN methods are able to expand the total coverage of labeled hGPCR-compound DTIs from 1.5 % to 5.6 % and 100 %, respectively. Naturally, the CSNN method contains a prediction confidence, thus implying that confident predictions do not ensure total coverage. The robust Argmax prediction still left 94.4 % of the DTIs unknown, which can not currently be predicted in a training free manner, as the data is too sparse to impute the missing data. Nevertheless, Argmax easily increased data coverage 4-fold from 370 K to 1.3 million unique drug-hGPCR interactions.

Given that LaFs is a strong, admissible, and a powerful inductive bias of network homophily, which is inherent in the DTI-hGPCR space, we sought to experimentally develop a high-throughput platform to compliment model predictions and to validate both the class probability and regression model.

Heterologous expression of hGPCRs in yeast offers advantage over traditional hGPCR screening methods because it is both cost-efficient and allow for accelerated screening of chemical libraries *(18, 19)* (See Supplementary Notes, Table S5 and supplementary file S2). We therefore constructed a small set of yeast platform strains for 7 hGPCRs previously reported to signal in yeast *(18, 20*–*24)*, which were chosen based on their immediate potentials as drug targets (supplementary table S4). We based our design on the system presented by Shaw et al. (2019) *(25)* yet with NanoLUC *(26)* as a reporter of relative luminescence units (RLU) instead of GFP for increased assay sensitivity *(19)* (See Figure 4a for schematic overview). We confirmed their performances by measuring dose-response curves (DRC) with agonists on each hGPCR (See Figure 4b) to further evaluate the expected quality of gathered data using the yeast platform (Supplementary Figures S20, S21 and table S6). The EC50 values observed for CHRM3 and ADRA2B were comparable to the most sensitive screens reported (Supplementary table S6*(12, 27)*). The dynamic ranges were in 5 out of 7 cases higher in yeast than observed for mammalian cell assays (Supplementary Figure S20, and table S6).

**Figure 4:**
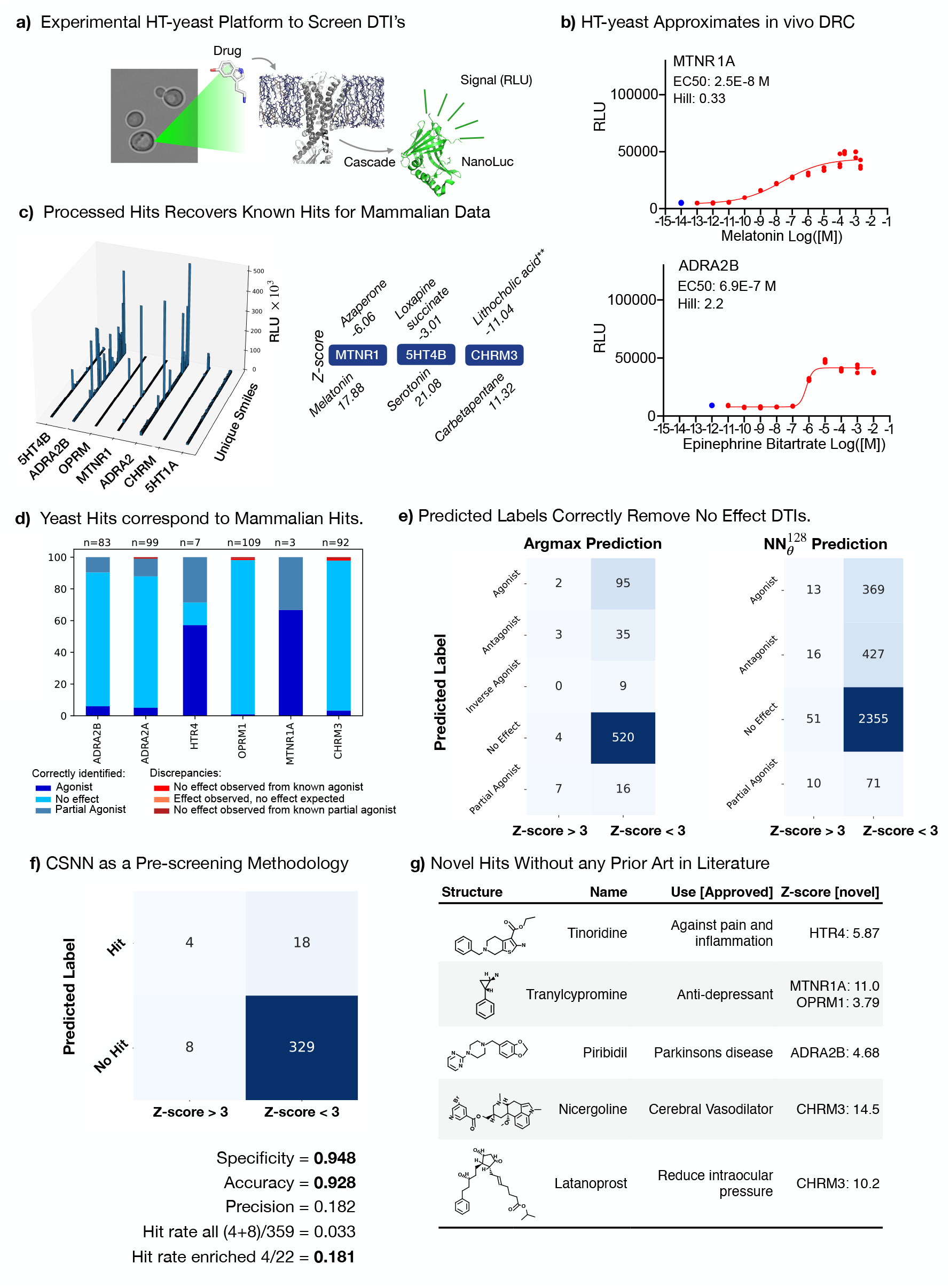
Experimental Validation using a Developed HT-yeast Biosensing Platform. **(a)** Overview of experimental HT-yeast platform based on designs by Shaw et al. 2019 *(25)*. The drug may induce the hGPCR and the signal is measured by a reporter in relative luminescence units (RLU). **b** Dose-response curves (DRCs) reveal large differences in dynamic range and sensitivity (See all hGPCR DRCs in supplementary Figure S20). **(c)** Overview over all-vs-all 3773 DTIs measured (539 compounds and 7 hGPCRs, left) with top hits and their associated Z-score (right), which recovers known agonists. **(d)** Using the CSN as a knowledge graph, we compare the signal response in HT-yeast of DTI with that found in mammalian systems and find a high-correspondance. **(e)** Heatmap of predicted label and the experimental Z-score for DTIs. Argmax predictions and neural network predictions can correctly filter out no effect DTIs. Naturally, the assay cannot capture some of the predicted labels in the Z-score and thus does not correlate with the Z-score bin. **(f)** CSNN can effectively be used to enrich the positive hits (Z-score > 3), with high specificity (0.948) and an enriched hit rate of 18 %. A predicted hit was defined the CSNN prediction for *K*_*i*_ < 100 nM and the label is not ‘No Effect’. The precision is markedly low (0.182), showing difficulty in positive hits. **(g)** Novel hits without prior art in literature. We use the CSN as a knowledge graph, sort for significant hits (|Z-score| > 3), without any database knowledge in the chemical neighbourhood. We further validate by a manual literature search. See additionally Figure S27 and Table S4.

We then assembled a chemically diverse library of 539 unique compounds, with 386 sourced from the Prestwick library and filtered based on any reported cell surface interactions, and 153 sourced from Chemfaces natural product library based on a substructure search for an indole ring. We screened the library against all 7 hGPCRs, and a comparison to prior art revealed that the reduced dynamic range and sensitivity of HTR1A in yeast made this part of the dataset unreliable. Furthermore, 36 compounds were found to dampen luminescence independently from hGPCR expression (Supplementary Figure S22-S23), rendering some negative hits (Z-score<-3) meaningless. This left a slightly reduced (3,018 remaining out of 3,773 measured DTIs), but high-quality, dataset with a 98.7% overall correspondence to prior art, i.e., from mammalian reporter cell lines (Figure 4c and d, supplementary Figure).

### Correlating Predictions with Experimental Values

Figure 4e and f relate the predicted labels with the experimental labels, which shows that CSNN are a viable pre-screening method to reduce the experimental load by removing non-interacting DTIs with a high specificity. In Figure 4e argmax predictions have a specificity of 0.992 for the ‘No Effect’ class, showing a high discriminative ability. With the 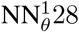 label prediction model, a higher number of DTI’s can be predicted due to inference considering all data in the chemical neighbourhood. Still here we found a high specificity (0.979) for the ‘No Effect’ class. Remarkably, however, other labels which should show a high correspondence with experimental labels (‘Partial Agonist’ and ‘Agonist’) have a low specificity and precision. Many predicted agonists have non-significant hGPCR activation. Labels such as ‘Antagonist’ and ‘Inverse Agonist’ are not expected to show any correspondence, due to the nature of the assay. Nevertheless, the methods developed can reliably be used to sort out true negative DTIs, which reduces the experimental workload to validate adverse or otherwise off-target effects significantly.

Finally, in Figure 4f, the experimental and predicted data is used to retrospectively assess the performance as a pre-screening methodology for DTI predictions. A experimentally significant ‘Hit’ is defined as a Z-score > 3, with the predicted label being assigned as a hit if (1) the label is not ‘No Effect’ and (2) the predicted *K*_*i*_ < 100 nM (using the regression model). The resulting confusion matrix shows that the methods are accurate for filtering out true negative DTI pairs (specificity = 0.948), but cannot reliably discriminate true positive hits (precision = 0.182). When combining the class label and regression model predictions, one can significantly enhance the DTI hit-rate (Fig. 4f), from 3.3 % to 18 %, thus reducing the experimental load to validate DTIs. This demonstrates CSNN as a useful, pre-screening methodology which can assist, but not replace, validation experimentally.

### Discovering Novel DTIs

In addition to DTI predictions made by CSNN, the underlying CSN can be used to probe the degree of novelty of measured DTIs with significant RLU, as all public database knowledge is query-able and accessible in a single CSN. Using this, we filter the experimental DTIs for significant hits (|Z-score| > 3) and without any neighbourhood data on the given hGPCR. We identified 14 novel hits without prior art on the hGPCR in the chemical neighbourhood. Six of these hits are highlighted in Figure 4g, and each of these represents new information on approved drugs. Tinoridines’ association with HTR4 adds indications for treating gastrointestinal disorders *(28)*, whereas Nicergoline interaction with CHRM3 indicates potential uses for treating Alzheimer’s disease and schizophrenia *(29)* and at the same time adds a potential mechanism of action to Nicergoline’s primary use, as CHRM3 is directly tied to blood-pressure control *(30)*. Similarly, the mechanism of action of Latanoprost for mitigating increased intraoccular pressure may be partially explained by the observed interaction with CHRM3. Some of these interactions also correspond well with known side-effects of the drugs, such as disturbances in the circadian rhythm associated with overdosing of Tranylcypromide, which is likely caused by the discovered interaction with MTNR1A*(31)* (See Supplementary Table S4). Thus, we further demonstrated the complimentary between a robust ML method and a high-throughput yeast biosensing system to discover and validate novel DTIs.

## 3 Discussion

hGPCR drug-activity databases are growing at astonishing rates, yet most compound-hGPCR interactions are absent. This tension remains unresolved by ML, which is routinely used to predict activities of new compounds, but rarely addresses how to impute missing data robustly in these databases. We have developed neighbourhood-to-prediction architectures which use labels as a feature and consistently filled out the gaps among the 23 million possible compound-hGPCR interactions. We further increased coverage from 1.5 % (370 K) to 5.6 % (1.3 million) of all possible DTIs in the database, robustly using a transparent training free prediction.

ML methods, akin to database sizes, are also growing at astonishing rates, yet often with little-to-none experimental validation as to the utility of the model. Furthermore, few guard-rails are imposed in terms of OOD detection, and public web-servers will produce predictions for any SMILES compound, regardless of the distance from the training set. We document this effect here and develop CSNN to prevent such OOD predictions, in which “No neighbourhood” implies no prediction. OOD predictions are problematic for experimental scientists which readily run into null-results. We demonstrate the applicability of neighbourhood-to-prediction methods to reduce OOD predictions, increasing the reliability and robustness of ML methods for drug-discovery. LaFs are strong inductive biases in homophilous networks and enable training free predictions which are robust, accurate, and interpretable. Finally, we complemented the *in silico* DTI discovery pipeline with a synergising HT-yeast platform to uncover novel DTIs. In particular, we utilise how our method is able to predict both mode of action and binding affinity to show a Pearson correlation of 0.75 between measured DTIs in yeast and the binding affinity of predicted agonists. Furthermore, by using the chemical neighbourhoods as a knowledge graph, we validated 14 novel compound-hGPCR interactions not described in literature or public databases, tentatively linking some with adverse effects. We imaging this utility could be used to pinpoint underrepresented chemical groups for future HTS campaigns on hGPCRs, which is relevant particularly when searching for novel patentable drug candidates, and at the same time validates off-target DTIs.

Chemical space networks face significant criticism, as subtle molecular modifications can change the toxicological and pharmacological behaviours of compounds *(32)*. However, our results indicate that reliable conclusions can be drawn from chemical neighbourhoods (in a training free manner) to correctly assign a bioactivity class and regression labels. This demonstrates the inherent network homophily in the hGPCR-compound space and offers efficient ways to increase coverage on the otherwise sparsely annotated drug-target interactions.

The results also link with a more fundamental discussion on whether graph neural networks require network homophily *(33, 34)*. The recent LaF framework demonstrates that advanced GNNs are on par in performance with training free GNNs, but this paradigm only works on homophilous graphs *(11)*. As we show in a benchmark with pdCSM, architectures without LaFs are outperformed by a training free prediction with LaFs on 15 out of 24 targets, and that this scales with network homophily. Thus, we demonstrate the utility of CSNN in using this advantage. Work has begun characterising “good” and “bad” network heterophily and building GNNs which are more robust to these properties *(33, 34)*. This is likely an interesting future line of work in the DTI setting for reliable database driven inference to uncover unknown DTIs.

The results presented here clearly demonstrate network homophily, while general discussion about whether this is inherent in chemical space is ongoing *(9, 35)*. Publicly available data does not sample chemical space independent and identically distributed (i.i.d), and deposited data is often a result of significant derivatization efforts to increase ligand efficiency, reduce toxicological properties, and increase availability pharmacokinetically. Thus, homophily is ‘over-sampled’ as often derivitisation efforts maintain the bioactivity class (e.g., Agonism), while improving other properties of the compound. This sampling bias then overestimates the true network homophily on which ML models are trained, which then retains the smoothness or locality inductive bias between the input and output spaces. This extends into train/test data splitting, leading to significant system leakage and an overestimation of model performance *(36)*. CSNN models outperform non-neighbourhood methods, but imply stricter requirements for reducing system leakage and thus evaluating models fairly. In the paradigm of LaFs, it will be increasingly critical to fairly evaluate and develop robust ML systems for DTI discovery in combination with experimental validation assays to assess true performance.

We anticipate that (i) neural networks that operate on the graph structure of chemical neighbourhoods rather than the graph structure of compounds using LaFs will enhance robust identification of in-distribution DTIs and limit OOD outliers, and (ii) that an increasing number of hGPCRs will become available to yeast and expand on the robust and cost-effective yeast platform to accommodate future DTI discovery. In fact, LaFs are not entirely new to biology: Alphafold2 has a ‘template’ enabled mode *(37)*, which can be viewed as a label of a neighbour. The label provides an approximate answer, which may be integrated to form a refined answer. By analogy, LaFs for DTI predictions provide an approximate answer for the bioactivity class and value of a DTI, which may be refined by a ML model. Point (i) is underway with established algorithmic *(15, 38)* and neural *(39)* network-based methods, yet remains limited to paired compounds in the latter case. Here, we take a significant step toward using LaFs on an arbitrarily large number of neighbours by using a one-vs-all database search during inference by CSNN. What remains to be done are end-to-end differentiable and explainable *(40)* neighbourhood-to-predictions. Point (ii) is also well-underway, with over 70 hGPCRs already reported to signal in yeast *(41)*. Neighbourhood-to-predictions architectures should additionally benefit biotechnology by rapidly identifying a suitable hGPCR for a given ligand to accelerate the development of sustainable biosynthesis of complex natural products and derivatives by utilising state-of-the-art microbial cell factories *(42, 43)*.

## Acknowledgements

This project has received funding from the Novo Nordisk Foundation, grant number NNF20CC0035580, and European Union Horizon 2020 research and innovation program (grant number 814645 (MIAMi) to MKJ). We would like to thank William Shaw for providing access to his yeast platform strains, which we used to build our hGPCR signalling yeast strains. This work was further supported by Danish Data Science Academy (to NGM), which is funded by the Novo Nordisk Foundation (NNF21SA0069429) and VILLUM FONDEN (40516).

## Funding

This work was supported by

The Novo Nordisk Foundation Center for Biosustainability (FGH, NGM, LGH, BL, TJ, CGAR, MKJ, EDJ)

The Danish Data Science Academy (NGM)

European Union Horizon 2020 research and innovation program (MKJ)

## Author contributions

Conceptualization: FGH, NGM, MKJ, CGAR, EDJ

Methodology: FGH, NGM, BL

Investigation: FGH, NGM Visualization: FGH, NGM

Chemical Library assembly: LGH, TJ

Funding Acquisition: MKJ, JDK, CGAR, EDJ

Supervision: MKJ, CGAR, EDJ

Writing - original draft: FGH, NGM, EDJ

Writing - review and editing: FGH, NGM, MKJ, CGAR, EDJ

## Material and Methods

### Computational methods

#### Graph Construction of Chemical Space Networks

The chemical space network can be formalised as a graph: 𝒢 = (𝒱, ℰ) with vertices (nodes) 𝒱 and edges ℰ, which may be parameterized by the adjacency matrix **A** ∈ ℝ^|𝒱|*×*|𝒱|^. Each unique compound corresponds with a unique node in 𝒱. To obtain this adjacency matrix, a similarity metric is required between any two compounds *(sim(d*_*i*_, *d*_*j*_), where *sim* : 𝒱 × 𝒱 → ℝ ∈ [0, 1]) associated with a real-valued vector (**d**_*i*_, **d**_*j*_ ∈ ℝ^*p*^). A natural definition is the Tanimoto similarity, which may be written in vector notation as:

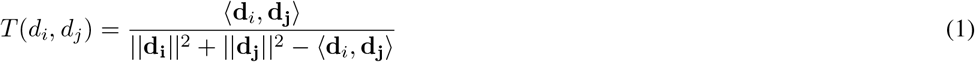

where ⟨**d**_*i*_, **d**_**j**_⟩ is the inner product, and ||**d**_**i**_||^2^ is the Euclidean norm squared. A näive implementation would calculate all pairwise Tanimoto similarity for the set of objects in *D* = {*d*_1_, *d*_2_, …, *d*_*n*_} (scaling as O*(n*^2^)). A graph could then be constructed by considering neighbours for which *sim(d*_*i*_, *d*_*j*_) ≥ *ϵ*. This is known as the *all-pairs similarity search* (APSS) problem, which Anastasiu et al. *(44)* have optimised by incorporating similarity bounds, thus significantly reducing the number of floating point operations. The approach, naturally, further exploits the commutative property (symmetry) of the Tanimoto similarity metric *(sim(d*_*i*_, *d*_*j*_) = *sim(d*_*j*_, *d*_*i*_)). Given a minimum threshold similarity *(ϵ)*, one can exclude objects whose norm are either too small or too large to be neighbours. We found that an all-vs-all pairs similarity search was necessary to accurately construct chemical neighbourhoods as low-dimensional embedding distances did not necessarily represent chemical similarity (Figure S26).

For a similarity threshold of *ϵ >* 0.4 we compute an all-vs-all similarity metric using Tanimoto similarity. Values less than ≤ 0.4 are set to zero to keep a sparse adjacency matrix (otherwise a computer runs into memory limitations 186K × 186K possible edges). We take inspiration from *(45, 46)*. Compounds are encoded with a 4096 bit Morgan fingerprint in RDKit (Chirality=True, radius=3) and concatenated with the 166 bit MACCS key substructure fingerprint. The bit strings undergo lossless compression by packing each successive 8 bits into a single byte while padding the end to ensure it is a multiple of eight, thus reducing the dimensionality of the input vector to **d**_**i**_ ∈ ℤ^533^. As bytes may have shared bits, the Tanimoto similarity of the packed bits is always T(A’,B’) ≤ T(A,B), where A’ and B’ are packed bits and A and B are unpacked bits. The similarity metric is computed in batches using SimSIMD^1^. The obtained adjacency matrix is upper triangular and is made symmetric by *A*_*sym*_ = *A* + *A*^⊤^. This adjacency matrix has 187,254 unique nodes corresponding to all unique compounds in the acquired ChEMBL data with the addition of the 550 compounds for the experimental library. To query related compounds the n-hop neighbourhood is used 𝒩_*j*_*(n)* = {*i* | *d(j, i)* ≤ *n*, ∀*i* ∈ 𝒱} where *d(j, i)* is the graph distance between vertex *i* and *j*. Practically, we restrict lookup to the 40 nearest neighbours for the training/testing/validation data due to compute speed in constructing neighbourhoods and to a 2-hop neighbourhood for visualisation tasks.

#### Data partitioning for Training, Testing and Validation

For all 550 compounds in the screened chemical library, we identify all molecules within a 2-hop neighbourhood and remove any compounds with over a 0.65 Tanimoto similarity to prevent data leakage for the experimental validation. Then, remaining compounds are assigned a Murcko scaffold index, and the data is split into a 80/10/10 train/test/val split using code adapted from https://github.com/chainer/chainer-chemistry/blob/master/chainer_chemistry/dataset/splitters/scaffold_splitter.py. This ensures that scaffolds in the test dataset are not present during training.

For each compound in each split we explode the pandas array such that each unique compound-receptor interaction(DTI) is a separate row. This facilitates a comparison with conventional drug-target interaction methods. We then precompute a compound representation **x**_*i*_ based on a pre-trained message-passing neural network from *(47)*. This numerically represents the chemical graph (MPNN at atom level with ChemProp) allowing for message-passing on the level of chemical neighbours rather than compounds. We choose learned representations as opposed to chemical fingerprints as they consistently outperform them *(40, 48)*. In the case of the benchmark in 2, we re-implement the fixed RDKit molecule representations (without feature selection) using the Supporting information from the pdCSM paper. This allows for a direct comparison, removing the effect of the compound representation. Due to lacking training and inference code, however, we could not entirely replicate their workflow. Correspondence with authors was unsuccessful (no response).

Likewise, we pre-compute target representations **p**_*i*_ using the ESM-2 protein language model and the canonical UniProt sequence (esm.pretrained.esm2_t6_8M_UR50D()) *(49)*.

For each unique compound-receptor data-point *(i)* we prepare a graph G_*i*_ and label **y**_*i*_ using the accessible graph data framework by PyTorch *(16)*. The graph is structured as 𝒢 = Data(**x**, ℰ, **e, p, y**), where **x** are the compound representations for the chemical neighbourhood and compound *i*, ℰ are the unweighted graph edges, **e** are the edge features, **p** is the target representation, and **y** is the label. The compound representations are **x** ∈ ℝ^|𝒱|*×*300^. The edges in ℰ are always directed and incident on the node to be classified, as illustrated in Figure 1d. The edge features are **e** ∈ ℝ^|ℰ|*×*7^ which includes the one-hot encoding of the neighbouring compound bioactivity (LaF in effect) as well as the Tanimoto similarity defined in the adjacency matrix. This allows for the node-classification task to be rephrased as a graph classification task. This also enables the Label as a Feature framework delineated in Section S3. As PyTorch requires **p** ∈ ℝ^|𝒱|*×d*^ we broadcast the target representation, which is only of dimension 1 × 320 (ESM-2, *(49)*). **y** is a one-hot encoding of the bioactivity class such that **y** ∈ 0, 1^6^. We provide the torch files dataset_train.pt, dataset_test.pt, dataset_val.pt to enable subsequent use and bench-marking of future methods *(50)*. An i.i.d sampled version is also provided, the results are comparable for both splitting methods.

### LaFs: Training-free argmax(.) predictions

To demonstrate the efficacy of training-free predictions using LaFs we compute the most likely label algorithmically. We sum over the all **e** for neighbouring nodes excluding the column, which represents the Tanimoto similarity, then divide by the number of edges to obtain a normalised output (frequency). The output for a given graph is a vector **e**_*tot*_ ∈ ℝ^6^ which the argmax(**e**_*tot*_) may be taken and compared with the ground truth label argmax(**y**) to determine the performance metrics. Intuitively, if the chemical neighbourhood has a majority of agonists, the compound in question is likely also an agonist. If network homophily is the case, then training-free prediction using LaFs will be a good approximation of the true label.

In Figure S3, the metrics are computed across all graphs in test, train, and validation to obtain the predicted and true labels for each graph. As the output is deterministic, no uncertainty is reported. Then common performance metrics are calculated (See later section).

### LaF: RF + N

Drug-target interaction (DTI) commonly has compound-to-prediction architectures, where a compound is represented with a numerical vector and passed through a ML architecture. In the case of simply ML models (RF or MLPs) we extend this to include LaF, by simply concatenating the compound representation, receptor representation, and the aforementioned frequency vector of the neighbourhood (RF + N) or simply the receptor and compound (RF) to evaluate the effect of +N (LaF).

The input vector is of dimension 300 + 320 (pretrained compound representation and ESM-2 target representation) and the output is the one-hot encoded bioactivity class label **y** ∈ 0, 1^6^. We use scikit-learn *(51)* and the ensemble random forest (RF) classifier as well as a simple multi-layer perceptron (MLP) as the baseline models. A hyper-parameter grid search is carried out with 2-fold cross validation:

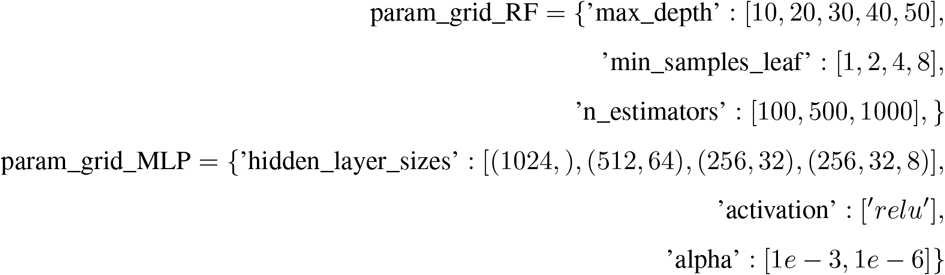

and the best parameters are chosen. Fixed parameters for the RF are class_weight=‘balanced’. For the MLP max_iter=2000, early_stopping=True, validation_fraction=0.1, n_iter_no_change=200.

An additional neighbourhood based RF + N and MLP + N is trained using the same hyper-parameters where the input vector is additionally concatenated with the normalised **e**_*tot*_ presented previously. This adheres to the permutation invariance property desired and is thus a valid although crude construction.

Five independent training’s are carried out to calculate the mean and standard deviation on the performance metrics, thus assessing the stability of training.

### LaF: increasing the expressive power of GNNs with CSNN

The chemical space neural network (CSNN) is designed as illustrated in Figure 2d. We instantiate CSNN as a weaker form of message-passing *(52)*, namely a graph convolutional neural network (GCN) with edge attributes ^2^. In the first block, a local representation is computed by aggregating information from neighbouring nodes and LaF using edge attributes to form **r**_*i*_ from the local latent representations 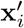:

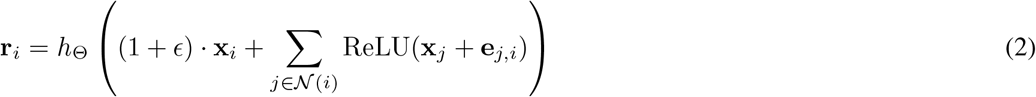

As the graph is directed upon the node with a missing label-to-be predicted (with only a 1-hop neighbourhood), no graph pooling is needed and a single GCN layer suffices: as the output dimension is already one-dimensional and represents a pooled representation.

We use an output dimension of 256 for the local representation using a hidden size of 256 and apply a ReLU() non-linearity. Then the local representation is concatenated with the target representation and the normalised **e**_*tot*_, which acts as skip connection, to form 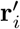. Finally, 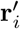 is transformed by a MLP (hidden size 512, 2 hidden layers, output dimension: 6) as:

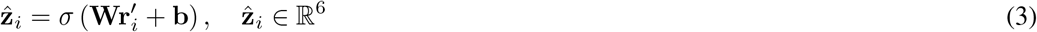

to form the predictions from which the logits are used to compute the cross-entropy loss. We compute a batch-wise loss and train with a batch size of 16 for 100 epochs with the Adam optimiser *(53)*, a learning rate of 0.00001 and weight decay of 0.0001:

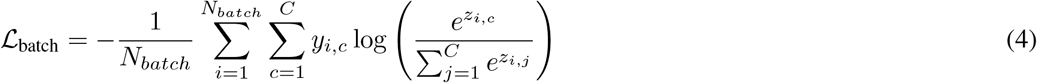

where *C* is the number of classes and *N*_*batch*_ is the number of batches of batch size 16.

### LaF: Multi-label and Multi-class

The previous method predicts only class-wise probabilities for a given drug-target interaction (receptor encoded by ESM-2 embedding), specified by the compound representation, target representation, and it’s chemical neighbourhood. To predict class-wise probabilities for all 128 hGPCR receptors and all six classes (multi-label & multi-class), we adapt the previous method by including all available data in the edge features (128 + 1) dimension with the class index at each position and 0 if no data is present. This is expanded to a one-dimensional vector of (128 × 7 = 896) to allow for a one-hot encoding of neighbouring data for the six-classes and the no data index. We use the same skip connection as previously but a output size of the GCN as 64. The concatenated 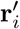 is of dimension 960 before transformation with a MLP. The final output is a one-dimensional vector of dimension 896 (128 × 7), which may be reshaped to a tensor of dimension 1 × 128 × 7 with a softmax applied along the third axis to obtain receptor-wise class probabilities. The true output is of dimension 1 × 128 and represents the class (in integers) for all 128 receptors. As no compounds have data for all receptors, the positions which are zero (the no data index) are masked when computing the loss on the output logits. We train with the Adam optimiser, a learning rate of 0.00005, and a weight decay of 0.0001. While inference produces class-wise predictions for all receptors, the performance metrics naturally only consider those with data available by using a mask.

#### 3.0.1 LaF: Regression Benchmark

We source the pdCSM training and test set from https://biosig.lab.uq.edu.au/pdcsm_gpcr/ and obtain the performance metrics on the blind test set from the supplementary information in *(17)*. This was done because training and inference code was not available, neither provided upon reasonable request. Thus, performance in the comparison has pdCSM models with feature selection vs. our equivalent models (RF + N) on the reproduced compound representations (using their SI).

To uphold the transductive node classification restriction, we allowed training data to dynamically refer to each other during inference and training: labels of neighbouring compounds within the training set are used as features. For the test set, the neighbourhood graph is constructed using only training data compounds and labels as features.

Of key importance, the pdCSM method trains a single model for each receptor. Instead, we concatenate the ESM-2 representation and train a single model on all training data across all receptors. We found that pooling the data in this fashion lead to increased performance (a single model see’s more data).

In Figure 3e the Pearson correlation coefficient is given per receptor (one point = one receptor) under the homophily assumption. This uses the mean-value of the labels in the neighbourhood as the prediction. Then the correlation coefficient is calculated per receptor between the predicted and true values. As illustrated, the models performance scales linearly with the degree of network homophily (for which the mean value is a good approximation). This is a highly receptor dependent effect: CHRM3 has a *ρ* = 0.87 while MTNR1A is more heterophilic *ρ* = 0.42. The receptor-wise breakdown is given in the table below:

##### Performance Metrics

For the compound-to-prediction methods (RF and MLP), the corresponding neighbourhood-to-prediction (RF + N, MLP + N,), and CSNN using LaFs, the performance metrics are calculated for each bioactivity class *i* as:

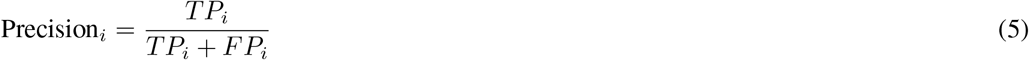

where *TP*_*i*_ is the number of true positives and *FP*_*i*_ is the number of false positives. The recall is calculated as

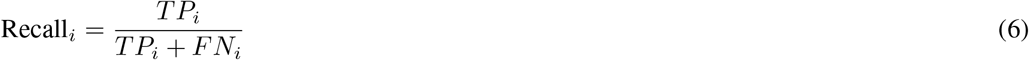

where *FN*_*i*_ is the number of false negatives. The F1 score is calculated as:

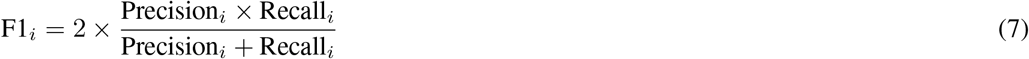

To summarise across all bioactivity classes, a weighted F1 score is computed:

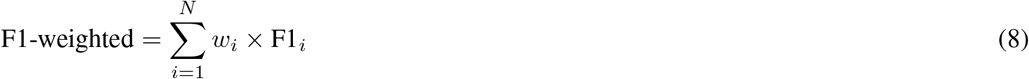

where *w*_*i*_ is the proportion of true instances for class *i* out of all instances, and *N* is the number of classes. For regression tasks, the Pearson correlation, Kendall correlation, and Spearman correlation coefficient, mean average error (MAE), and mean square-root error (MSE) are calculated with sklearn.

##### Receptor distances and clustering

ESM-2 distance metrics were obtained by calculating the cosine similarity between all-pairs. Obtaining a distance metric from a shared chemical space between any two receptors was calculated as follows: For a given receptor pair *i, j* the dataframe was reduced to only consider compounds for which data was available for both receptors. Then we calculate a corrected Cramer’s V statistic, which measures correlation between two variables which are categorical. If less than 5 co-occurring or no co-occurring compounds existed, the distance was set to its maximum (1). Highly correlated hGPCR receptors were assigned a value of (1 - Cramer’s V), thus a distance of zero for completely correlated receptors. The phylogenetic trees in Supplementary Figure S19 were generated from distances calculated between all 128 selected receptors based on; 1. ESM-LLM distance metrics *(49)*, 2. from the chemical space they are able/unable to sense, and 3. from a multiple sequence alignment using muscle 5.1*(54)*. Three distinct trees were then generated using the Bio.Phylo package*(55)*, and visualised using ETE V3*(56)*.

##### Z-score Analysis

For each molecule-receptor pair a single-point relative luminescence unit (RLU) was measured as described above, then log_10_ transformed, subtracted by the mean *(µ)* RLU for the given receptor, and then divided by the standard deviation *σ* for the given receptor according to:

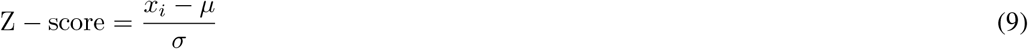

where *x*_*i*_ is the log_10_ raw RLU signal. log_10_ transformation prior to Z-score analysis was necessary to attain a Gaussian distribution, which is amenable to Z-score analysis.

To rescue outliers a modified Z-score was used, as defined by:

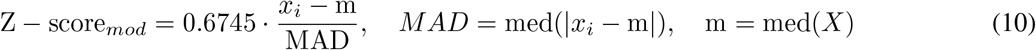

where *x*_*i*_ is the log-transformed RLU signal, MAD is the median absolute deviation, and *m* is the median value for a given receptor. The effect of the modified Z-score is qualitatively shown in Supplementary Figure S24, which illustrates a broadening of the kernel density estimate fit with respect to the canonical Z-score. The figure further shows the cut-off for the hit/no-hit threshold which is set at | Z − score | > 3.

#### Experimental methods

##### Chemical library

386 compounds were sourced from the Prestwick library, selected by reported receptor associations and chemical diversity, stock concentrations at 10mM. 153 compounds were sourced from Chemfaces chemical library, selected by a substructure search for the indole ring moiety, stock concentrations acquired at 1mM. All chemicals (Supplementary File S1) had a reported purity of 95% or higher

##### Gene Synthesis

All biosynthetic genes used in the current study are listed in Supplementary Table S7. Genes were synthesized by IDT (San Diego, USA) and if stated, optimized for expression in S. cerevisiae.

##### Cloning and yeast transformation

All plasmids were constructed by USER cloning *(57)* and propagated in E. coli DH5*α* competent cells. Genes for integrative Easy Clone - marker free vectors were prepared according to Jessop-Fabre et al. *(58)* and the gRNA expression cassettes as previously described *(59)*. All plasmids were verified by Sanger sequencing prior to yeast transformation. The integrative plasmids were linearized by NotI enzyme (New England Biolabs, MA, USA). The yeast transformations were done using standard Li-acetate methods according to Gietz and Schiestl *(60)*. The integration of heterologous genes was verified as described in CasEMBLR *(58, 59)*, The strains generated in this study are listed in supplementary Table S8.

##### Yeast strains and cultivation

Yeast strains ScS1-7 expresses NanoLuc upon stimulation of their respective human G-protein coupled receptors. The parental strains ScFH236 and ScFH237 were generated by changing the sfGFP reporter to Nano-Luciferase (NanoLuc) in the platform strains yWS2266 and yWS2267 published by W. Shaw et al. *(25)*. This was achieved by transformation of a transient Cas9 plasmid (pCfB5270), transient sfGFP specific gRNA plasmid (pFH69), and a repair template with homology to the upstream promoter and downstream part of sfGFP, containing the NanoLuc coding sequence and the tCYC1 terminator, which was linearised from a plasmid vector (pHT9). ScS1,2,3,5, 6, and 7 were generated by integrating 5HTR1A, 5HTR4, ADRA2A, OPRM1, CHRM3, or MTNR1A, respectively into ScFH237, ScS4 was generated by integrating ADRA2B into ScFH236. This design combines advantages from two of the most well known yeast GPCR biosensors, described in Kapolka et al. and Miettinen et al. *(18, 19)*. To generate biomass for the assays, yeast was inoculated from cryostock to 5 mL SC-trp and incubated ON at 30 °C 250 rpm. The next day, 20 hours before the assay, 5ml of saturated culture was added to 45ml SC-trp in a 500ml shakeflask and incubated overnight 30C 200RPM. Next 25 mL culture was centrifuged at 5000 x g for 5 min in 50ml falcon tubes, and the supernatant was discarded. The pellet was then resuspended in pH buffered SC-trp at pH 7.2 *(61)*. OD600 was measured and adjusted to 5 before continuing with the procedures described in “Bioactivity assays”

##### Chemical preparations

For the DRCs shown in Figure 4d and supplemantary Figure S20, DAMGO-Enkephalin acetate salt, Serotonin HCL, Acetylcholine chloride, Epinephrine bi-tartrate and Melatonin were acquired from Sigma-Aldrich and 10X stocks of each were prepared in sterile mq water containing 10% DMSO, in the concentrations listed in supplementary file S3, including a no-ligand control of only 10%DMSO in sterile Milli-Q water. The ChemFaces and Prestwick chemical libraries were acquired pre-solubilised in DMSO at 1mM and 10mM respectively. 10X stocks were prepared in 96 well plates (Greiner 655160) by diluting a fraction of the supplied stock to 100*µ*M in sterile mq water, with a final DMSO concentration of 10%. One positive control and three negative control wells were reserved on all plates containing the chemical libraries.

##### Bioactivity assays

For the chemical library screen, all 10X stocks aswell as a no-ligand control were diluted 1:10 in 96 well plates (Greiner 655160) in cell cultures prepared according to “yeast strains and cultivation”, resulting in 1 well pr. receptor / compound combination with 100*µ*L of sensing cells and 10*µ*M of compounds. A gap of 4 minutes between loading of each plate was maintained to allow sequential measurements in a SynergyMX microtiter plate reader (BioTek). For the DRCs shown in Figure 4, 3 replicates of each concentration of ligands with relevant receptors were prepared aswell as a no-receptor strain control for each ligand. For supplementary Figure S20, a 1% v/v final concentration of DMSO control was run in triplicate along with triplicates of each of the selected compounds from our chemical library at 10*µ*M concentrations (1% v/v DMSO). All plates were covered with a breathe-easy breathable membrane and incubated for 4 h at 30 °C 300 rpm. Twenty minutes before incubation was completed, a lysis mix of 1.33% v/v Furimazine stock (NanoLuc substrate) from Promega in CelLyticTM Y cell lysis reagent (Sigma-Aldrich) was mixed and 12 *µ*L was distributed to wells of white small volume 96-well plates (Greiner 675083). Following incubation, the plates were vortexed for 5 sec. before the membrane was removed and 4 *µ*L of each well containing sensing cells was transferred to the plate containing the lysis mix and the plate was incubated at RT for 18 minutes and then placed in a SynergyMX microtiter plate reader (BioTek). The luminescence was measured at 3 time points for the chemical library screen: 18.5, 20, and 21.5 min. after mixing, and 2 time points for the DRCs: 20 and 21 min. after mixing. The settings were filter-luminescence, with gain 150 and 0.5 sec. integration pr. well, the average of each of those time points is reported as single replicates pr. well.

##### Dose response curve calculation

All DRCs shown in Figure 4d and supplementary Figure S20 were generated in Graphpad Prism 9.5.0. The models chosen for HTR4, ADRA2A, ADRA2B and MTNR1A, were all standard 4-parameter non-linear regressions. The models chosen for HTR1A and CHRM3 were a bell shaped DRC, and a bi-phasic DRC respectively. Since the DRC of OPRM1 did not plateau at the higher concentrations, a 5-parameter non-linear regression was applied for the best fit for visualising the trend. All additional information related to the fitted curves is available in supplementary file S3.

## Footnotes

1 https://github.com/ashvardanian/SimSIMD

2 https://pytorch-geometric.readthedocs.io/en/latest/generated/torch_geometric.nn.conv.GINEConv.html#torch_geometric.nn.conv.GINEConv

