## Supplementary Information for "Labels as a Feature: Network Homophily for Systematically Discovering human GPCR Drug-Target Interactions"

**This PDF file includes:**

supplementary Figures S1-S27

supplementary Tables S1-S10

**The following supplementary material for this manuscript is currently available upon request, and will be deposited in a public file repository upon publishing in a peer-reviewed journal:**

S1. Annotated dataset containing smiles, raw RLU, calculated Z scores, lists of nearest neighbors, their bioactivity label, and the WPA contributions of each entry, for all tested compounds and hGPCRs.

S2. Data file of cost analysis of hGPCR-screens

S3. Complete annotated and labelled dataset of prior art on the 128 included hGPCRs (Sourced from ChEMBL and IUPHAR/BPS Guide to Pharamcology (*12*, *13*).

S4. Concentration ranges, corresponding measured RLU, and fitted curve values from GraphPad Prism. Corresponding to data in Supplementrary Figure S20.

S5. Zip folder of Pytorch files containing our CSNN implementation

### Supplementary Notes

#### S2 Prior Work

The chemical homophily principle was developed together with Chemical space networks (CSNs), which allow for visualisation and interpretation of chemically related molecules. CSN's and similar methods have been developed to rationally engineer drugs by considering the chemical neighbourhood and building structure–activity relationship (SAR) (*46*), leading to canonical domain-specific language such as 'activity cliffs' (*35*). However, these methods are difficult to scale due to the quadratic dependence on number of compounds and arbitrary design choices in their construction and thus been decreasing in utility (*32*, *62*). Chemical similarity thresholds are essential in limiting the connectivity of CSNs to enable interpretation. (*32*). These network-based methods have found their way to established chemical bioinformatics companies (e.g. Chemaxon with Neo4j[3]) and have been key tools for SAR campaigns (*46*). Preliminary results on successfully utilising CSNs for lead-compound identification were recently demonstrated (*63*), using an edge propagation technique. Nevertheless, interpolation remains challenging, as the chemical space is insufficiently annotated and sparse.

Concurrently and analogously to CSNs, knowledge-graphs have emerged as expressive ways to mine and infer information from large relational-databases using dyadic interactions. Knowledge graphs are widely used across various fields: in search engines to improve search results, at the interface with machine learning models for knowledge retrieval (*64*), and in diverse biological applications (interactome like data) (*65–67*).

Recent work has begun reconciling knowledge graphs with graph neural networks that operate on them, for knowledge-aware inference in architectures termed: Knowledge Enhanced Graph Neural Networks (*68*, *69*). Our work formalises this framework in the DTI setting for hGPCR bioactivity prediction, building on previous work in other domains.

#### S3 A new paradigm: Labels as a Feature

An emerging paradigm known as training free graph neural networks (TFGNNs (Published: TMLR 8/2024) (*11*), show how labels as a features (LaF) is an admissible operation in a transductive node classification tasks: labels of neighbouring nodes are used to update the learned representations. The paradigm proves how this increases the expressive power over Graph Neural Networks (GNNs). Empirically the results show that even without training, prediction accuracy is incredibly high on common benchmark GNN datasets. Here, we demonstrate similar training free predictions that are on-par or outperform conventional trained compound-to-prediction architectures. Instead of framing the task as a transductive node classification task, we reframe it as a graph classification task with neighbourhood labels as edge features (Figure 1d) in a directed (transductive) graph.

#### S4 Explainable AI

Explainable AI (XAI) encompasses: Transparency, Justification, Informativeness, and Uncertainty estimation (*40*). The completely algorithmic predictions (argmax and mean value, taking the inductive bias of network homophily) are transparent (closed-form expression), justified (in relation to available data) but has low informativeness. A mean value prediction is not incredibly new nor useful information to target discovery or derivatization efforts. In the perspective of uncertainty estimation: while the class probabilities of the model do look calibrated (correlate with AUC-ROC and F1 scores), this is not a rigorous uncertainty quantification technique. In this respect, future work on neighourhood-to-prediction architectures could

[3]https://chemaxon.com/blog/presentation/neo4j_presentation

take great advantage of Gaussian processes (*70–72*) or Conformal prediction (*73*). The former works in smooth input spaces, which would likely work in homophilic networks, and the latter is a more flexible and rigorous uncertainty quantification technique which is underexplored in relation to black-box ML models for small-molecule property prediction (*74*). A natural extension to the method proposed are fully differentiable models, circumventing pre-computed compound representations and enabling feature-attributions, which is critical for explainable small molecule GNNs. Gradient-based feature attribution would then go back from prediction to the chemical neighbourhood (a list of compounds).

#### S5 Cost-efficient experimental chemical screening against GPCRs

When screening thousands of interactions, the price pr. well is a deciding factor on the total scope of a project. The calculation of these prices are, however not trivial, as there are many factors determining the total costs, such as salaries and differences in the laboratory requirements for handling mammalian cell lines, radioactive substances, or yeast. We chose to simplify this equation by only accounting for material costs associated with performing a screen of 4000 individual drug-GPCR interactions in vitro for all known methods available to us. Our combined calculations are available in supplementary File S2, and a summary of the available screen types and their associated advantages/disadvantages is available in supplementary table S2. This analysis identified yeast as the cheapest and most reliable option at 1$/well, where the cheapest alternative is Calcium staining assays on mammalian cell lines engineered to express the GPCR, for which we estimated a price of 1.8$/well.

#### S6 Identifying receptor independent effects

By normalising the raw reads of relative luminescence units with receptor-wise Z-score statistics, we were able to set a unified threshold of $|Z|>3$ ($p$-value $< 0.003$) for hit-determination, and the data could then be compared to prior art registered in our annotated dataset (Figure S22B and C). The distribution of hits across chemical space are available in Supplementary Figure S22A. Across all tested GPCRs, we observed 92% correspondence between prior art, consisting of 474 individual DTIs, which have a description in prior art and have been screened in this study (Supplementary Figure S22B). 381 of the 474 comparable DTIs are labelled with "no effect", and if only partial agonism and agonism is investigated, the correspondence drops to 68%. However, the majority of all discrepancies are caused by a single GPCR, as HTR1A performed exceptionally poorly, missing out on 19 out of 28 expected agonists and partial agonists, likely due to the reduced sensitivity and low dynamic range observed in the DRC (Supplementary Figure S20). When excluding HTR1A, the total correspondence rises to 89% for agonists and partial agonists, and to 95% including no effect DTIs. These results demonstrate the conservation of GPCR signalling in yeast across a variety of DTIs. The largest group of discrepancies were entries in ChEMBL and IUPHAR/BPS guide to pharmacology being classified as "no effect", which our screen identified as hits (supplementary figure S22). Hexachlorophene and chloroxine (known antifungals), also generally dampened the signal to a similar degree across all GPCRs (Except OPRM1 for chloroxine). Finally, bromocriptine displayed a strongly dampening effect on the CHRM3 GPCR only. By re-screening these compounds on the parental yeast strain ScFH237, we discovered that halogenated phenothiazines, low hydrophilic-lipophilic balance steroids, and all compounds carrying a 7-membered ring, dampened the luminescence signal independently from GPCR signalling (Supplementary Figure S23) and were consequently excluded from our downstream analyses.

#### S7 Other discrepancies

We observed cases where GPCR activation was measured despite prior art suggesting otherwise. For example, chloroxine activated OPRM1 (Z=8.8), and serotonin activated MTNR1A (Z=5.0) (Supplementary figure S23C). Both have previously been classified as having “no effect" on the respective GPCRs (CHEMBL1201862 (*75*), and CHEMBL1130532 (*76*), respectively). These could be explained by the comparatively high dynamic ranges on the yeast platform, which may have allowed it to detect DTIs that other screens might have missed. Chloroxine is known to bind OPRK1 (*77*), so based on phylogeny, it is not surprising that it also binds to OPRM1 as shown in our screen, despite the conflicting information from prior art. Activation of MTNR1A by serotonin has already been shown in other publications relying on yeast platforms (*18*, *43*), however, this information is not deposited on either ChEMBL or the IUPHAR/BPS guide to pharmacology, which is why an older assay, is the only reference on this DTI in the dataset ((*77*)). This demonstrates how the high dynamic ranges on the yeast platform (Supplementary Figure S20) can help catch DTIs, which other screening campaigns have missed. Finally, there is a total of 5 previously reported DTIs which the yeast based screen did not catch:

1. Nalbuphine on OPRM1
2. Tramadol on OPRM1,
3. Apomorphine on ADRA2A
4. Pilocarpine on CHRM3
5. Bethanechol on CHRM3

Nr. 2 can be explained, as tramadol is a prodrug, which is only active once oxidised by the human CYP2D6, which the yeast does not have, hence no activity is observed. The others are true mismatches, and could indicate a yeast bias on receptor signalling. In summary, our findings revealed 92 % correspondence between prior research in mammalian cells and the yeast platform, underscoring the viability of yeast as a cost-effective substitute for human cell lines (Supplementary Table SS6). HTR1A accounted for most discrepancies due to poor signalling in yeast, while other variations stemmed from compounds that reduced the reporter signal irrespective of GPCRs, indicating yeast-specific effects. By excluding GPCR-independent signal-dampening compounds and HTR1A, the quality of the gathered experimental dataset was drastically increased, and the correspondence to prior art was now 98%

#### S8 Chemical Space Network Construction in 20 Minutes

To realise CSNN, we first constructed CSNs from an adjacency matrix (Figure 2a), which encodes the graph structure connecting similar compounds (Figure 2b). We tackled two major challenges in large CSN utilisation:

**(i) Accelerated CSN construction** By using available tools to parallelize the calculation of Jaccard/Tanimoto similarity (*78*), we reduced the calculation of an all-vs-all CSN for 187 K compounds to 20 minutes on a home-computer (32.4 billion edges possible on a MacBook Pro 32 GB RAM, M1 Max @ 2.0 GHz x 10 cores). This makes the method accessible and can be done for any compound library. For moderately sized compound libraries (10 K), a CSN was established in 0.952 $\pm$ 0.002 seconds. In head-to-head comparison, published methods reported a 1 hour computation for 80 K compounds (*45*), our methods completed the same in 3.7 minutes, which was 16-fold faster than state-of-the art. The resulting computation was a sparse adjacency matrix which describes the connectivity of the graph (See Figure 1a) and forms the basis for querying and visualisation of data with CSNN.

**(ii) Open-access CSN tools to Query Available Data** We enabled the broader use of CSNs by developing a standalone user-friendly script, akin to (*45*), for visualising and utilising chemical neighbourhoods to understand the rapidly expanding datasets (Supplementary Figure S17), which are being procured. Our script queries the CSN and automates visual summaries of the neighbourhoods (Figure 1b). The visual neighbourhoods give an overview of available data and help infer possible DTIs of a given compound. These tools Querying of the hGPCR dataset and visual reporting was achieved in < 6 seconds, and an output .csv was produced with associated DTIs, ChEMBL ID, and compound information as well as Figure 1a,b.

### S9 Supplementary Figures

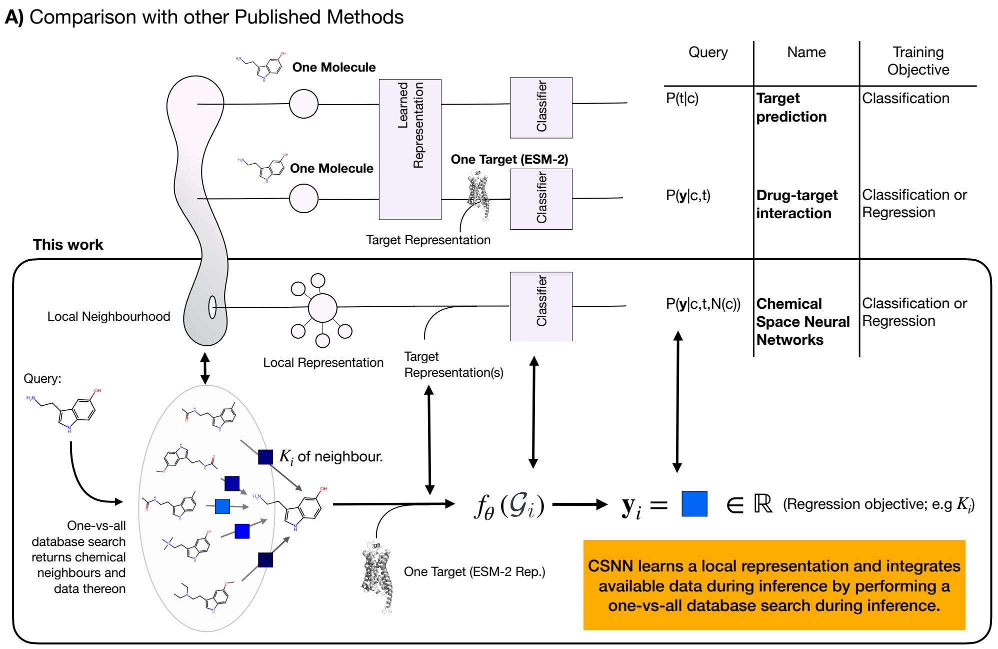


**Figure S1**: **Comparison of compound-to-prediction vs. neighbourhood-to-prediction architectures using LaFs**. Most drug-target interaction prediction methods indirectly either provide a conditional likelihood for the target given the compound P(tlc) or the conditional likelihood of activity y given the compound and target pair: P(ylt,c). These both take a *single* compound from chemical space (as visualised by the line) and use RDKit of MPNNs (like ChemProp) to produce a learned representation which is fed to a trained classifier. In the latter case, a target representation is concatenated to the compound representation. This work (bottom row), establises LaFs using neighbourhood-to-prediction architectures. An example for Serotonin is given to visually explain inference: the query is used in a one-vs-all database search which returns chemical neighbours and data thereon. This is a list of compounds and an accompanying list of labels (LaFs). Using a transductive node classification scheme, but re-framed as graph classification we can produce a single output by message-passing on the chemical neighbourhood graph. As stated in Section **??**, LaFs provide an approximate answer to guide ML model predictions.

**a)** NN$^6_\theta$ test metrics by class with only confident predictions.

| | | Precision | Recall | F1-Score | Support |
|---|---|---|---|---|---|
| | **No Effect** | 0,9713 | 0,9237 | 0,9469 | 4391 |
| | **Allosteric binding** | 0,9540 | 0,9706 | 0,9622 | 5234 |
| **Threshold > 0.8** | **Antagonist** | 0,9534 | 0,9854 | 0,9691 | 4031 |
| | **Inverse Agonist** | 0,9153 | 0,8438 | 0,8780 | 64 |
| | **Partial Agonist** | 0,9085 | 0,7771 | 0,8377 | 166 |
| | **Agonist** | 0,9091 | 0,9701 | 0,9386 | 268 |
| | Overall | 0,9576 | 0,9574 | 0,9571 | 14154 |

**b)** NN$^6_\theta$ test ROC-AUC at optimal class thresholds.

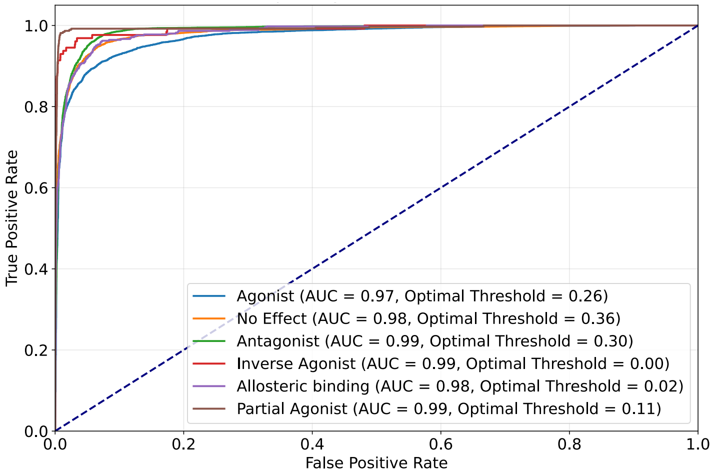


**c)** NN$^6_\theta$ test ROC-AUC non-optimised thresholds.

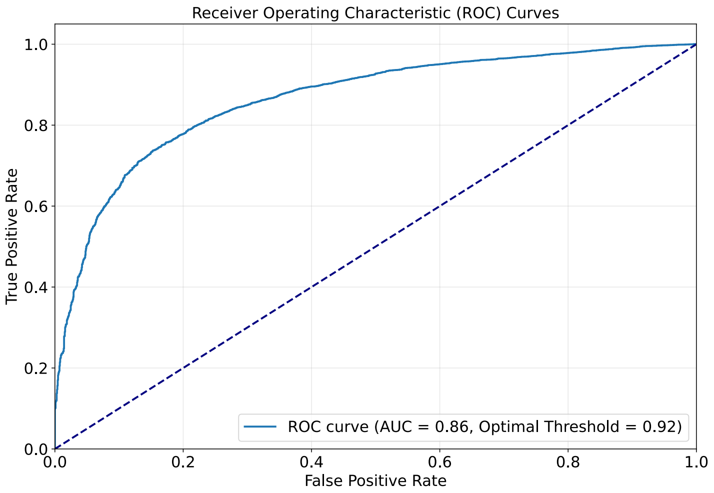


**Figure S2**: **NN$^6_\theta$ Test Metrics**. **(a)** Summarises the precision, recall, F1-score and statistical support for predictions with a class label probability > 0.8. Summaries are produces on a class-by-class basis. **(b)** ROC-AUC plots considering the class probability $p_i$ and the true label $y_i \in 0, 1$. The AUC is provided for optimised probability thresholds. **(c) Setting a single threshold for all classes significantly reduces model performance: AUC. Given results in Figure 2**g, S3-S5 we suggest user to use a probability threshold of > 0.8 or to look at the entropy of the logits.

**a)** $NN_{\theta}^{6}$ Confusion matrix on test data unnormalised (raw counts, left) and normalised (row-wise, right) at varying logits probability.

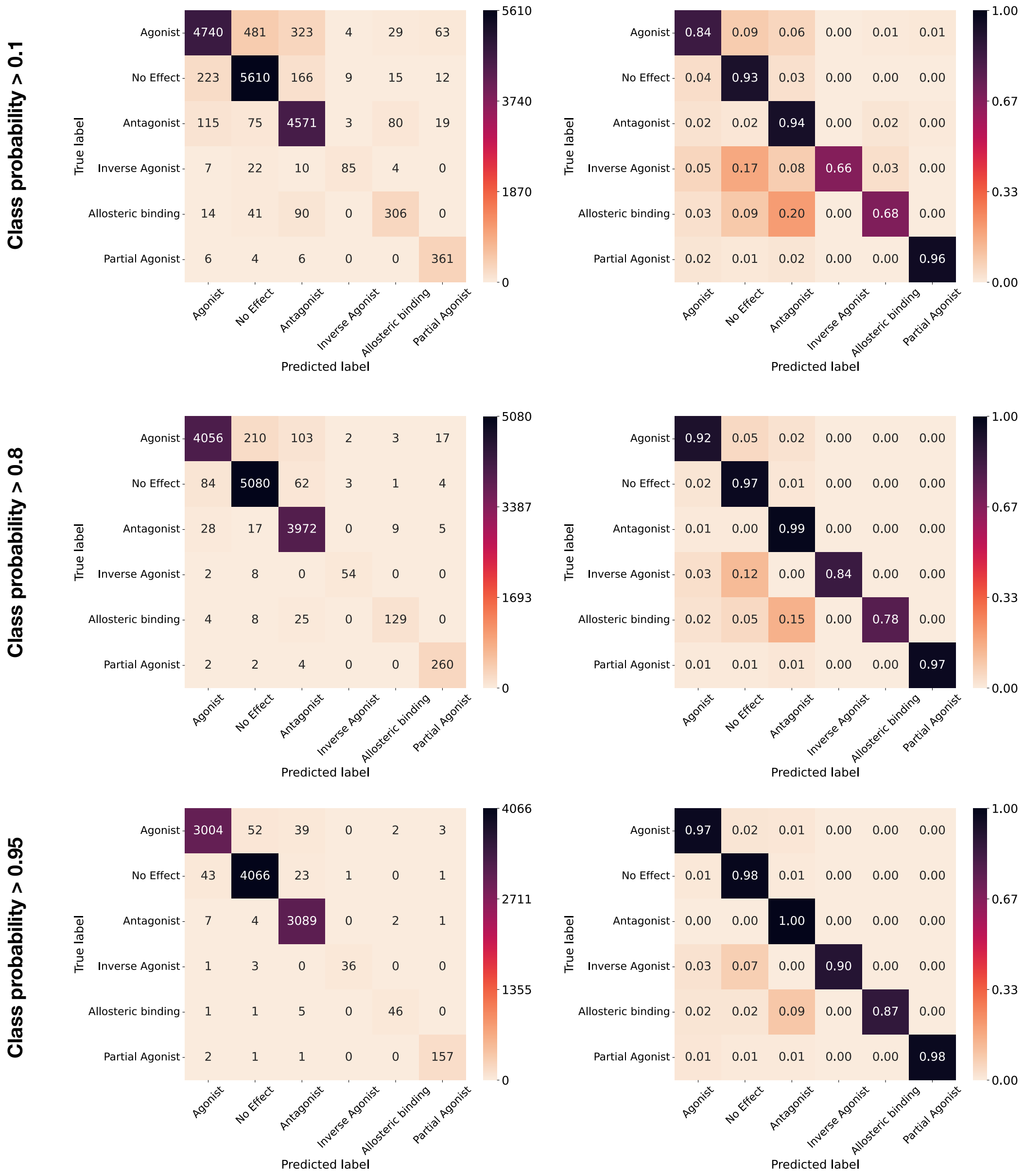


**Figure S3**: **Confusion Matrices for Filtered by Class Probability**. Un-normalised raw counts (left) and row-wise normalised confusion matrices (right). At a class probability of >0.8 the classification is excellent, even more so at 0.95 although with far less predictions passing the filter.

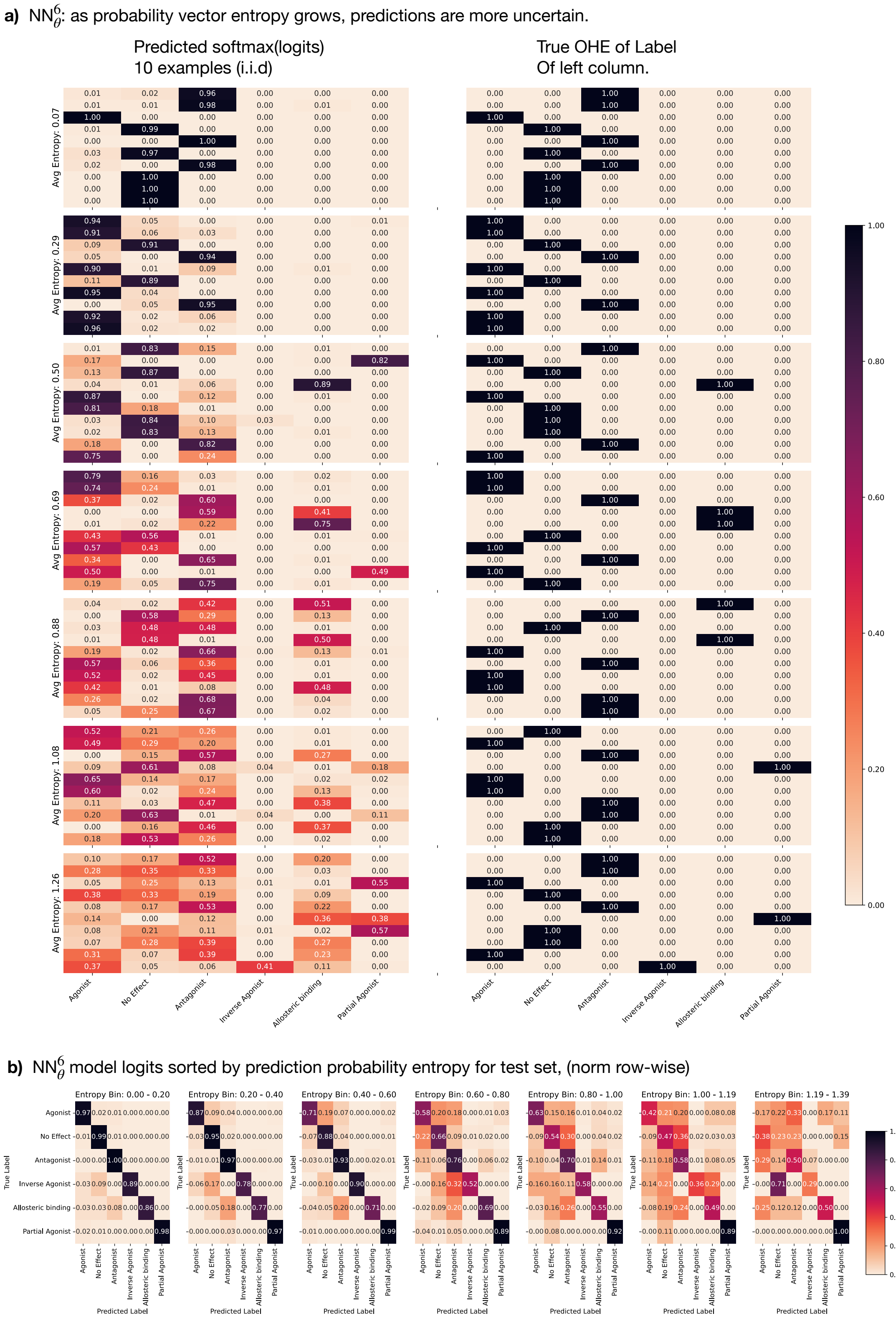


**Figure S4**: **Relating the Class Probability Entropy with the Predictive Performance of the Model**. **(a)** We calculate the class probability entropy (low if strongly peaked, high if dispersed) and sample 10 test examples i.i.d from each entropy bin. Top column shows that the one-hot encoding of the true labels (right) is strongly reflected in the predicted class labels (left, which are softmax(logits)). As the prediction entropy grows, the probability is spread across classes and looses it's correlation with the true one-hot encoding of the class label. **(b)** We compute confusion matrices on the test set, filtered by entropy bins to observe where probability is incorrectly assigned. For example, at 0.6-0.8 the antagonist and inverse agonist class is confused for 32% of predictions.

**a)** $NN_{\theta}^{6}$ model logits sorted by prediction probability entropy for test set (unnormalised)

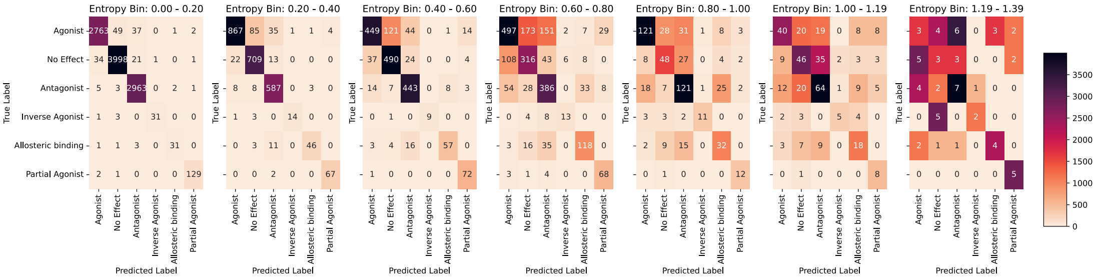


**b)** $NN_{\theta}^{6}$ predictions: AUC-ROC falls as model logits sorted by prediction probability entropy grow.

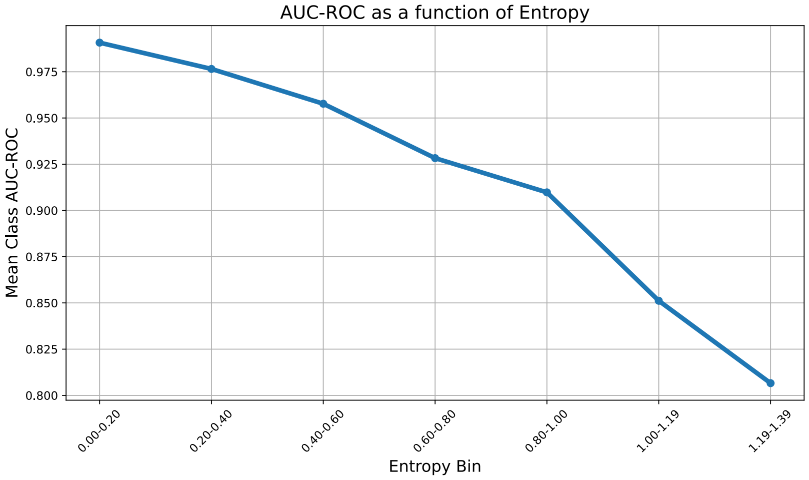


**Figure S5**: **Characterising the model performance filtering by prediction entropy shows well calibrated confidence**. **(a)** The same as Figure S4b but in unnormalised form. **(b)** The mean class AUC-ROC as a function of the sorted entropy bin shows a strong negative correlation between prediction entropy (non-peaked probability in class label) and the discriminative performance.

**a)** A single forward pass for all 128 receptors: $\text{NN}_{\theta}^{128}$, test metrics breakdown by class

| | | Precision | Recall | F1-Score | Support |
|---|---|---|---|---|---|
| Non-Threshold Predictions | **No Effect** | 0.8573 | 0.8265 | 0.8416 | 7221 |
| | **Allosteric binding** | 0.7514 | 0.7143 | 0.7324 | 385 |
| | **Antagonist** | 0.8552 | 0.8635 | 0.8593 | 6424 |
| | **Inverse Agonist** | 0.7632 | 0.4462 | 0.5631 | 130 |
| | **Partial Agonist** | 0.6905 | 0.3194 | 0.4367 | 454 |
| | **Agonist** | 0.8374 | 0.9201 | 0.8768 | 5205 |
| At optimal ROC-AUC Threshold | **No Effect** | 0.8123 | 0.8797 | 0.8446 | 7221 |
| | **Allosteric binding** | 0.3130 | 0.9351 | 0.4691 | 385 |
| | **Antagonist** | 0.8345 | 0.8812 | 0.8572 | 6424 |
| | **Inverse Agonist** | 0.0716 | 0.8538 | 0.1321 | 130 |
| | **Partial Agonist** | 0.1855 | 0.9009 | 0.3076 | 454 |
| | **Agonist** | 0.8006 | 0.9500 | 0.8690 | 5205 |

**b)** $\text{NN}_{\theta}^{128}$ ROC-AUC curve by class breakdown.

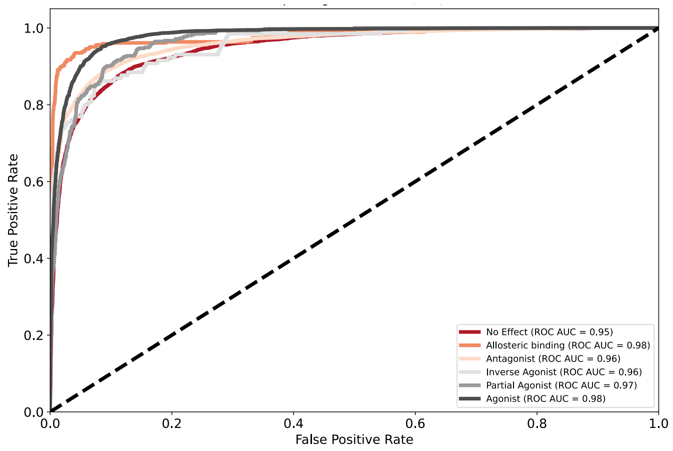


**c)** $\text{NN}_{\theta}^{128}$ performance is correlated with training class support.

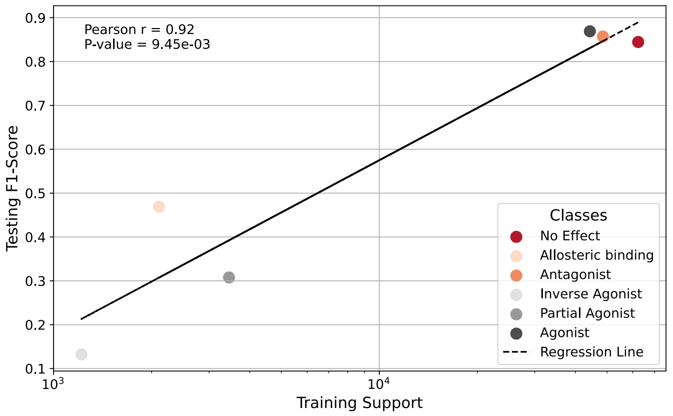


**Figure S6**: **Performance Metrics for the $\text{NN}_{\theta}^{1}28$ model**. **(a)** Class-wise breakdown of precision, recall, and F1-score. Due to the class imbalance (low support), at optimal ROC-AUC the F1-score falls significantly for the low support classes and recall increases. **(b)** ROC-AUC plot using the probability $p_i$ for the class label and the true label $y_i \in 0, 1$ shows strong descriminative performance. **(c) Testing F1-score correlates strongly with the training support.**

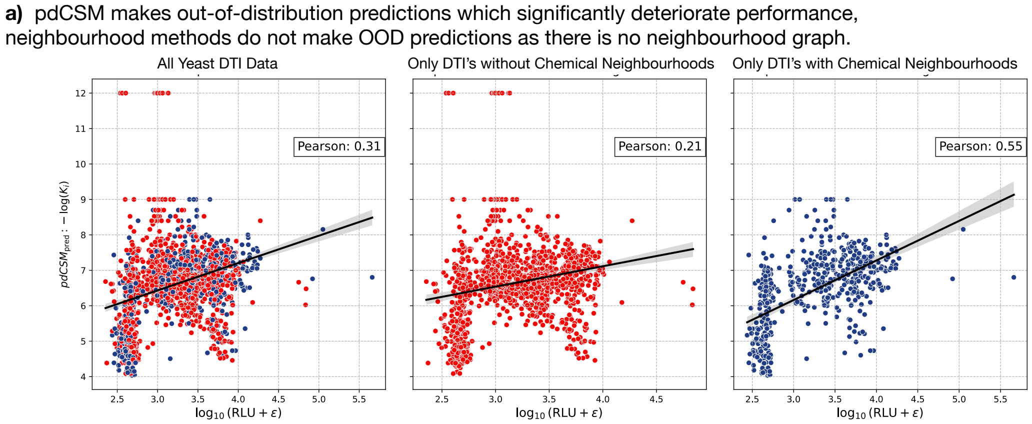


**Figure S7**: **pdCSM Bioactivity Predictions of Yeast Dataset Show OOD Behaviour**. For all investigated DTIs for which we have measured experimental RLU, we compare the correlation with the predicted $K_i$ (left) which shows low Pearson correlation and many predictions at 12, indicating OOD behaviour. 12 corresponds to a $K_i$ of picomolar, which is rare. Middle: Using our CSN we filtered for prediction for which there does not exists a chemical neighbourhood on a given DTI, this significantly reduces the correlation. Right: If we instead only pick DTIs for which there *do exist* chemical neighbourhoods, then the Pearson correlation markedly increases. This suggests that pdCSM makes out-of-distribution predictions that significantly deteriorate the performance of the model on unseen DTIs which are experimentally validated.

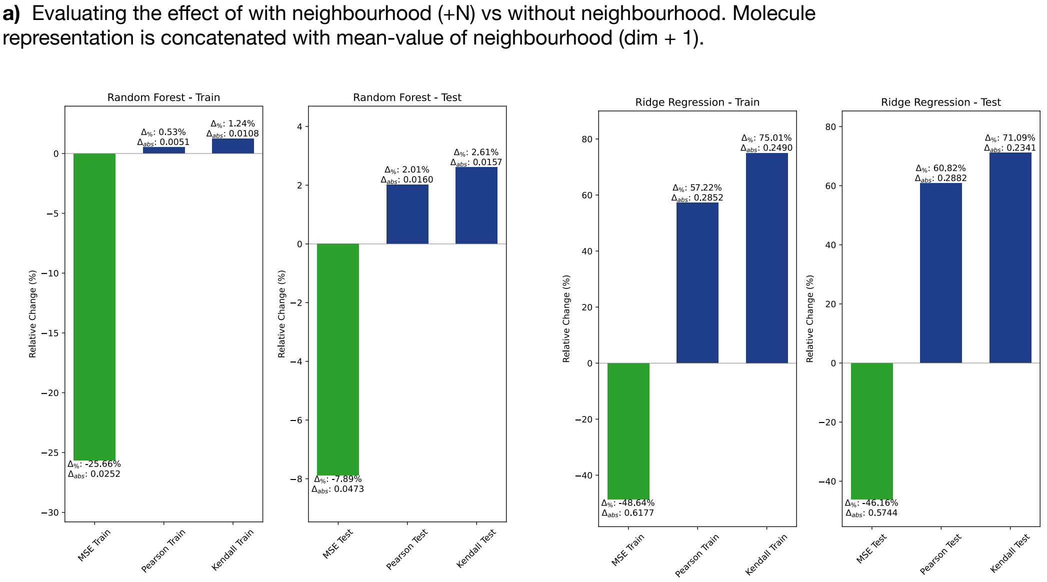


**Figure S8**: **Probing the LaF Effect**. Using two common ML models, we compare their relative performance on the training and testing test when *using* LaFs against not using LaFs. In the regression setting this corresponds to a change of input dimension by 1 (the mean value of the neighbourhood). For the higher-capacity RF model the increase in Pearson and Kendall correlation is smaller, but the inclusing of LaFs drive down the MSE significantly. In Figure 3d we provide a Table summarising the LaF effect on a low-capacity model like ridge regression. Here the results are illustrated, which shows that for both the training and testing set the MSE is reduced while the Pearson and Kendall correlation are increased significantly upon the inclusion of LaFs.

**a)** Graph Homophily: Mean-value predictions are good approximations. Per-receptor metrics reported.

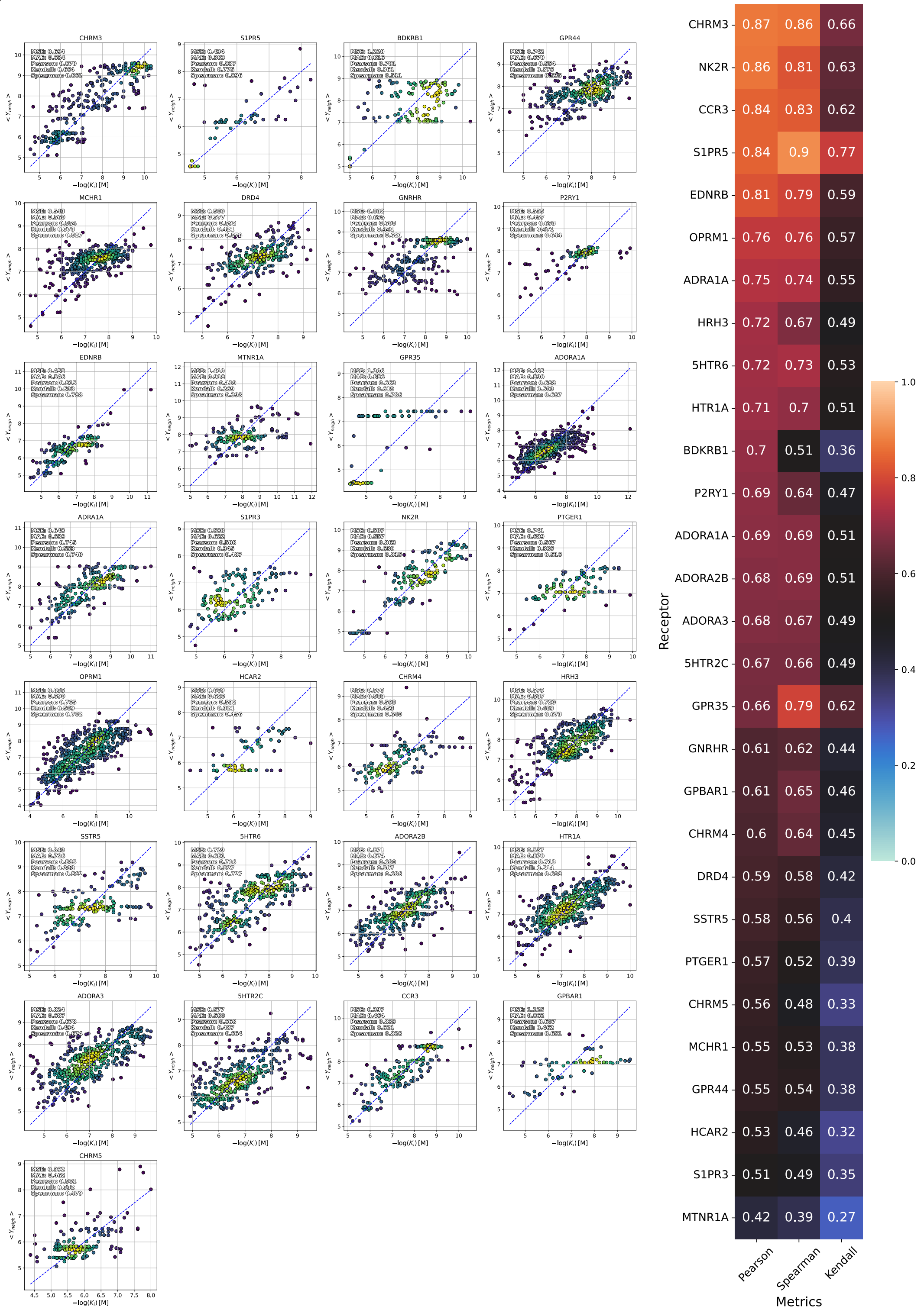


**Figure S9**: **Network Homophily as a Training Free Prediction**. If a network is homophilous, the mean value is a good approximation of the true value. In extension to Figure 3b,c,e and Table S1 we provide regression plots split by hGPCR to illustrate the performance of the mean value ($< Y_{neigh} >$ in detail). The heatmap (right) illustrates the Pearson, Spearman, and Kendall correlation coefficient under this training free prediction.

**a)** Graph Homophily: Comparing several aggregation techniques (mean, max, weighted, max similarity compound).

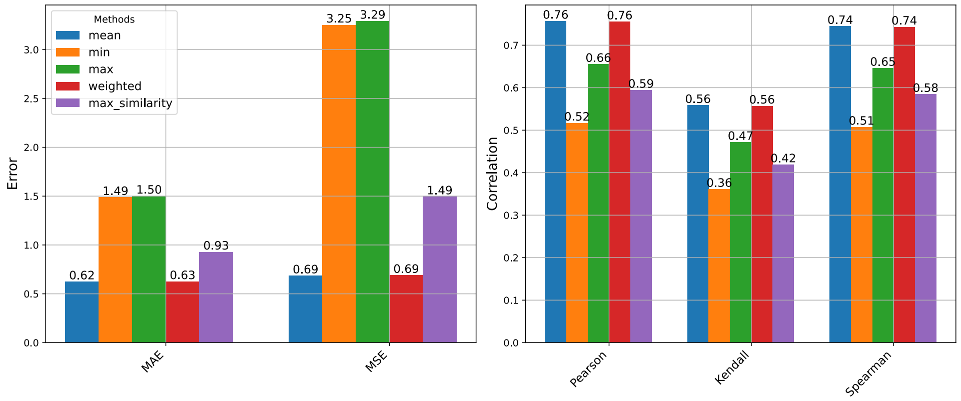


**b)** Graph Homophily: The test set regression plots excluding neighbourhoods by similarity.

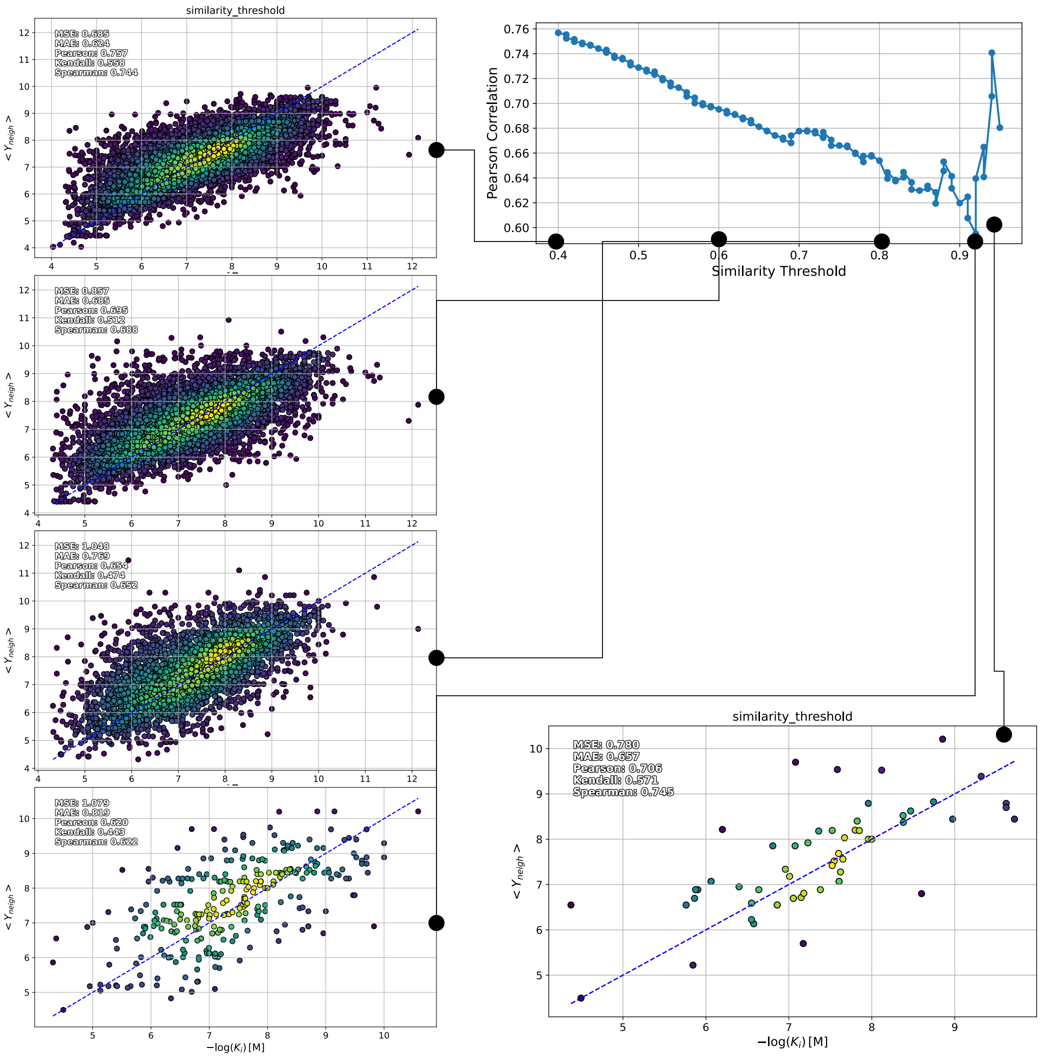


**Figure S10**: **Training Free Predictions, Aggregating Techniques**. **(a)** We find that the mean value offers a reliable training free prediction for the pdCSM test set. Min and max value, expectedly, are bad predictions. Surprisingly, taking only the most-similar molecule y-value (LaF), the prediction is also bad. **(b)** Pearson correlation filtered for the similarity threshold shows a striking negative correlation. Regression plots are shown at selected intervals, showing also the reduction in data-points. Curiously, keeping all neighbourhood data ensures the best prediction.

**a)** Train/Test metrics for MSE-loss objective on pdCSM data (CSNN). Fixed RDKit representation.

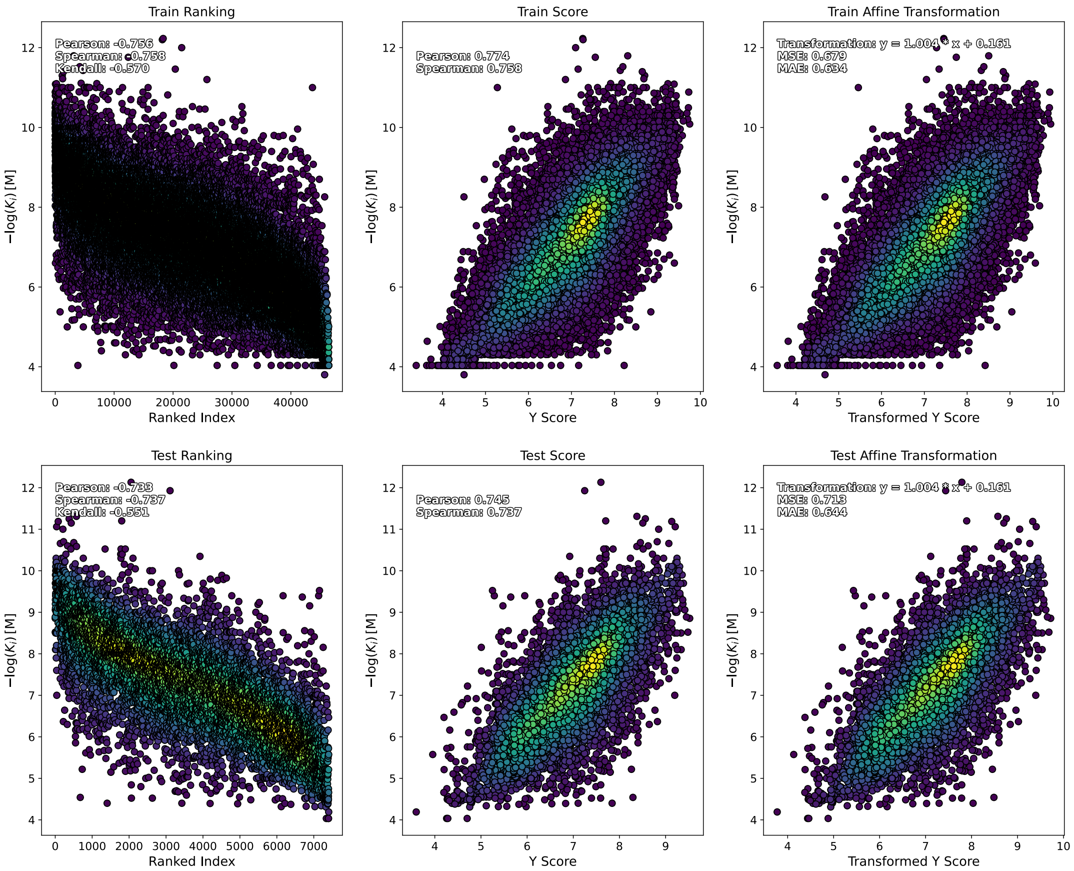


**Figure S11**: Train/Test metrics for the pdCSM dataset using the CSNN model, a MSE-loss, and fixed RDkit compound representations (copying those from the original paper, without feature selection).

**a)** Training curves for BT-loss objective.

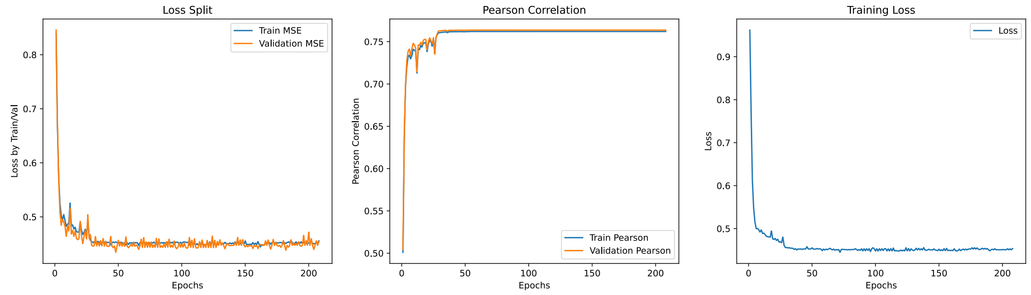


**BT-loss - fixed**

**b)** Train/Test metrics for BT-loss objective on pdCSM data (CSNN). Fixed RDKit representation.

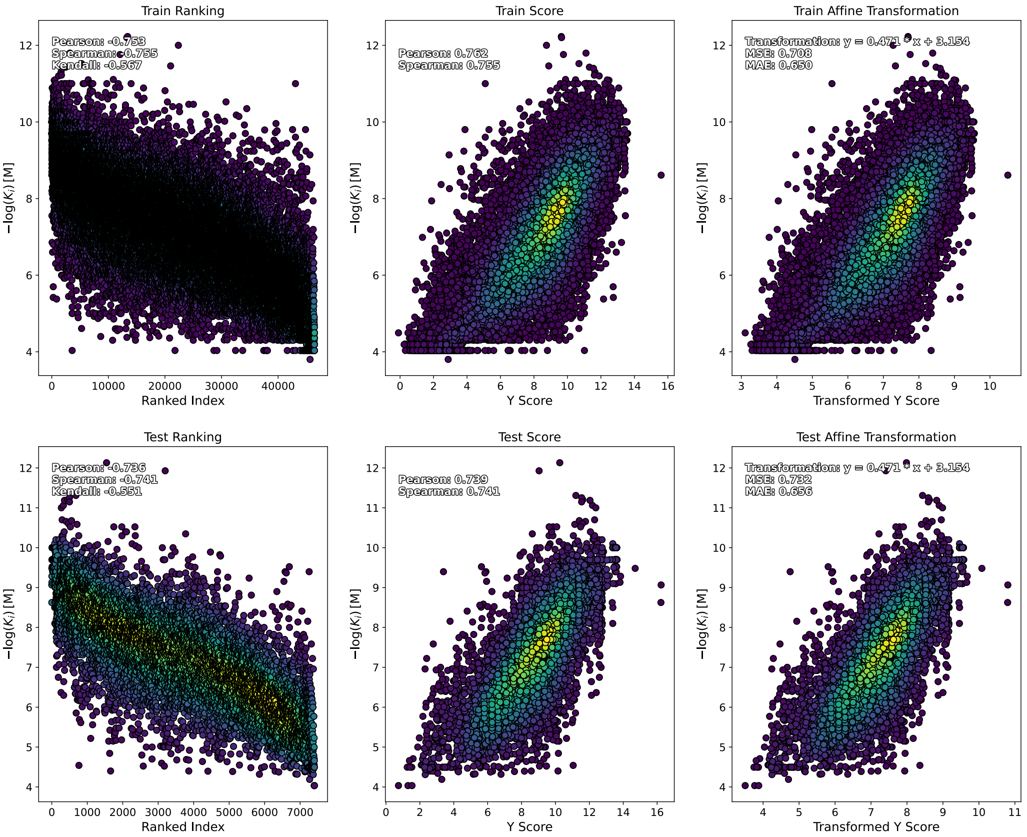


**Figure S12**: Train/Test metrics for the pdCSM dataset using the CSNN model, a BT-loss (contrastive), and fixed RDkit compound representations (copying those from the original paper, without feature selection). Due to the BT-loss not learning the true y-label an affine transformation is fit on the training data and used to align the predicted scores with the true scores for error metric calculation. The same affine transformation is performed on the test set. BT-loss models learn a ranking function rather than a regression objective, which up to an affine transformation, is equivalent to the true y-label.

**a)** Train/Test metrics for MSE-loss objective on pdCSM data (RF + N). Fixed RDKit representation.

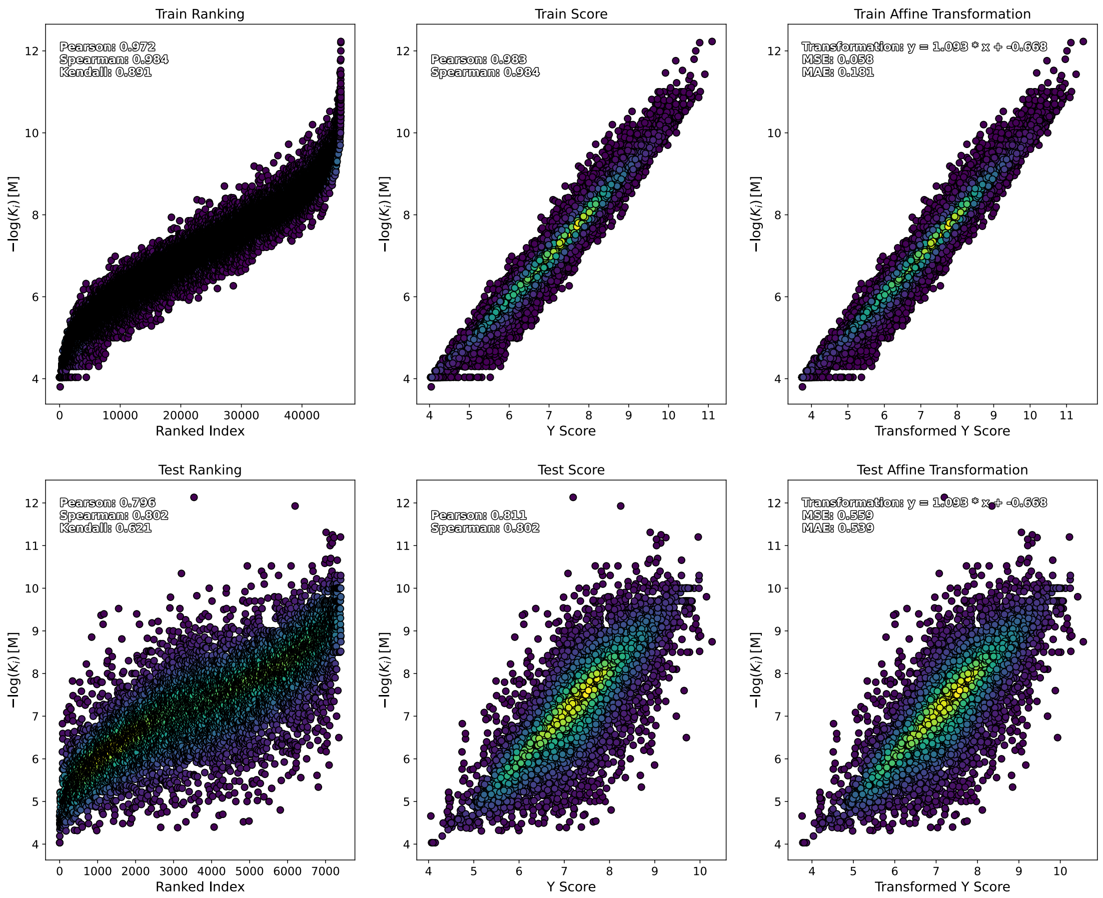


**Figure S13**: Train/Test metrics for the pdCSM dataset using the RF + N model, a MSE-loss, and fixed RDkit compound representations (copying those from the original paper, without feature selection). Interestingly, Figure S11, S12 show no signs of over-fitting for the CSNN models (similar Pearson on training/testing data), while the RF + N model looks to be partially over-fitted (although with best test metrics).

**a)** Train/Test metrics for MSE-loss objective on pdCSM data (RF + N). Fixed RDKit representation. Only **HTR1A** data shown.

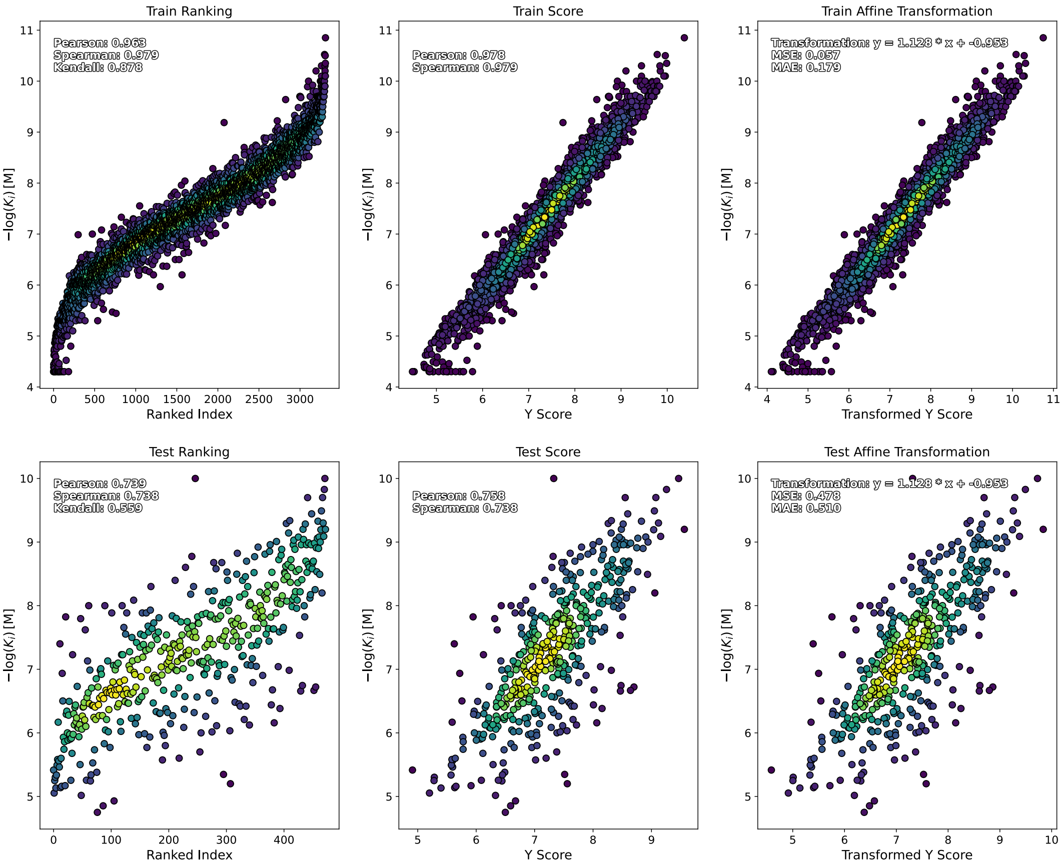


**Figure S14**: Train/Test metrics for the pdCSM dataset using the RF + N model, a MSE-loss, and fixed RDkit compound representations (copying those from the original paper, without feature selection). The model is trained on all data and only training and testing data points for DTIs involving the HTR1A hGPCR is shown.

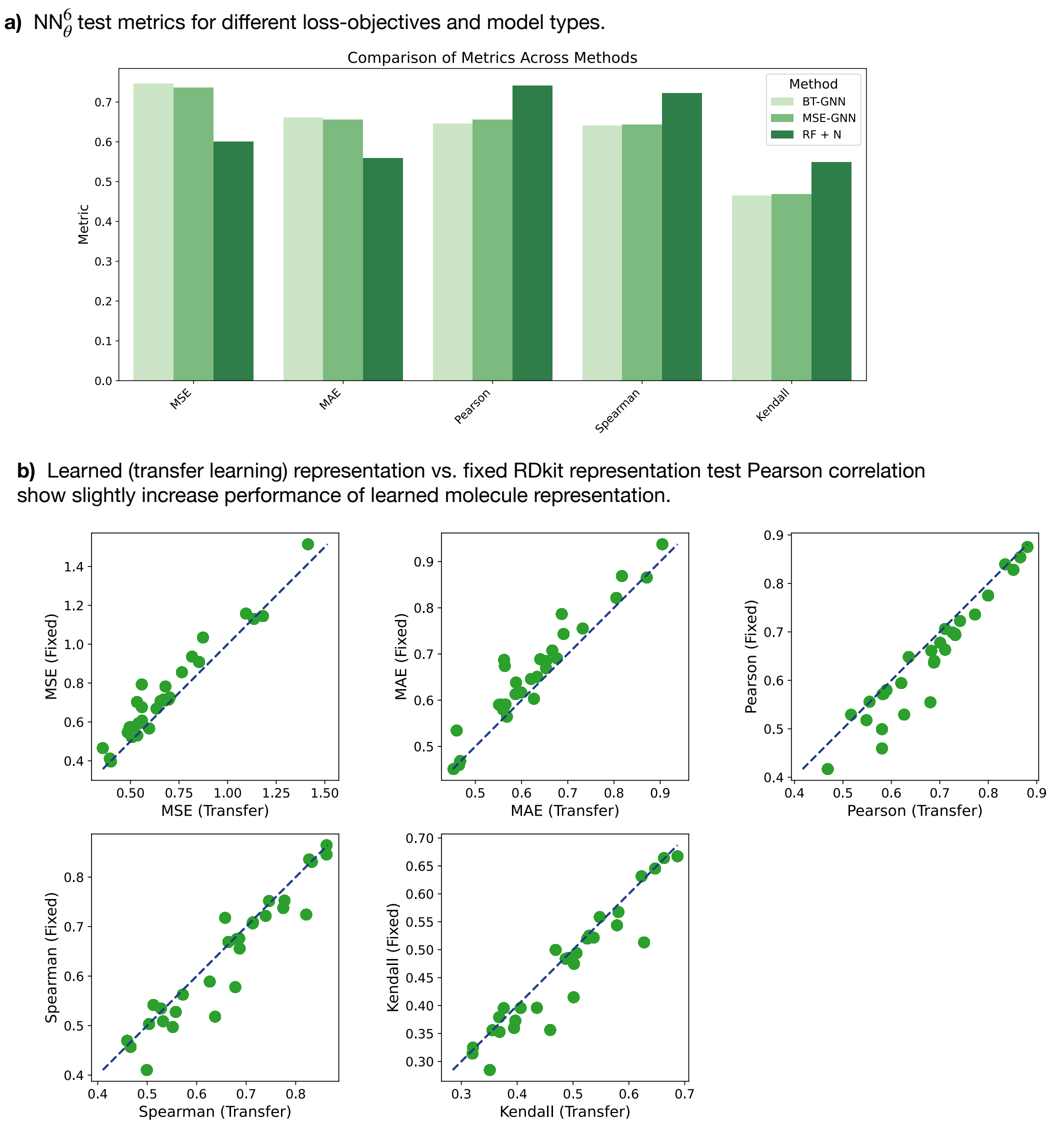


**Figure S15**: **(a)** A comparison of test metrics between ML models on the pdCSM dataset show that the RF + N model outperforms the others. **(b)** We compare the node-embedding (compound representation) for the same model (size and training objective, training epochs), which shows that learned representations (obtained from ChemProp) slightly outperform the fixed RDKit representation.

**a)** Train/Test metrics for RF + N on pdCSM data. Fixed RDKit representation. Only **CHRM3** data shown.

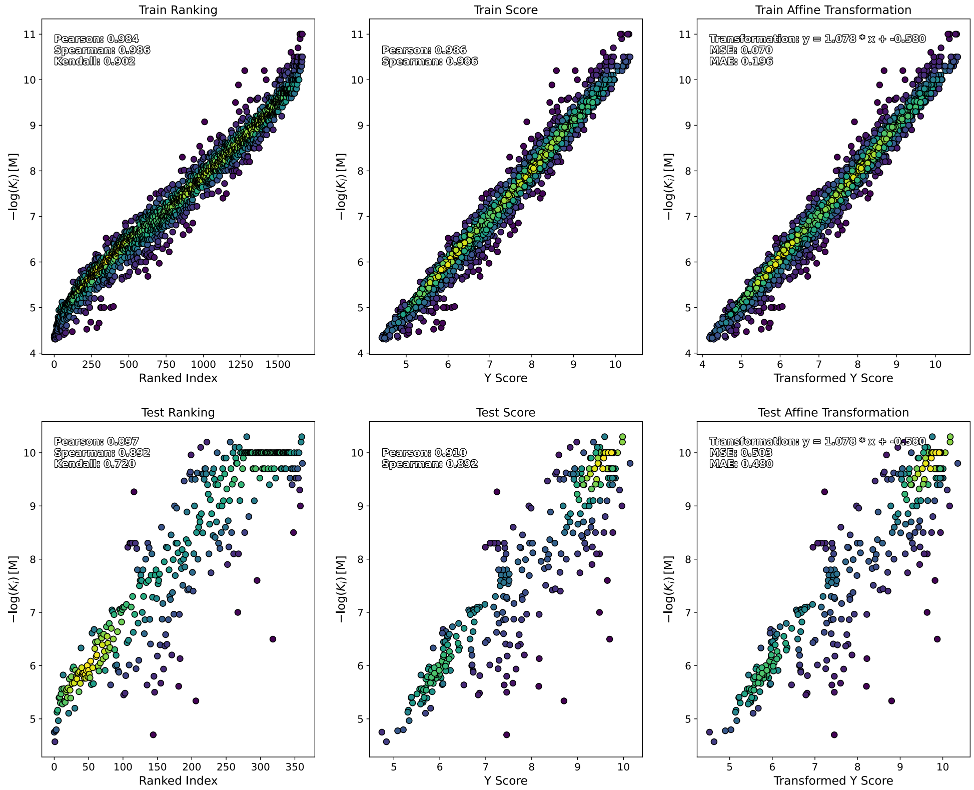


**b)** Train/Test metrics for MSE-loss objective on pdCSM data (CSNN). Fixed RDKit representation. Trained only on **CHRM3** data shown (rather than all data with receptor representation, above). High variance across epochs, lower test metrics.

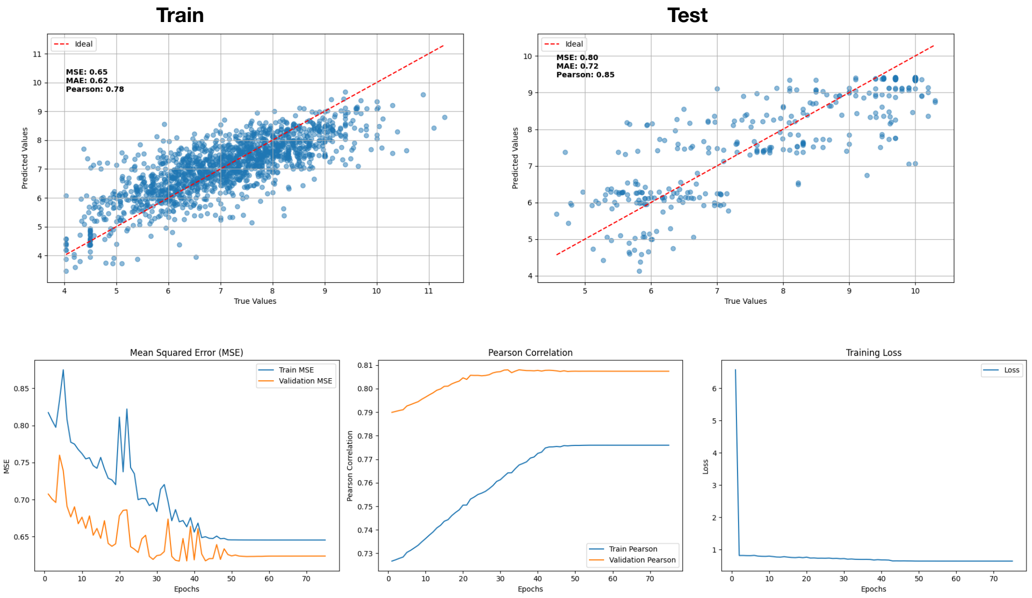


**Figure S16**: **Comparing one-target-one-model vs One-model-for-all**. The original pdCSM method (*17*) trains a RF model per hGPCR (thus avoiding an explicit target representation). We compare the settings (a) and (b) and found that training was unstable due to a high-variance (small batches and little data). A single model which handles all data, necessarily representing both the compound and target representations was found to be the most efficient solution.

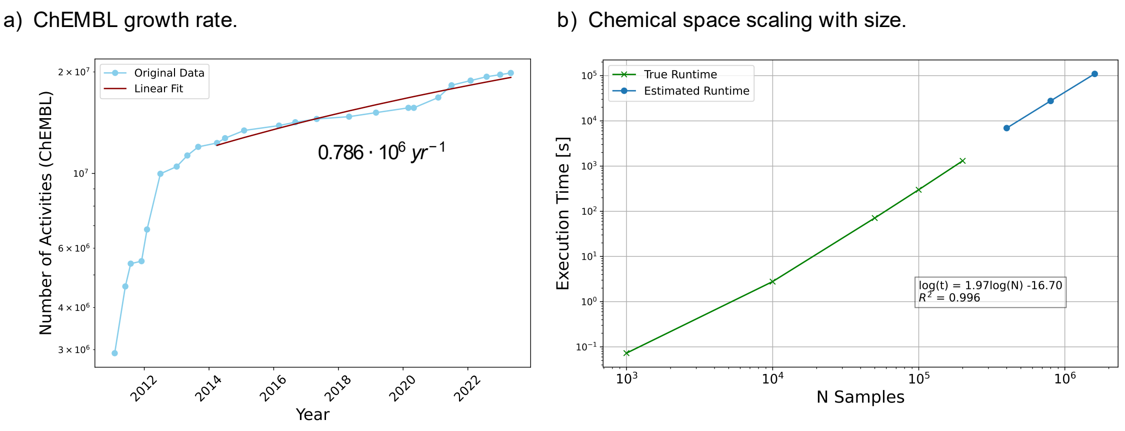


**Figure S17**: **(a)** ChEMBL growth rate per year estimated using data from (*79*), which documents activites for each ChEMBL release. **(b)** Chemical space network construction computation time, as calculated on a home computer (MacBook, Apple M1 Max, 32 GB RAM). For tens of thousands of compounds, the execution time is on the order of seconds; for hundreds of thousands, it is on the order of minutes. For compound libraries larger than 1 million compounds the runtime was estimated, but could not be executed due to lacking memory. Although, looking up the 1-hop neighbourhood for a single compound is extremely fast for a 1.6 million compound library 1.55 ± 0.05 seconds (using randomly samples ZINC smiles strings with precomputed fingerprints). For less than 20K compounds, a batch all-vs-all CSN can be computed in less than 3.9 seconds. For larger libraries a one-vs-all calculation is performed for all rows (upper-triangular).

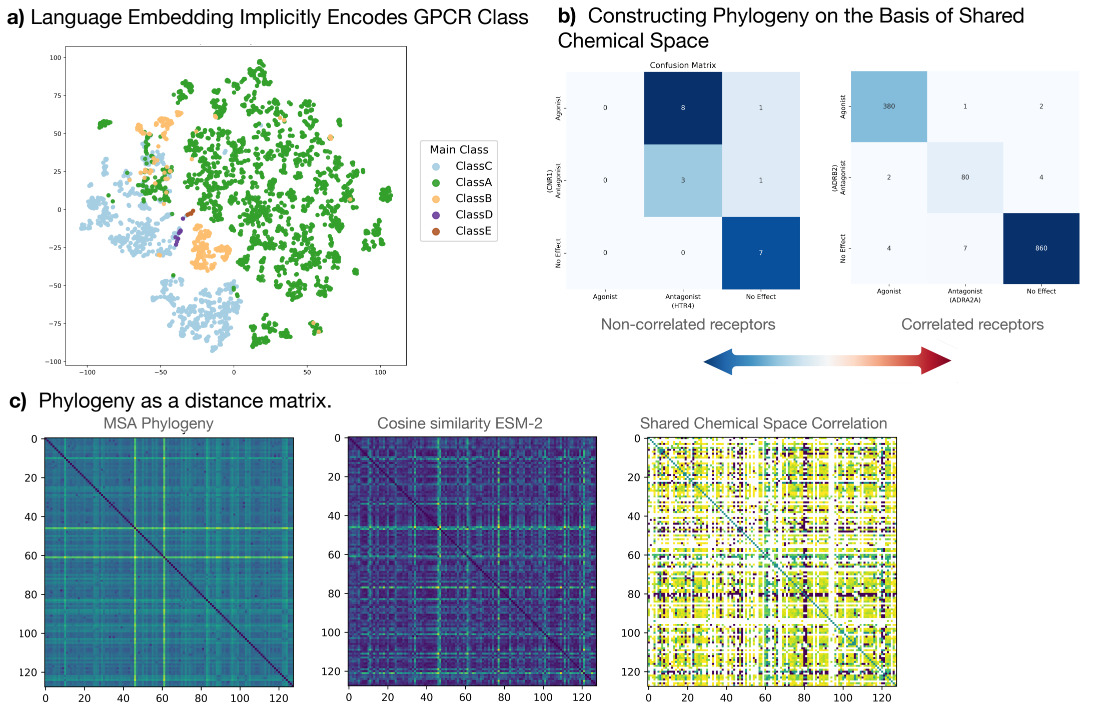


**Figure S18**: **(a)** Protein Language model (ESM-2) embedding of hGPCR using the ground truth labels from the GDS dataset (*80*), which is comprised of 8354 unique GPCRs. Embedding were dimensionality reduced using t-SNE (*81*). **(b)** Our unique labeled dataset allows for the identification of correlated hGPCRs on the basis of a shared chemical space. For example, ADRA2A and ADRB2 are known to be related phylogenetically by MSA, which is reflected in the populated diagonal, implying that a compound that is an agonist on one hGPCR is likely also to be on the other. For less correlated hGPCRs, the chemical space will be less shared (off-diagonal entries), from which a metric can be derived using Cramer's V to obtain a correlation between categorical variables. **(c)** The output of all three methods (i) MSA, (ii) ESM-2 embeddings, and (iii) shared chemical space are a distance matrix from which the phylogenetic trees in S19 are derived.

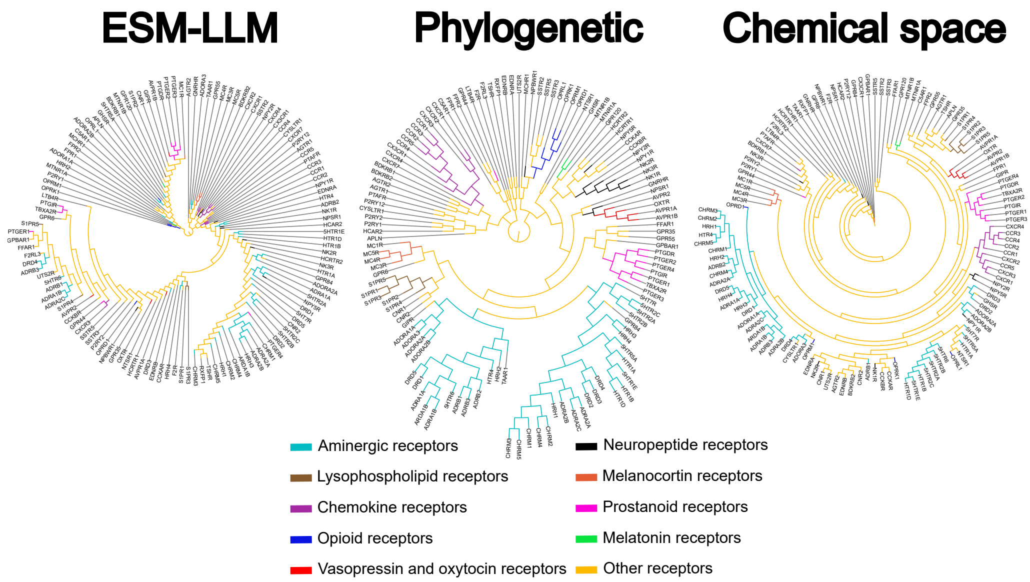


**Figure S19**: hGPCR clustering showing tree structures generated from distances calculated by the ESM-2 Large Language model (ESM-LLM), multiple sequence alignment (MUSCLE), and from the chemicals they are able/unable to sense. Phylogenetic trees are based on distance matrices (See Figure S18)

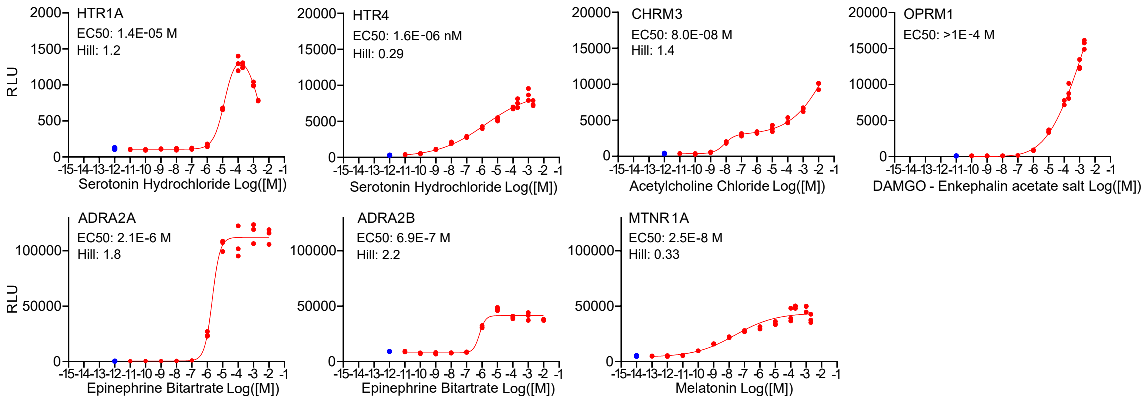


**Figure S20**: Dose responses of 7 chosen hGPCRs with known agonists. Red dots are individual replicates at each concentration (n=3), blue dots are 0 M negative controls (n=3), not included in the generation of the DRCs. All DRCs are generated using built-in functions in GraphPad Prism 9.5.0. For OPRM1 The EC50 is given as a lower bound, 50% of the maximum observed RLU of this hGPCR as we were unable to plateau reporter output at high concentrations. hGPCR independent effects were determined using the parent strains (ScFH236 & 237)(Supplementary Figure S21).

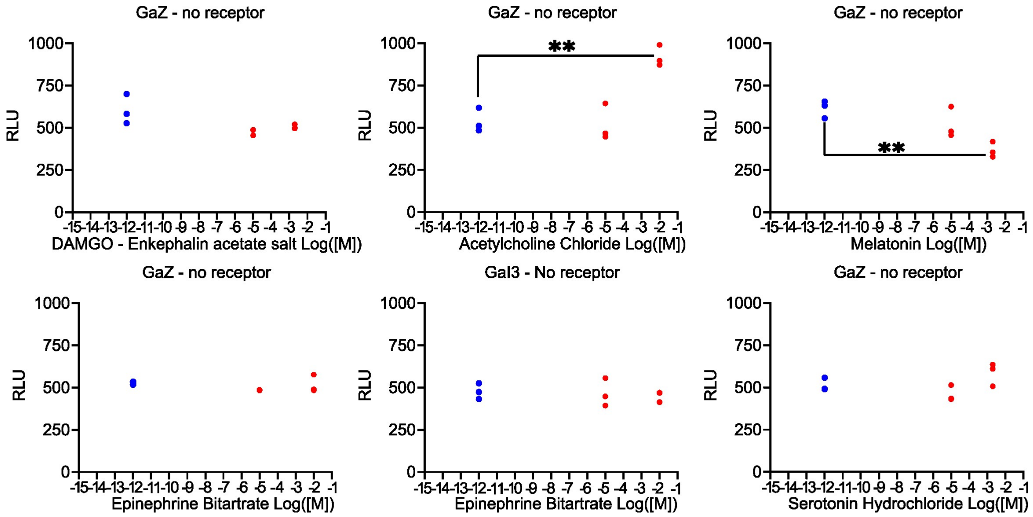


**Figure S21**: **No-hGPCR controls for Figure 2**, Blue dots are 0-concentration controls placed artificially on the logarithmic scaled x-axis. Red dots represent individual datapoints (N=3 at all tested concentrations) of no-hGPCR control strains tested against concentrations of the ligands used in Figure 4d and S20. hGPCR independent effects are observed for acethylcholine chloride at 10 mM and for melatonin at 2 mM (Supplementary figure S21). The early response of acetylcholine chloride against CHRM3 (1E-10 – 1E-5 M) plateaus before the unspecific effect begins, allowing us to determine an EC50 and Hill coefficient for the specific effect on the hGPCR. For MTNR1A, the receptor unspecific effect of melatonin indicates that MTNR1A mediated signalling might not have reached a true plateau. **p<0.005

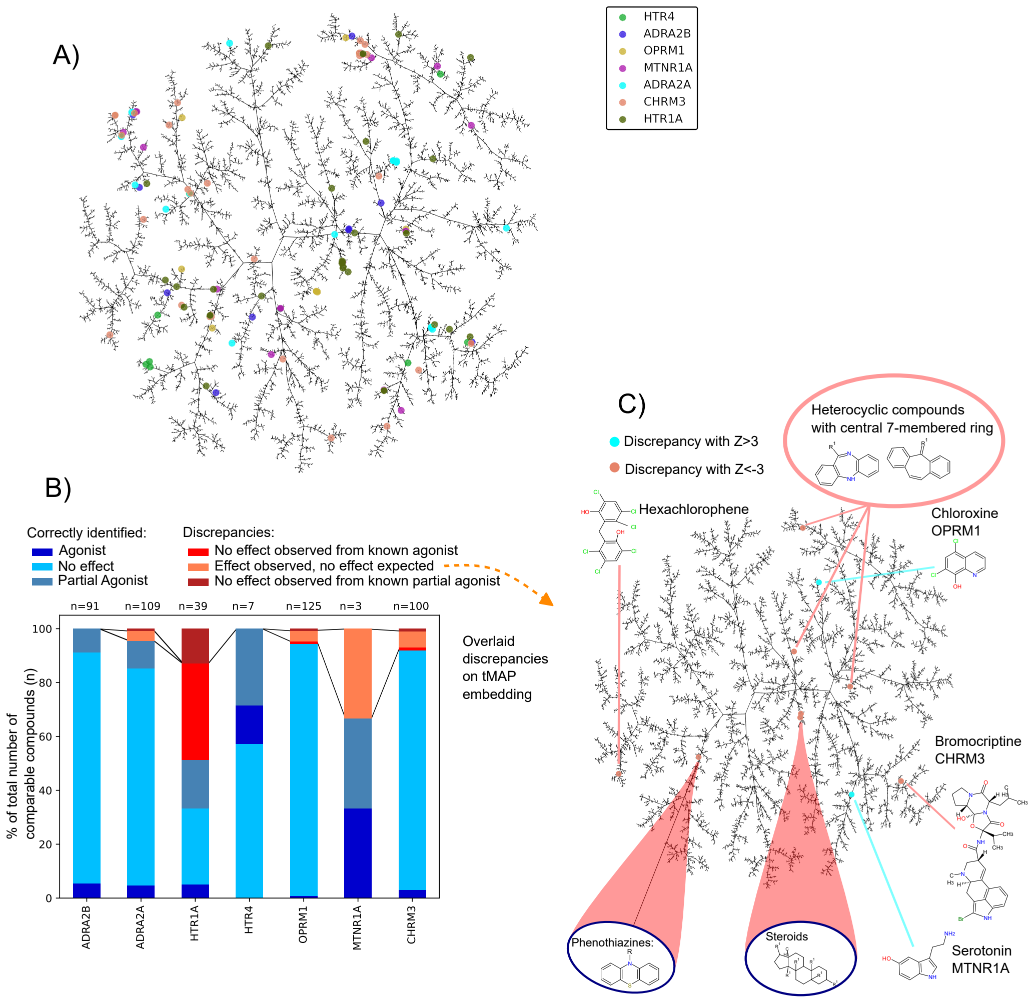


**Figure S22**: tMAP embedding (*82*), generated on the dataset of chemicals targeting one or more of the 128 investigated GPCRs. A) The hits obtained from the tested chemical library is highlighted colored by which GPCR the hit was observed on. B) hit discrepancies identified by 1:1 comparison with prior art. C) Discrepancies, where no effect was expected, but an effect was observed, was overlaid on the tMAP of the chemical space. Some discrepancies clustered together.

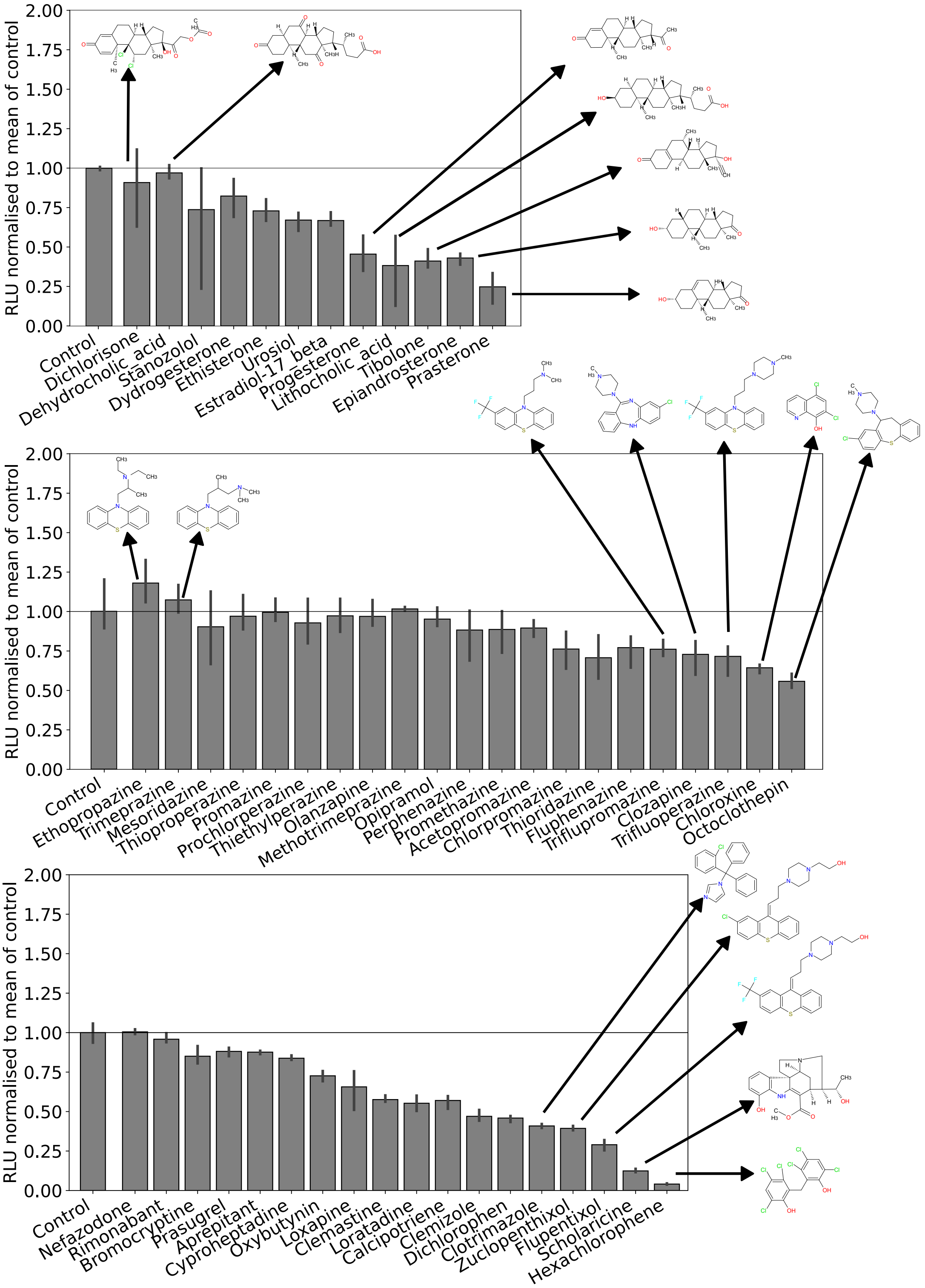


**Figure S23**: GPCR independent effects on yeast, assessed as RLU normalised to the mean RLU of negative controls (1% DMSO) n=3. Top panel contains measurements of steroid class compounds, showing that more oxidised steroids like Dichlorisone and dehydrocholic acid do not affect the signal dampening in the same way as more reduced steroids. Middle panel shows measurements for phenothiazines and similar compounds, illustrating that especially halogenated phenothiazines and 7-membered ring containing compounds dampen the signal. Chloroxine was also included here, as it dampened the signal on non-OPRM1 receptors. Bottom panel contains more tri-cyclic compounds aswell as other compounds selected due to observations of generally dampening effects on luminescence. Compound structures rendered for the 5 most inhibiting compounds in each plot, and for other compounds of interest. Any compound where the standard deviation does not reach the mean of the controls is removed from further analysis due to bias from receptor independent effects. Controls in each plot represent separate triplicates of measurements on RLU upon exposure to 1% DMSO, which is present in all samples. N=3 for all bars.

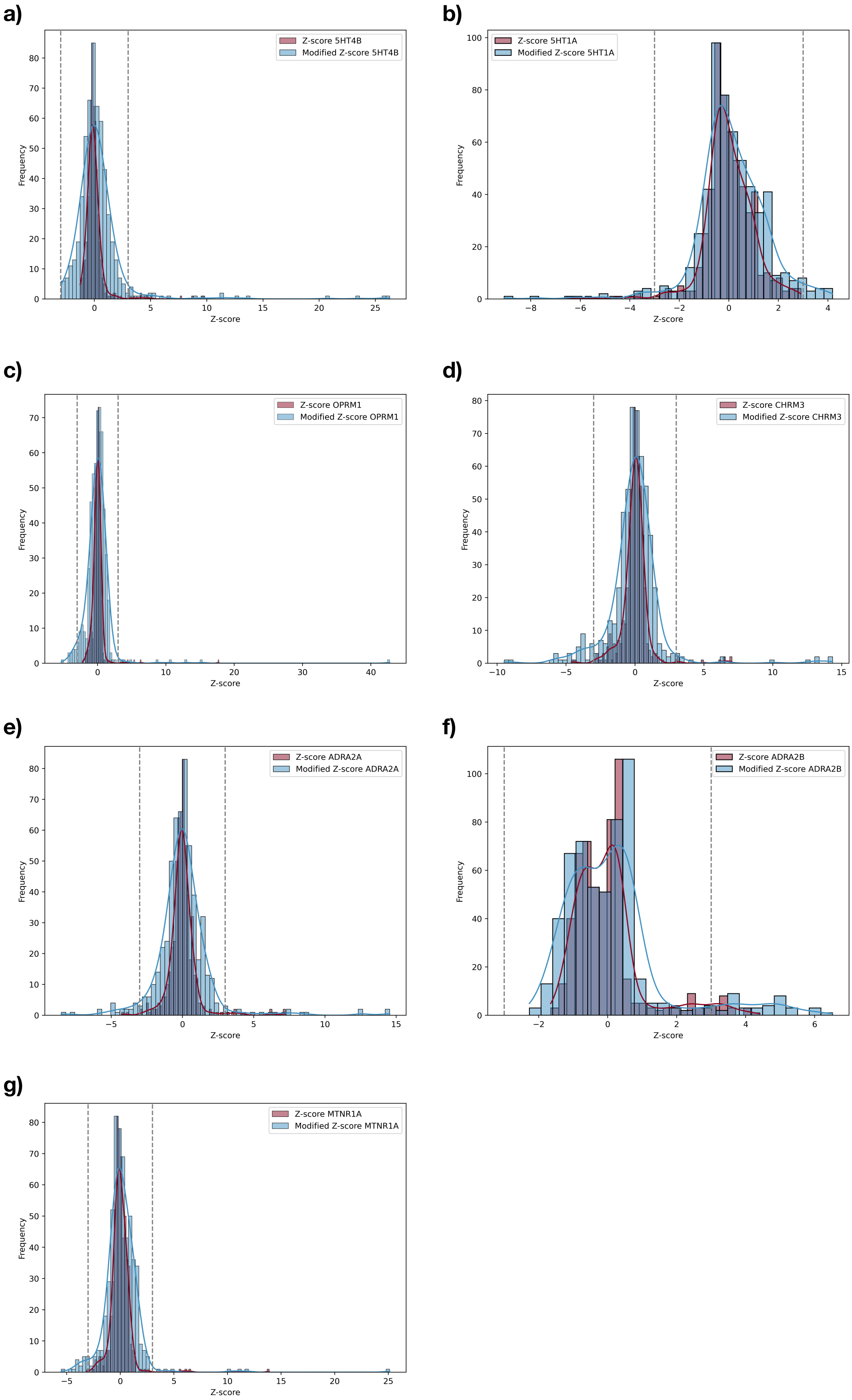


**Figure S24**: **Effect of Modified Z-score on recovering hits**, data is shown for each hGPCR and the histogram fit is plotted using kernel density estimation to contrast the effect of the Z-score (Eq. 9) against the modified Z-score (Eq. 10).

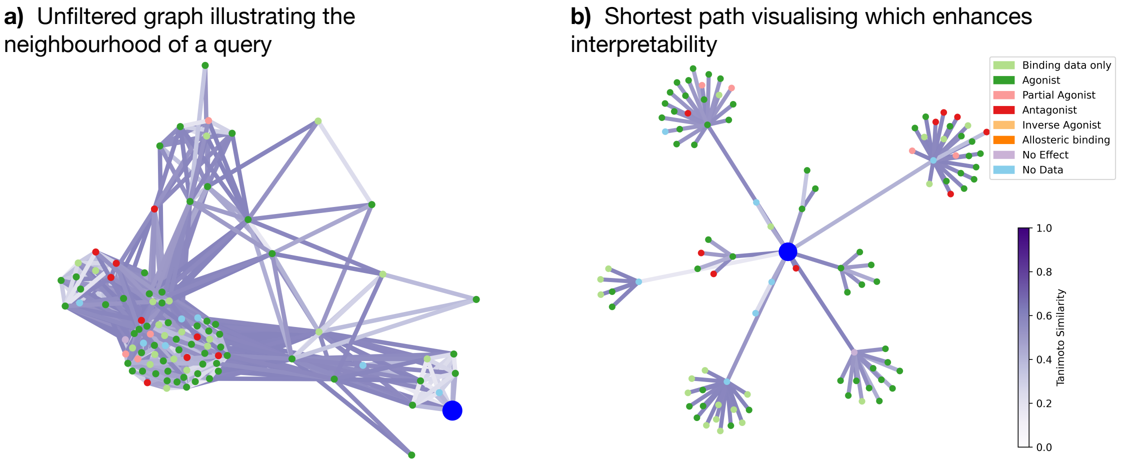


**Figure S25**: Enhancing interpretability of graph neighbourhoods for a given query by considering only the shortest paths from the query to all other codes. This converts the graph into a tree graph that all expand radially from the query.

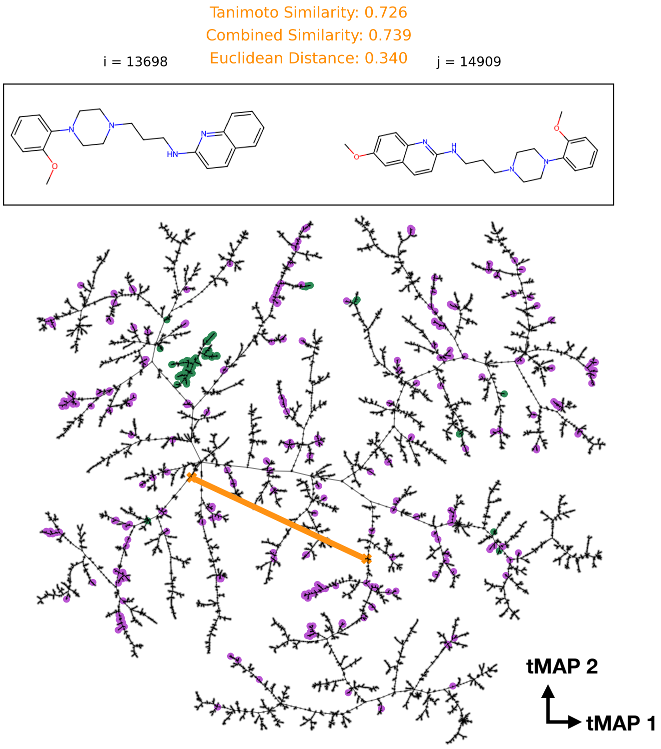


**Figure S26**: To avoid a graph structured chemical space an embedding space can be used to efficiently find k-nearest neighbours. We tried using the tMAP (*82*) projection, which we use for visualising chemical space, and which is developed for this chemical use case, but found that molecules which have a high Tanimoto similarity do not necessarily lie close in the embedding space. This makes finding k-NN problematic and as such a all-vs-all pairs similarity search was carried out.

**a)** Extended Table for Novel Hits.

| Name | Use [Approved] | Z-score [novel] | Predicted Label | Predicted -log10(Ki) | Z-score [confirmed] |
|---|---|---|---|---|---|
| Tinoridine | Against pain and inflammation | HTR4: 5.87 | OPRM1: No Effect<br>CHRM3: No Effect | OPRM1: 4.113<br>CHRM3: 6.630 | OPRM1: 0.068<br>CHRM3: -0.553 |
| Piribidil | Parkinsons disease | ADRA2B: 4.68 | OPRM1: No Effect<br>HTR1A: Antagonist<br>HTR1A (argmax): Partial Agonist | OPRM: 4.042<br>HTR1A: 5.656 | OPRM1: 0.336<br>HTR1A: 0.747 |

**Figure S27**: Extended Table for Figure 4g, which summarises also DTIs for which predictions were produced (data in the chemical neighbourhood was available). We see that "No Effect" label predictions are reflected in the non-statically significant Z-score. Piribidil has a label disagreement from the argmax prediction and the $\mathrm{NN}_{\theta}^{6}$ model and shows a non-statistically significant Z-score as well as low predicted $K_i$ values (from RF + N model).

#### S10 Supplementary Tables

**Table S1**: Performance metrics under the homophilly assumption for pdCSM dataset regression task: the mean value over the chemical neighbourhood is a good approximation of the true value. Sorted in order of descending Pearson correlation coefficient.

| Method | hGPCR | MSE | MAE | Pearson | Kendall | Spearman |
|---|---|---|---|---|---|---|
| mean | CHRM3 | 0.694 | 0.634 | 0.87 | 0.664 | 0.862 |
| mean | NK2R | 0.507 | 0.557 | 0.863 | 0.63 | 0.815 |
| mean | CCR3 | 0.397 | 0.464 | 0.839 | 0.621 | 0.828 |
| mean | S1PR5 | 0.434 | 0.383 | 0.837 | 0.775 | 0.896 |
| mean | EDNRB | 0.455 | 0.546 | 0.815 | 0.593 | 0.788 |
| mean | OPRM1 | 0.835 | 0.69 | 0.765 | 0.569 | 0.762 |
| mean | ADRA1A | 0.648 | 0.639 | 0.745 | 0.553 | 0.74 |
| mean | HRH3 | 0.579 | 0.587 | 0.72 | 0.489 | 0.673 |
| mean | 5HTR6 | 0.729 | 0.651 | 0.716 | 0.527 | 0.727 |
| mean | HTR1A | 0.537 | 0.57 | 0.713 | 0.514 | 0.698 |
| mean | BDKRB1 | 1.22 | 0.816 | 0.701 | 0.361 | 0.511 |
| mean | P2RY1 | 0.535 | 0.457 | 0.693 | 0.471 | 0.644 |
| mean | ADORA1A | 0.665 | 0.59 | 0.688 | 0.509 | 0.687 |
| mean | ADORA2B | 0.571 | 0.574 | 0.68 | 0.507 | 0.686 |
| mean | ADORA3 | 0.824 | 0.687 | 0.678 | 0.494 | 0.674 |
| mean | 5HTR2C | 0.577 | 0.58 | 0.668 | 0.487 | 0.664 |
| mean | GPR35 | 1.306 | 0.856 | 0.663 | 0.619 | 0.786 |
| mean | GNRHR | 0.882 | 0.695 | 0.608 | 0.441 | 0.621 |
| mean | GPBAR1 | 1.125 | 0.862 | 0.607 | 0.462 | 0.651 |
| mean | CHRM4 | 0.573 | 0.583 | 0.598 | 0.453 | 0.64 |
| mean | DRD4 | 0.56 | 0.577 | 0.592 | 0.421 | 0.578 |
| mean | SSTR5 | 0.849 | 0.726 | 0.585 | 0.398 | 0.562 |
| mean | PTGER1 | 0.741 | 0.689 | 0.567 | 0.386 | 0.516 |
| mean | CHRM5 | 0.392 | 0.462 | 0.561 | 0.332 | 0.479 |
| mean | MCHR1 | 0.543 | 0.56 | 0.554 | 0.378 | 0.527 |
| mean | GPR44 | 0.742 | 0.67 | 0.554 | 0.376 | 0.535 |
| mean | HCAR2 | 0.669 | 0.626 | 0.532 | 0.321 | 0.456 |
| mean | S1PR3 | 0.58 | 0.622 | 0.508 | 0.345 | 0.487 |
| mean | MTNR1A | 1.41 | 0.918 | 0.419 | 0.269 | 0.393 |

**Table S2**: Tasks performed by state of the art predictive algorithms for DTI predictions

| Input | Task | Explanation | Notation | Type | Refs |
|---|---|---|---|---|---|
| *ci* | *yi* | Does the compound have GPCR activity? | *P(yi∣ci)* | Regression, Classification | (*6*, *83*)) |
| *ci,tj* | *yi* | Given a compound and its target representation, is the compound active? | *P(yi∣ci,tj)* | Regression, Classification | |
| *ci,cj* | *yi,yj* | Given two compounds, how active are they? | *P(yi,yj∣ci,cj)* | Regression, Classification | (*39*) |
| *ci* | *tj* | Given a compound, predict its target. | *P(ti∣ci)* | Classification | (*15*) |
| *ci* | | | | | (*84*) |
| Inference from Neighbourhood | *aj* | Given a compound, predict adverse drug events. | *P(ai∣ci)* | Regression, Classification | |
| *ci,tj,Ni(yj)* | *yi* | Given a compound, it's neighbourhood activity on one target j, and a target representation tj, is it active? | *P(yi∣ci,Ni(yj),tj)* | Classification | (ours) |
| *ci,Ni(yn),tn* | *yi,n* | Given a compound, it's neighbourhood activity on all n targets, and target representations, on which target is ci active? | *P(yn,i∣ci,Ni(yn),tj)* | Classification | (ours) |

| **Class** | **Precision** | **Recall** | **F1-Score** | **Support** |
|---|---|---|---|---|
| Agonist | 0.9256 | 0.8746 | 0.8994 | 61835 |
| No Effect | 0.9120 | 0.9312 | 0.9215 | 57575 |
| Antagonist | 0.8940 | 0.9338 | 0.9135 | 51418 |
| Inverse Agonist | 0.7745 | 0.7058 | 0.7386 | 1499 |
| Allosteric Binding | 0.7046 | 0.6621 | 0.6827 | 4208 |
| Partial Agonist | 0.8179 | 0.9119 | 0.8623 | 2621 |
| Accuracy | | | 0.9039 | 179156 |
| Macro Avg | 0.8381 | 0.8365 | 0.8363 | 179156 |
| Weighted Avg | 0.9042 | 0.9039 | 0.9036 | 179156 |

**Table S3**: Argmax(.) performance metrics across all 128 hGPCR. Metrics are shown per bioactivity class, and the data illustrates the remarkable predictability of a compound given its neighborhood.

**Table S4**: The 7 hGPCRs investigated in vitro in this study, and the known treatable conditions for which the respective targets have been used clinically.

| hGPCR Target | Known target indications |
|---|---|
| HTR1A | Anxiety(*85*)<br>Depression(*85*)<br>Pain(*85*)<br>Schizophrenia(*85*)<br>Neuroprotection(*85*) |
| HTR4 | Gastro-intestinal disorders(*28*)<br>Anxiety(*86*) |
| CHRM3 | Mucous membrane dryness (*87*)<br>Blood pressure regulation(*30*)<br>Alzheimers disease(*29*)<br>Schizophrenia(*29*) |
| OPRM1 | Pain(*88*)<br>Addiction(*88*) |
| MTNR1A | Sleep–Wake Disorders(*31*) |
| ADRA2A & ADRA2B | Pain(*89*)<br>Peri-operative ischaemia(*89*)<br>Blood pressure regulation(*89*) |

Table S5: **a.** Prices estimated for the costs of performing an assay similar to the one described in this paper in commercially available GPCR screening systems. **b.** We used a low volume 96wp format for our small screen of 4000 compounds, converting to 384 wp format as demonstrated by Miettinen et al 2022.(*19*) will reduce the cost of plates by up to 55%, lowering the total cost pr. well to 0.91USD. **c.** Not all suppliers state their prices outside confidential quotes, this list only contains suppliers with listed prices. Other suppliers include Multispan, Molecular Devices, and Innoprot. **d.** For the costs of multiplexing, the prices estimated in the respective publications were used, these values reflect the price pr. well, where a single well can assay more than 300 GPCRs. The supporting calculations are available in supplementary file S1. The prices listed are for the operational costs only, excluding instrument acquisition, overhead, and salary. Other sources estimate costs around 10 times higher for mammalian cell based assays(*98*)

| Assay | Advantages | Limitations | Costs: 4000 tests across 7 receptors(USD/well) (**a**) (**c**) |
|---|---|---|---|
| cAMP | Simple to perform.<br>High sensitivity<br>Non-lytic methods available for real-time measurements(*90*) | Gi/o or Gs coupled receptors or chimeric g-proteins only. | PerkinElmer:<br>Alphascreen - 1.79$<br>LANCE - 3.21$<br>Cisbio:<br>HTRF® - 1.97$<br>Eurofins:<br>HitHunter™ - 6.73$ |
| IP1 | More robust alternative to Ca2+ mobilisation assays(*91*) | Gq or Gi coupled receptors or chimeric g-proteins only | Cisbio:<br>HTRF-IP-One® - 2.84$ |
| Ca2+ mobilization | Cheap compared to most other mammalian cell line based assays. | Rapid and transient signal not suitable for slow binding.<br>Gq or Gi coupled receptors or chimeric g-proteins only(*92*) | Eurofins:<br>ChemiBrite™ - 1.80$ |
| $\beta$-arrestin recruitment | Can be used to study biased signalling<br>G-protein independent - very applicable to deorphanization<br>High sensitivity, broad dynamic range.(*93*) | Does not determine Ga-mediated signaling.<br>Most versions of this assay require receptor editing.(*93*) | Eurofins:<br>PathHunter® -11.7$<br>ThermoFischer:<br>Tango™ - 8.58$ |
| Receptor internalisation | Studies biased signaling<br>G-protein independent - very applicable to deorphanization(*94*) | Does not determine Ga-mediated signaling. | Eurofins:<br>PathHunter® -11.4$ |
| GTP$\gamma$[35S] /Eu binding. | Works with any Ga coupling.<br>Fast and simple protocol.(*95*) | Sensitivity lower in HTS format.<br>May require radioactive substances(*95*) | PerkinElmer - 5.29$ |
| Radioligand binding | High sensitivity, broad dynamic range.<br>Fast and simple protocol(*96*)(*97*). | GPCR must have a known ligand.<br>May require radioactive substances<br>Can not determine receptor signalling. | PerkinElmer:<br>Radioligand competition - 5.89$<br>Cisbio:<br>HTRF Tag-lite - 2.96$ |
| Multiplexed screening | Very high throughput<br>low ligand requirements | Requires biased signalling in mammalian cell assays(*98*).<br>Requires Ga-signalling in yeast cell assays(*18*). | PRESTO-Salsa-17$(**d**)(*98*) |
| Yeast biosensor | High dynamic range.<br>Fast and simple protocol.<br>Significantly cheaper than mammalian cell lines | GPCR must be functionally signalling in yeast. | Our work -1.01$ **b** |

**Table S6**: **overview of EC50, Hill coefficient and detection limits observed compared to previously reported assays.** Lowest EC50's reported for each hGPCR for the same ligand as was used in this study. **a.** The only reported data in ChEMBL on ADRA2B against epinephrine is from a radioligand competition assay, and as such the value reported is a $K_i$ corresponding to competition with [3H]Rauwolscine.

| hGPCR | EC50 [M] (This study) | Detection limit [M] (This study) | Hill Coefficient (This study) | Lowest EC50 [M] (Reported in ChEMBL) | Assay type (Reported in ChEMBL) |
|---|---|---|---|---|---|
| ADRA2A | 2.1E-06 | 1.0E-08 | 1.80 | 3.0E-08 (*99*) | [35S]GTP$\gamma$S |
| ADRA2B | 6.9E-07 | 1.0E-07 | 2.20 | 4.0E-7 (*12*) | Radioligand competition[a] |
| HTR4 | 1.6E-06 | 1.0E-10 | 0.29 | 1.2E-08 (*100*) | cAMP |
| HTR1A | 1.4E-05 | 1.0E-06 | 1.20 | 5.0E-10 (*101*) | [35S]GTP$\gamma$S |
| CHRM3 | 8.0E-08 | 1.0E-09 | 1.40 | 5.6E-08 (*27*) | cAMP |
| MTNR1A | 2.5E-08 | 1.0E-11 | 0.33 | 2.2E-11 (*102*) | cAMP |
| OPRM1 | >1.0E-05 | 1.0E-07 | ND | 3.0E-10 (*103*) | cAMP |

Table S7: Genes used in this study

| Gene name | accesion | Codon opti-mised? | Sequence |
|---|---|---|---|
| HTR4 | Q13639-1 | Yes | ATGGATAAATTGGATGCTAATGTTTCTTCTGAAGAAGGTTTTGGTTCTGTTGAAAAAG<br>TTGTTTTGTTGACTTTTTTGTCTACTGTTATTTTGATGGCTATTTTGGGTAATTTGTTGG<br>TTATGGTTGCTGTTTGTTGGGATAGACAATTGAGAAAAATTAAAACTAATTATTTTAT<br>TGTTTCTTTGGCTTTTGCTGATTTGTTGGTTTCTGTTTTGGTTATGCCATTTGGTGCTAT<br>TGAATTGGTTCAAGATATTTGGATTTATGGTGAAGTTTTTTGTTTGGTTAGAACTTCTT<br>TGGATGTTTTGTTGACTACTGCTTCTATTTTTCATTTGTGTTGTATTTCTTTGGATAGAT<br>ATTATGCTATTTGTTGTCAACCATTGGTTTATAGAAATAAAATGACTCCATTGAGAAT<br>TGCTTTGATGTTGGGTGGTTGTTGGGTTATTCCAACTTTTATTTCTTTTTTGCCAATTA<br>TGCAAGGTTGGAATAATATTGGTATTATTGATTTGATTGAAAAAAGAAAATTTAATC<br>AAAATTCTAATTCTACTTATTGTGTTTTTATGGTTAATAAACCATATGCTATTACTTGT<br>TCTGTTGTTGCTTTTTATATTCCATTTTTGTTGATGGTTTTGGCTTATTATAGAATTTAT<br>GTTACTGCTAAAGAACATGCTCATCAAATTCAAATGTTGCAAAGAGCTGGTGCTTCTT<br>CTGAATCTAGACCACAATCTGCTGATCAACATTCTACTCATAGAATGAGAACTGAAA<br>CTAAAGCTGCTAAAACTTTGTGTATTATTATGGGTTGTTTTTGTTTGTGTTGGGCTCCA<br>TTTTTTGTTACTAATATTGTTGATCCATTTATTGATTATACTGTTCCAGGTCAAGTTTG<br>GACTGCTTTTTTGTGGTTGGGTTATATTAATTCTGGTTTGAATCCATTTTTGTATGCTT<br>TTTTGAATAAATCTTTTAGAAGAGCTTTTTTGATTATTTTGTGTTGTGATGATGAAAG<br>ATATAGAAGACCATCTATTTTGGGTCAAACTGTTCCATGTTCTACTACTACTATTAAT<br>GGTTCTACTCATGTTTTGAGAGATGCTGTTGAATGTGGTGGTCAATGGGAATCTCAAT<br>GTCATCCACCAGCTACTTCTCCATTGGTTGCTGCTCAACCATCTGATACTTGA |
| HTR1A | P08908 | Yes | ATGGACGTATTATCACCTGGACAGGGCAACAACACAACAAGTCCCCCTGCACCTTTC<br>GAGACAGGCGGGAACACAACAGGCATCAGTGACGTGACAGTATCATACCAGGTGAT<br>CACATCCTTACTACTTGGGACATTAATATTCTGCGCCGTCCTGGGAAACGCATGCGTA<br>GTGGCAGCCATAGCCCTGGAGAGGAGTCTGCAGAACGTCGCAAACTACCTGATCGGC<br>AGCTTAGCCGTAACAGACTTAATGGTCTCAGTGTTAGTGCTTCCGATGGCCGCATTAT<br>ACCAGGTGCTAAACAAGTGGACGCTGGGACAGGTGACATGCGACCTTTTCATCGCGT<br>TAGACGTCTTATGCTGCACATCATCGATCCTTCACCTTTGCGCAATCGCCCTTGACCG<br>TTACTGGGCCATCACCGACCCCATCGACTACGTGAACAAGAGGACCCCTAGGCGTGC<br>GGCCGCATTAATATCGCTGACCTGGTTAATCGGATTCCTAATATCCATACCGCCTATG<br>TTAGGATGGAGAACCCCGGAAGACCGTAGCGACCCCGACGCCTGCACGATAAGCAA<br>GGACCACGGATACACCATCTACAGTACATTCGGAGCATTCTACATCCCTTTATTACTA<br>ATGTTAGTACTTTACGGCCGTATATTCCGTGCCGCAAGGTTCAGGATCCGTAAGACA<br>GTAAAGAAGGTCGAGAAGACCGGAGCAGACACAAGACACGGGGCCAGTCCTGCTCC<br>GCAGCCAAAGAAGAGTGTGAACGGCGAGAGTGGGTCGAGGAATTGGAGACTGGGCG<br>TCGAGTCGAAGGCCGGCGGAGCATTATGTGCTAACGGGGCGGTCAGGCAAGGTGAC<br>GACGGCGCAGCGTTAGAGGTAATAGAGGTCCACAGGGTCGGAAACAGCAAGGAGCA<br>CTTACCTTTACCTTCTGAGGCCGGACCTACCCCGTGCGCGCCTGCCAGCTTCGAGAGG<br>AAGAACGAGAGGAACGCAGAGGCGAAGCGTAAGATGGCCTTAGCAAGGGAGCGTA<br>AGACAGTGAAGACACTAGGCATCATAATGGGAACCTTCATACTATGCTGGTTACCTT<br>TCTTCATAGTCGCCCTTGTCCTTCCCTTCTGCGAGTCGTCCTGCCACATGCCGACATTA<br>CTTGGGGCAATCATCAACTGGCTTGGCTACAGTAACTCCCTTCTAAACCCCGTGATCT<br>ACGCCTACTTCAACAAGGACTTCCAGAACGCCTTCAAGAAGATAATCAAGTGCAAGT<br>TCTGCCGTCAGTAG |

Continued on next page

Table S7: Genes used in this study (Continued)

| | | | |
|---|---|---|---|
| ADRA2A | P08913 | Yes | ATGTTCCGGCAGGAGCAGCCACTCGCCGAGGGTTCCTTTGCCCCCATGGGAAGTCTT<br>CAGCCTGACGCCGGCAACGCTAGCTGGAATGGCACCGAAGCGCCCGGTGGAGGGGC<br>TCGGGCTACCCCATATTCTCTCCAGGTCACACTGACCCTGGTTTGTCTGGCCGGTCTC<br>CTGATGCTGCTGACGGTCTTTGGCAATGTTTTGGTGATAATTGCGGTGTTCACCTCCA<br>GAGCACTCAAGGCTCCCCAGAACCTGTTCTTGGTCAGCCTGGCGTCTGCCGATATTCT<br>GGTCGCCACCCTCGTAATTCCGTTTTCTCTTGCCAATGAGGTTATGGGTTACTGGTAT<br>TTTGGTAAGGCATGGTGCGAGATATACCTTGCTCTCGACGTTCTCTTCTGCACCAGCA<br>GTATTGTGCATCTGTGCGCAATTAGTCTGGACAGATATTGGTCCATCACGCAGGCAA<br>TAGAATATAACTTGAAAAGGACTCCTCGGCGCATTAAGGCTATCATCATCACTGTCT<br>GGGTCATTAGCGCTGTGATAAGCTTCCCTCCACTTATTTCCATTGAAAAAAAAGGAG<br>GAGGCGGAGGGCCGCAGCCTGCGGAGCCCCGCTGTGAGATCAACGATCAGAAATGG<br>TATGTCATCTCCTCCTGCATTGGGTCCTTCTTCGCCCCATGTCTGATTATGATCCTCGT<br>ATATGTTAGGATTTATCAGATAGCTAAGCGAAGGACCCGGGTCCCCCCATCCAGGAG<br>AGGTCCAGACGCAGTGGCCGCTCCCCCCGGAGGCACAGAAAGGAGACCTAACGGTC<br>TGGGACCAGAGCGCTCCGCAGGACCTGGAGGTGCAGAAGCCGAGCCCTTGCCGACA<br>CAACTGAACGGAGCTCCCGGCGAACCAGCCCCAGCAGGTCCTCGCGATACTGACGCA<br>CTGGACCTCGAAGAGTCCTCTAGCAGCGATCATGCAGAGCGCCCTCCAGGGCCTAGG<br>AGACCTGAAAGAGGACCCAGAGGTAAAGGCAAGGCCCGGGCAAGTCAAGTGAAACC<br>GGGAGACTCTCTGCCTCGGAGGGGTCCAGGTGCTACTGGTATAGGGACTCCTGCCGC<br>CGGTCCCGGAGAAGAGCGGGTCGGCGCCGCTAAGGCAAGTCGCTGGCGGGGCAGAC<br>AAAACCGGGAAAAAAGGTTTACATTCGTGCTCGCGGTAGTGATTGGCGTCTTCGTCG<br>TGTGCTGGTTTCCATTCTTTTTACCTATACCCTGACAGCCGTCGGCTGTTCTGTTCCA<br>CGGACACTGTTCAAGTTCTTTTTCTGGTTTGGCTACTGTAACTCAAGCCTGAACCCGG<br>TAATCTATACAATATTTAACCACGATTTTCGGAGGGCTTTTAAAAAGATCCTGTGCAG<br>AGGCGATAGGAAACGAATCGTTTAA |
| ADRA2B | P18089 | Yes | ATGGATCATCAGGATCCCTATTCTGTCCAGGCCACTGCGGCCATCGCTGCCGCTATCA<br>CTTTTCTCATTCTCTTTACCATCTTCGGGAACGCGCTTGTAATTCTGGCGGTGCTCACC<br>AGCAGGTCACTCAGGGCACCACAAAACCTTTTTCTGGTCTCCCTCGCGGCAGCTGAC<br>ATTCTCGTCGCAACCCTCATCATTCCCTTTTCACTCGCTAATGAGTTGTTGGGATATTG<br>GTATTTTCGCCGAACCTGGTGCGAGGTGTATCTCGCCCTGGACGTACTGTTTTGCACC<br>TCTTCAATTGTCCATCTTTGTGCAATCAGCCTGGACAGATATTGGGCCGTGTCAAGGG<br>CTCTGGAGTATAATAGCAAGCGGACACCTAGGAGGATCAAGTGCATAATCCTTACAG<br>TCTGGCTCATAGCGGCAGTAATAAGCCTGCCCCCCCTTATATACAAAGGAGATCAGG<br>GCCCTCAACCCAGAGGGCGGCCACAATGCAAGCTGAATCAAGAGGCTTGGTACATTT<br>TGGCGAGCAGCATCGGAAGCTTCTTTGCCCCATGCCTGATCATGATTTTGGTATACCT<br>CAGGATTTACCTGATCGCTAAGCGGTCTAATAGACGCGGGCCTCGGGCTAAGGGAGG<br>ACCAGGCCAGGGGGAATCCAAGCAGCCTAGGCCTGATCACGGCGGCGCACTTGCCA<br>GTGCCAAGCTGCCCGCACTGGCCTCCGTCGCGTCAGCAAGAGAAGTCAATGGCCACA<br>GTAAATCTACTGGTGAAAAGGAAGAGGGCGAGACCCCTGAAGATACCGGAACCCGC<br>GCACTGCCTCCAAGTTGGGCTGCGCTGCCAAACTCTGGGCAGGGACAGAAGGAGGG<br>GGTGTGCGGCGCAAGTCCCGAGGACGAGGCCGAGGAGGAGGAAGAAGAAGAGGAG<br>GAAGAGGAGGAATGCGAGCCTCAAGCAGTGCCTGTAAGTCCAGCCTCTGCATGCTCA<br>CCTCCTCTTCAGCAGCCACAGGGCTCCAGAGTCCTCGCTACCCTCAGAGGGCAAGTG<br>CTGCTCGGAAGGGGTGTCGGTGCCATTGGAGGGCAATGGTGGCGCCGGCGGGCACA<br>GCTGACCAGAGAAAAGCGCTTCACCTTTGTGCTCGCCGTCGTGATCGGCGTATTCGTT<br>CTCTGCTGGTTCCCATTTTTCTTCTCCTACTCCTTGGGCGCGATTTGTCCTAAGCATTG<br>CAAAGTACCACATGGGCTCTTTCAATTTTTTTTCTGGATTGGTTATTGCAATAGCTCA<br>CTGAATCCTGTGATTTACACCATTTTCAATCAGGATTTCCGGCGGGCCTTTAGACGCA<br>TTCTGTGCAGGCCGTGGACCCAGACTGCATGGTAA |
| CHRM3<br>Δil3 | P20309 | Yes | ATGACGCTGCATAATAATTCAACCACAAGCCCTCTGTTCCCTAACATCAGTAGTAGCT<br>GGATACACTCTCCTAGCGACGCCGGACTTCCTCCCGGAACCGTGACCCACTTTGGGT<br>CATATAACGTCTCTCGGGCCGCAGGCAATTTTTCTTCCCCAGATGGGACAACCGACG<br>ATCCGCTCGGCGGCCACACTGTATGGCAGGTGGTTTTCATAGCCTTTCTGACCGGCAT<br>TCTGGCTCTGGTCACAATCATCGGCAATATTCTGGTAATTGTGTCCTTCAAGGTAAAC<br>AAACAACTGAAGACTGTCAATAACTACTTTCTGCTTAGTTTGGCATGCGCCGATCTGA<br>TTATCGGAGTGATAAGTATGAATCTCTTCACTACTTACATCATTATGAACAGATGGGC<br>TCTGGGTAACTTGGCTTGCGACCTGTGGCTCGCAATTGACTATGTCGCAAGCAACGC<br>CAGTGTGATGAACCTTCTGGTTATTAGCTTCGACCGCTATTTTAGCATTACACGCCCT<br>CTGACCTACAGAGCGAAACGCACTACAAAGCGGGCAGGAGTTATGATCGGCCTGGC<br>CTGGGTCATTTCCTTCGTGCTCTGGGCACCAGCCATCTTGTTCTGGCAGTATTTCGTC<br>GGTAAGAGGACAGTCCCACCAGGCGAGTGCTTCATCCAATTTTTGTCCGAGCCAACC<br>ATAACATTCGGCACAGCCATCGCGGCATTTTACATGCCAGTGACTATCATGACCATTC<br>TGTACTGGAGAATCTACAAAGAAACCGAAAAGCGGACAAAGGAGTTGGCAGGGCTT<br>CAGGCATCAGGCACAGAGACCCGGTCACAGATAACCAAGCGCAAGCGAATGTCACT<br>CGTGAAAGAAAAAAAAGCAGCACAGACATTGAGCGCCATCTTGCTCGCTTTTATCAT<br>AACATGGACACCCTACAATATCATGGTCCTCGTGAACACTTTCTGCGACTCCTGTATT<br>CCTAAGACTTTCTGGAACCTCGGCTACTGGCTGTGCTATATCAACAGTACAGTTAACC<br>CTGTGTGCTACGCTCTCTGTAACAAGACCTTTAGGACCACGTTCAAGATGCTCCTCCT<br>GTGCCAGTGTGACAAAAAGAAACGACGAAAGCAGCAATATCAGCAGCGACAGAGTG<br>TGATCTTTCATAAACGGGCGCCAGAACAAGCGCTCTAA |



Continued on next page

Table S7: Genes used in this study (Continued)

| | | | |
|---|---|---|---|
| OPRM1 | P35372-1 | No | ATGGACAGCAGCGCTGCCCCCACGAACGCCAGCAATTGCACTGATGCCTTGGCGTAC<br>TCAAGTTGCTCCCCAGCACCCAGCCCCGGTTCCTGGGTCAACTTGTCCCACTTAGATG<br>GCAACCTGTCCGACCCATGCGGTCCGAACCGCACCGACCTGGGCGGGAGAGACAGC<br>CTGTGCCCTCCGACCGGCAGTCCCTCCATGATCACGGCCATCACGATCATGGCCCTCT<br>ACTCCATCGTGTGCGTGGTGGGGCTCTTCGGAAACTTCCTGGTCATGTATGTGATTGT<br>CAGATACACCAAGATGAAGACTGCCACCAACATCTACATTTTCAACCTTGCTCTGGC<br>AGATGCCTTAGCCACCAGTACCCTGCCCTTCCAGAGTGTGAATTACCTAATGGGAAC<br>ATGGCCATTTGGAACCATCCTTTGCAAGATAGTGATCTCCATAGATTACTATAACATG<br>TTCACCAGCATATTCACCCTCTGCACCATGAGTGTTGATCGATACATTGCAGTCTGCC<br>ACCCTGTCAAGGCCTTAGATTTCCGTACTCCCCGAAATGCCAAAATTATCAATGTCTG<br>CAACTGGATTCTCTCTTCAGCCATTGGTCTTCCTGTAATGTTCATGGCTACAACAAAA<br>TACAGGCAAGGTTCCATAGATTGTACACTAACATTCTCTCATCCAACCTGGTACTGGG<br>AAAACCTGCTGAAGATCTGTGTTTTCATCTTCGCCTTCATTATGCCAGTGCTCATCATT<br>ACCGTGTGCTATGGACTGATGATCTTGCGCCTCAAGAGTGTCCGCATGCTCTCTGGCT<br>CCAAAGAAAAGGACAGGAATCTTCGAAGGATCACCAGGATGGTGCTGGTGGTGGTG<br>GCTGTGTTCATCGTCTGCTGGACTCCCATTCACATTTACGTCATCATTAAAGCCTTGG<br>TTACAATCCCAGAAACTACGTTCCAGACTGTTTCTTGGCACTTCTGCATTGCTCTAGG<br>TTACACAAACAGCTGCCTCAACCCAGTCCTTTATGCATTTCTGGATGAAAACTTCAAA<br>CGATGCTTCAGAGAGTTCTGTATCCCAACCTCTTCCAACATTGAGCAACAAAACTCC<br>ACTCGAATTCGTCAGAACACTAGAGACCACCCCTCCACGGCCAATACAGTGGATAGA<br>ACTAATCATCAGCTAGAAAATCTGGAAGCAGAAACTGCTCCGTTGCCCTAA |
| MTNR1A | P48039 | Yes | ATGCAAGGTAATGGTTCTGCTTTGCCAAATGCTTCTCAACCAGTTTTGAGAGGTGATG<br>GTGCTAGACCTTCTTGGTTGGCTTCTGCTTTAGCTTGTGTTTTGATTTTCACCATCGTT<br>GTCGATATCTTGGGTAACTTGTTGGTTATCTTGTCCGTCTACCGTAACAAGAAATTGA<br>GAAACGCTGGTAACATCTTCGTTGTTTCTTTGGCTGTTGCTGATTTGGTTGTTGCTATC<br>TATCCATATCCACTGGTCTTGATGTCCATTTTTAACAACGGTTGGAACTTGGGTTACT<br>TGCATTGTCAAGTTTCTGGTTTCTTGATGGGTTTGTCCGTTATTGGTTCCATTTTCAAC<br>ATTACCGGTATCGCCATTAACAGGTACTGTTACATTTGCCACTCACTGAAGTACGACA<br>AGTTGTACTCTTCTAAGAACTCCTTGTGCTACGTTTTGTTGATCTGGTTGTTAACTTTG<br>GCTGCTGTTTTGCCTAATTTGAGAGCTGGTACATTGCAATACGATCCAAGAATCTACT<br>CTTGTACCTTCGCTCAATCTGTTTCTTCTGCTTACACTATTGCCGTTGTCGTTTTCCATT<br>TTTTGGTCCCAATGATTATCGTCATCTTCTGCTACTTGAGAATCTGGATTTTGGTCTTG<br>CAAGTCAGACAAAGAGTTAAGCCAGATAGAAAGCCAAAATTGAAGCCACAAGACTT<br>CAGAAACTTCGTTACCATGTTTGTGGTTTTCGTTTTGTTCGCTATTTGTTGGGCTCCAT<br>TGAACTTTATTGGTTTGGCAGTTGCTTCTGATCCAGCTTCTATGGTTCCAAGAATTCC<br>AGAATGGTTGTTCGTTGCTTCTTACTACATGGCTTACTTCAACTCTTGTTTGAACGCCA<br>TTATCTACGGCTTGTTGAATCAGAACTTTAGGAAAGAGTACAGGCGTATCATCGTTTC<br>TTTGTGTACTGCTAGAGTTTTCTTCGTCGATTCCTCTAATGATGTTGCCGATAGAGTTA<br>AGTGGAAACCATCTCCATTGATGACCAACAACAATGTTGTCAAGGTTGACTCCGTTG<br>GATCCTGA |
| NanoLuc | (*26*) | No | ATGGTCTTCACACTCGAAGATTTCGTTGGGGACTGGCGACAGACAGCCGGCTACAAC<br>CTGGACCAAGTCCTTGAACAGGGAGGTGTGTCCAGTTTGTTTCAGAATCTCGGGGTG<br>TCCGTAACTCCGATCCAAAGGATTGTCCTGAGCGGTGAAAATGGGCTGAAGATCGAC<br>ATCCATGTCATCATCCCGTATGAAGGTCTGAGCGGCGACCAAATGGGCCAGATCGAA<br>AAAATTTTTAAGGTGGTGTACCCTGTGGATGATCATCACTTTAAGGTGATCCTGCACT<br>ATGGCACACTGGTAATCGACGGGGTTACGCCGAACATGATCGACTATTTCGGACGGC<br>CGTATGAAGGCATCGCCGTGTTCGACGGCAAAAAGATCACTGTAACAGGGACCCTGT<br>GGAACGGCAACAAAATTATCGACGAGCGCCTGATCAACCCCGACGGCTCCCTGCTGT<br>TCCGAGTAACCATCAACGGAGTGACCGGCTGGCGGCTGTGCGAACGCATTCTGGCGT<br>AA |

**Table S8**: Yeast strains utilised in this study, ScFH236 and 237 are based on design 4 presented in Shaw et al 2019 (*25*)

| | | |
|---|---|---|
| ScFH236 | BY4741-MATa,sst2Δ0,far1Δ0,bar1Δ0,ste2Δ0,ste12Δ0,gpa1Δ0,ste3Δ0,mf(alpha)1Δ0,mf(alpha)2Δ0,mfa1Δ0,mfa2Δ0,gpr1Δ0,gpa2Δ0,LexO(6x)-pLEU2m-NanoLuc-tTDH1-pCCW12-STE2-tSSA1-pPGK1-GPA1-YIGLC-tENO2-pRAD27-LexA-PRD-tENO1-URA3 | This study |
| ScFH237 | BY4741-MATa,sst2Δ0,far1Δ0,bar1Δ0,ste2Δ0,ste12Δ0,gpa1Δ0,ste3Δ0,mf(alpha)1Δ0,mf(alpha)2Δ0,mfa1Δ0,mfa2Δ0,gpr1Δ0,gpa2Δ0,LexO(6x)-pLEU2m-NanoLuc-tTDH1-pCCW12-STE2-tSSA1-pPGK1-GPA1-YIGLC-tENO2-pRAD27-LexA-PRD-tENO1-URA3 | This study |
| ScS1 | ScFH237 + PCCW12-HsHTR1A-TCYC1 | This study |
| ScS2 | ScFH237 + PCCW12-HsHTR4-TCYC1 | This study |
| ScS3 | ScFH237 + PCCW12-HsADRA2A-TCYC1 | This study |
| ScS4 | ScFH236 + PCCW12-HsADRA2B-TCYC1 | This study |
| ScS5 | ScFH237 + PCCW12-OPRM1-TCYC1 | This study |
| ScS6 | ScFH237 + PCCW12-HsCHRM3-TCYC1 | This study |
| ScS7 | ScFH237 + PCCW12-HsMTNR1A-TCYC1 | This study |

**Table S9**: **Selected Subset of pdCSM Dataset**. hGPCRs were selected which also occurred in the set of 128 hGPCRs investigated in this paper. Number of training (47013) and test (7114) data points are summarised.

| Target hGPCR | Test Entries | Train Entries |
|---|---|---|
| 5HTR2C (P28335) | 353 | 2765 |
| 5HTR6 (P50406) | 345 | 2699 |
| ADORA1A (P30542) | 424 | 3409 |
| ADORA2B (P29275) | 274 | 1835 |
| ADORA3 (P0DMS8) | 413 | 3100 |
| ADRA1A (P35348) | 253 | 1645 |
| BDKRB1 (P46663) | 148 | 608 |
| CCR3 (P51677) | 184 | 947 |
| CHRM3 (P20309) | 310 | 1698 |
| CHRM4 (P08173) | 141 | 837 |
| CHRM5 (P08912) | 139 | 820 |
| DRD4 (P21917) | 276 | 2059 |
| EDNRB (P24530) | 172 | 815 |
| GNRHR (P30968) | 276 | 1097 |
| GPBAR1 (Q8TDU6) | 71 | 372 |
| GPR35 (Q9HC97) | 72 | 408 |
| GPR44 (Q9Y5Y4) | 342 | 2407 |
| HCAR2 (Q8TDS4) | 70 | 434 |
| HRH3 (Q9Y5N1) | 464 | 3133 |
| HTR1A (P08908) | 420 | 3370 |
| MCHR1 (Q99705) | 435 | 3286 |
| MTNR1A (P48039) | 152 | 891 |
| NK2R (P21452) | 160 | 762 |
| OPRM1 (P35372) | 624 | 4651 |
| P2RY1 (P47900) | 107 | 461 |
| PTGER1 (P34995) | 101 | 640 |
| S1PR3 (Q99500) | 149 | 939 |
| S1PR5 (Q9H228) | 68 | 349 |
| SSTR5 (P35346) | 171 | 576 |

**Table S10**: **Addressing similarities and differences of CSNN with published methods**. We highlight that most published methods are compound-to-prediction architectures (which answer the query P(y|c,t)), while CSNNs are distinctly different due to their neighbourhood-to-prediction architecture (P(y|c,t,$\mathcal{N}$(c))).

| **Article** | **Prediction Objective** | **Model Architecture** | **Difference** | **Similarity** | **Notes** |
|---|---|---|---|---|---|
| Lim et al. J. Chem. Inf. Model. 2019, 59, 9, 3981–3988 | Classification (bind/not bind) P(y\|t,c) | GAT | Require explicit 3D binding poses for training. Input for inference is an explicit 3D pose. Works only on a single ligand-target pose during inference. | Use GAT architecture. | Distantly related. Not directly comparable. |
| Jiang et al., RSC Adv, 2020, 10:20701-20712 | Regression (DTA), P(y\|t,c) | GCN and GAT | Target is represented as a contact map, which carries far less information than the ESM embeddings. They predict drug binding affinity. Not directly developed for GPCRs. They operate only on a single drug-target pair during inference. | Addresses drug-target affinity (DTA), which our extended experiments also address. Graph neural network usage. | Conceptually dissimilar, does not use chemical neighborhoods, chemical space networks, or message passing at this level. |
| Bguyen et al., Bioinformatics, 2021, 37(8): 1140-1147 | Regression (DTA), P(y\|t,c) | GNNs (GIN, GAT, GCN) | We kindly stress again that they operate only on a single drug-target pair during inference. Uses one-hot encoding of sequence, which is a very sparse representation. | GIN, GAT, GNN usage. Ibid. as above with DTA. | Method cannot integrate available data dynamically during inference |
| Sun et al., Bioinformatics, 2024, 40(3): btae135 | Classification (bind/not bind) P(y\|t,c) | GNN (to integrate information from concatenated pre-trained representations) | Uses handcrafted descriptors/featurization (which are always outperformed by learned representations), but they operate only on a single drug-target pair during inference. | Adresses DTI and uses GNNs. ESM is used. Pre-trained Chemformer is used for a learned representation. | Method cannot integrate available data dynamically during inference. |
| Velloso et al. Bioinformatics Advances, Volume 1, Issue 1, 2021, vbab031 | Regression (DTA), P(y\|c) (one model per target) | RF, Gradient Boosting. | Use handcrafted descriptors and simple ML models (XGboost, RF). Again, we kindly point out that they operate only on a single drug-target pair during inference. The repeated occurrence of this seems to indicate that a misunderstanding of the paper has occurred. | GPCR focus, DTA. They, however, build one model per target rather than a single model (their method doesn't enable this either without the addition of extremely sparse OHE or ESM-2 embeddings). | We have benchmarked against their data, as it is well documented and has available test-train splits. |
| Kanai et al., Molecules. 2021 ;26(17):5131 | Classification (bind/not bind) P(y\|t,c) | SVM | The tensor product of compound and protein is an elegant approach to model their joint space. However, they operate only on a single drug-target pair during inference. Although this paper is the most relevant suggestion so far. | Uses a similarity metric, with a kernel, to make binary activity predictions with a SVM. This is partly related as they implicitly model neighbourhoods in the tensor product space with a decision boundary. | Interesting use of kernel based method to look at similarities between compounds and GPCR targets. |

**Table S10**: **Addressing similarities and differences of CSNN with published methods**. We highlight that most published methods are compound-to-prediction architectures (which answer the query P(y|c,t)), while CSNNs are distinctly different due to their neighbourhood-to-prediction architecture (P(y|c,t,$\mathcal{N}$(c))).

| **Article** | **Prediction Objective** | **Model Architecture** | **Difference** | **Similarity** | **Notes** |
|---|---|---|---|---|---|
| Cai et al. J Chem Inf Model. 2021 Apr 26;61(4):1570-1582 | P(y\|t,c), binary classification (active, inactive). | Distilled sequence alignment embedding (DISAE, fine-tuned) and pretrained MPNN for compound. | Use only a single compound-target pair during training and inference. | GPCR focus and classification objective. | |
| Lu et al. IEEE J Biomed Health Inform. 2023 doi: 10.1109/JBHI.2023.3307928 | Both classification and regression P(y\|t,c) | CNN | We stress again the fact that they operate only on a single drug-target pair during inference. Fig 5. Shows that in fact their correlations are neither strong nor good predictions. Model shows strong 'mode seeking' behavior, which our benchmark on DTA does not. Perhaps a point which applies to many of the aforementioned papers is also the lack of experimental validation. | GPCR focus. | |
| Ahmed et al. Sci Rep 11, 9510 (2021) | Classification (bind/not bind) P(y\|t,c) | RF/ XGBoost / SVMs | They operate only on a single drug-target pair during inference. No experimental verification. Use simple algorithmic descriptors from RDkit, which since the paper in 2019 have strong support for always being outperformed by learned representations: doi.org/10.1021/acs.jcim.9b00237 | GPCR focus. We in fact use their hyperparameter searches to guide ours, thus allowing for a comparison with published DTI approaches. As we show in Figure 3C, RF/MLP methods performs markedly worse in comparison to CSNN or argmax(). | The GPCR_LigandClassify.py paper. Their test/train data is not available nor reproducible. |
| Zhang et al. Briefings in Bioinformatics, Volume 25, Issue 4, July 2024, bbae281 | Both classification and regression P(y\|t,c) | GNN | We point out again that they operate only on a single drug-target pair during inference. Single model per target, no way of doing one model for all targets. AUC discriminatory values are ≈ 0.72 (0.48-0.93) (AUC = 0.5). Average pearson correlation is 0.39, again showing far lower performance. | GPCR focus. Perform experimental verification. | |
| Cai T, et al.. Bioinformatics ;38(9):2561-2570 | Classification (bind/not bind) P(y\|t,c) and bioactivity class (antagonist / agonist). | Unclear (Use of several pretrained models). | Out of distribution testing, which is quite elegant. OOD for proteins split and chemical similarity split. We kindly stress the fact that the query (P(y\|t,c)), is unlike CSNN. The method operates only on a single drug-target pair during inference. Fig 5. Training curves indicate significant difficulty in training (high variance) and the OOD training curves barely show any changes in the precision (nothing learned by model). No experimental validation. | Data sources are quite similar. | |
| Huang et al. J Cheminform 16, 10 (2024). doi.org/10.1186/s13321-024-00806-3 | P(y\|t,c) in the Regression setting (EC50) | CatBoost, RF, and LightGBM | Uses rule-based embedding (for both sequence and compound) and concatenate them, thus only allow for prediction of a single drug-target interaction, without using available data during inference. This is a compound-to-prediction architecture unlike CSNN. | GPCR focus. Similar data-type. | Does not deal with using available data during inference. |
